# Synteny-based genome assembly for 16 species of *Heliconius* butterflies, and an assessment of structural variation across the genus

**DOI:** 10.1101/2020.10.29.359505

**Authors:** Fernando A. Seixas, Nathaniel B. Edelman, James Mallet

## Abstract

*Heliconius* butterflies (Lepidoptera: Nymphalidae) are a group of 48 neotropical species widely studied in evolutionary research. Despite the wealth of genomic data generated in past years, chromosomal level genome assemblies currently exist for only two species, *Heliconius melpomene* and *H. erato*, each a representative of one of the two major clades of the genus. Here, we use these reference genomes to improve the contiguity of previously published draft genome assemblies of 16 *Heliconius* species. Using a reference-assisted scaffolding approach, we place and order the scaffolds of these genomes onto chromosomes, resulting in 95.7-99.9% of their genomes anchored to chromosomes. Genome sizes are somewhat variable among species (270-422 Mb) and in one small group of species (*H. hecale*, *H. elevatus* and *H. pardalinus*) differences in genome size are mainly driven by a few restricted repetitive regions. Genes within these repeat regions show an increase in exon copy number, an absence of internal stop codons, evidence of constraint on non-synonymous changes, and increased expression, all of which suggest that the extra copies are functional. Finally, we conducted a systematic search for inversions and identified five moderately large inversions fixed between the two major *Heliconius* clades. We infer that one of these inversions was transferred by introgression between the lineages leading to the *erato*/*sara* and *burneyi*/*doris* clades. These reference-guided assemblies represent a major improvement in *Heliconius* genomic resources that should aid further genetic and evolutionary studies in this genus.

## Introduction

Advances in sequencing technology have revolutionized the field of evolutionary biology. Generating short-read genomic datasets is now common practice, enabling investigation of fundamental evolutionary processes including the genetic basis of adaptive traits, dynamics of selection on particular alleles, and demographic histories of populations. In order to exploit the power of low-cost short-read data, one must usually align reads to a reference genome.

The availability of high-quality reference genomes can determine the breadth and power of comparative and population genomic analyses in evolutionary studies. For instance, placing genome scaffolds on chromosomes allows one to contrast patterns between autosomes and sex chromosomes which has been important for understanding speciation (Coyne and Orr 1989; Coyne 2018; Prowell 1998; Masly and Presgraves 2007; Fontaine et al. 2015; Ellegren et al. 2012; Seixas et al. 2018; Martin et al. 2019).

Anchoring scaffolds to chromosomes can also enable discovery of divergence and gene flow along chromosomes and how it is modified by recombination rate variation (Schumer et al. 2018; Martin et al. 2019). Furthermore, chromosome-level assemblies have been shown to greatly improve the power and resolution of genome-wide association and QTL studies (Benevenuto et al. 2019; Markelz et al. 2017). However, high-quality, chromosome-level, contiguous reference genome assemblies are often limited to one or a few species in many groups of taxa, especially in non-model organisms. This is partly due to the fact that generating near-complete chromosome-level assemblies normally requires integrating a mixture of high fidelity short-read sequencing data (today typically Illumina), and more costly long-read sequencing data (such as PacBio or Nanopore), genetic linkage mapping, optical (restriction site) mapping, and/or chromatin interaction frequency data (Hi-C) (Rice and Green 2019; Ghurye and Pop 2019; Yang et al. 2020; Wei et al. 2020; Deschamps et al. 2018; Yu et al. 2019). These methods can be prohibitively expensive and time consuming, especially for entire clades.

With 48 described species, *Heliconius* butterflies are a prime example of an adaptive radiation where multiple chromosome-level reference assemblies could improve evolutionary analyses. Currently, published high-contiguity genome assemblies (hereafter, reference genomes) exist for only two species – *H. melpomene* (Davey et al. 2017) and *H. erato* (*H. erato lativitta –* Lewis et al., 2016; *H. erato demophoon* – Van Belleghem et al., 2017). While these chromosome-level reference assemblies are essential tools for genomic studies in *Heliconius*, each has limitations. At 275 Mb, *H. melpomene* has the smallest *Heliconius* genome assembled to date (Edelman et al. 2019). Mapping short-read sequencing data from other species with larger genomes to this reference genome is likely to result both in the loss of information, due to loss of ancestral orthologous sequence in the *H. melpomene* genome, and spurious read mapping to similar but non-orthologous regions. In contrast, the two *H. erato* reference genomes (383 and 418 Mb) are among the largest *Heliconius* genomes assembled to date. However, while these might be appropriate for studies focusing on closely related species (e.g. species within the *erato* clade), mapping accuracy decreases in more divergent species (Prüfer et al. 2010) and better results are obtained when mapping to closer reference genomes (Gopalakrishnan et al. 2017). Also, as we move from comparative (e.g. phylogenomic) towards more functional genetics studies (Lewis et al. 2016; Lewis and Reed 2019; Pinharanda et al. 2019), this genus could benefit greatly from higher-quality species-specific genomic resources.

Recently, *de novo* draft genomes of 16 *Heliconius* species (Supplemental Table S1) have been assembled (Edelman et al. 2019). These genomes were generated from Illumina PCR-free libraries sequenced at deep coverage (at least 60X coverage) using paired-end 250-bp reads on the Illumina Hi-Seq 2500 and assembled using *w2rap* (Clavijo et al. 2017), an extension of the DISCOVAR *de novo* genome assembly method (https://software.broadinstitute.org/software/discovar/blog/; Love et al. 2016; Weisenfeld et al. 2014). This strategy results in high-quality genomes in terms of read accuracy, contiguity within scaffolds, and genome completeness (87.5-97.3% complete single copy core BUSCO genes present; Edelman et al. 2019). Nonetheless, because these assemblies (hereafter, *w2rap* assemblies) used only short-read data, they were considerably more fragmented (scaffold N50 = 23-106 kb) than the *Heliconius* reference genomes. Furthermore, scaffolds were not assigned to chromosomes.

A cost-effective approach for improving the contiguity of existing draft genomes is to use synteny-based methods that identify potentially adjacent scaffolds from multi-species alignments. Such methods are particularly efficient if high-quality reference genome assemblies of closely related species are available, and especially if there is high synteny between the genomes of the draft and reference assemblies (Alonge et al. 2019), as in *Heliconius*. While a limited number of genomic rearrangements have been identified in *Heliconius* (Davey et al. 2017; Jay et al. 2018; Edelman et al. 2019; Meier et al. 2020), even species as divergent as *H. melpomene* and *H. erato*, which last shared a common ancestor over 10 million years ago, remain highly collinear (Davey et al. 2017). Synteny-based assembly should thus be especially effective within this genus.

Here, we exploit the chromosome-mapped assemblies of the *H. melpomene melpomene* and *H. erato demophoon* reference genomes to guide improvement of contiguity of the *w2rap* draft genome assemblies of 16 *Heliconius* species. The *w2rap* scaffolds were ordered, oriented and anchored onto chromosomes, resulting in a level of completeness of the scaffolded *w2rap* assemblies similar to that of reference genomes. A potential weakness of our synteny-based assembly method is that it can miss structural variation among species where it occurs. However, we use these scaffolded *w2rap* assemblies (hereafter, reference-guided assemblies) to identify clade-specific local genomic expansions due to local duplications with potentially functional consequences. To estimate how much structural variation we might be missing, we also carry out a systematic search for candidate inversions in the genus using the original *w2rap* scaffolds to detect break-points, and demonstrate that the results can be used to investigate phylogenetic uncertainty and gene flow deep in the tree of *Heliconius* species.

## Results

### Reference-guided genome assemblies and annotation

Alternative haplotype scaffolds in the *w2rap* assemblies were first merged using HaploMerger2, reducing the numbers of scaffolds by 31.3-64.6% and total assembly length by 3.4-25.9% (Supplemental Table S2). These haplotype-merged scaffolds were then assembled using our reference guided approach. Standard metrics for the resulting assemblies can be found in Supplemental Table S2. Contiguity of all assemblies was considerably improved, with a reduction in the numbers of scaffolds to 0.9-16.8% of the original *w2rap* assemblies (Supplemental Table S2; Supplementary Figs. S1-2). N50 length values were 14.2-20.0 Mb when using the *H. melpomene* genome as reference (the N50 of the *H. melpomene* reference genome is *ca*. 14.3 Mb) and 7.1-11.5 Mb when using the *H. erato demophoon* genome (*H. erato demophoon* reference genome N50 is *ca.* 10.7Mb). In general, scaffolds in the reference anchored to chromosomes have a single corresponding scaffold in each of our reference-guided assemblies (Supplemental Figs. S3-S36). Overall, 94.5-99.5% and 91.6-99.7% of bases in each reference-guided assembly were anchored to chromosomes using the *H. melpomene* and *H. erato* references, respectively (Supplemental Table S2; Figure 1B; Supplemental Fig. S37). For each species, the proportion of reference-guided assembly length anchored to chromosomes was higher in assemblies guided by the genome of the phylogenetically closest species, *H. doris* being the only exception. This species is distant from both reference genomes, but has been inferred to be phylogenetically closer to *H. melpomene* (Edelman et al., 2019; Kozak et al., 2018; Kozak et al., 2015). However, it shows a 0.2% higher proportion of the assembly length included in scaffolds anchored to chromosomes using *H. erato demophoon* as the reference, likely because the larger genome of *H. erato* contains ancestral sequence that was lost by the smaller *H. melpomene* genome but retained in the early branching *H. doris*.

**Figure 1.**
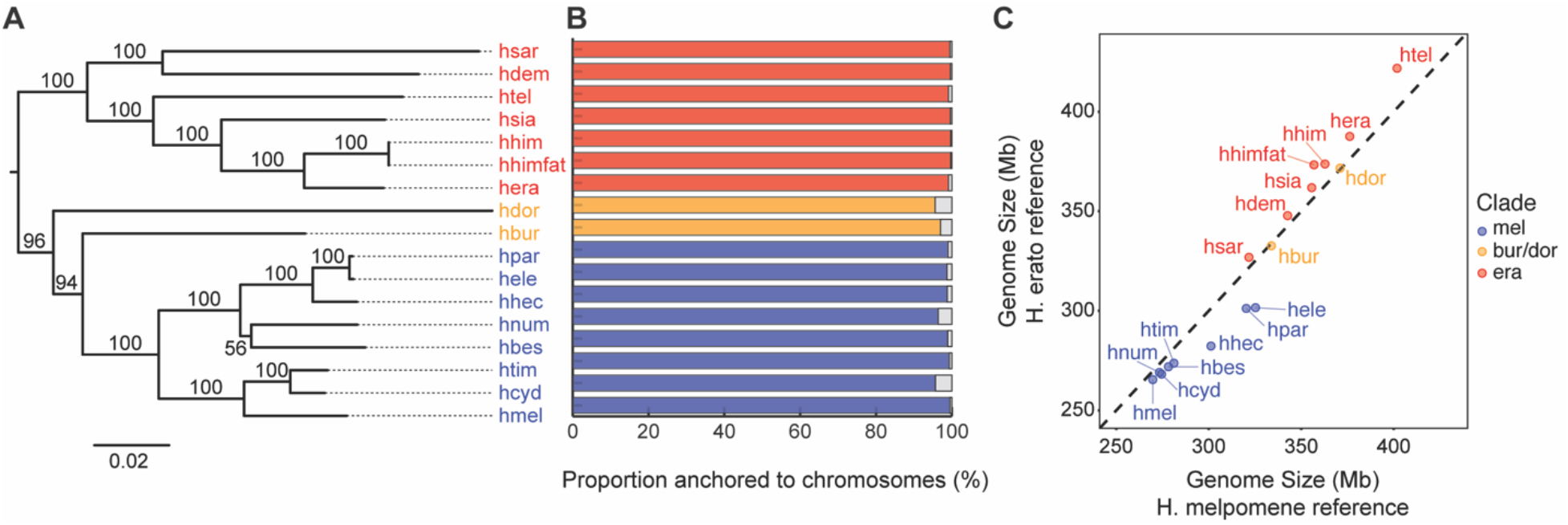
Reference-guided assemblies. **A** – Maximum-likelihood tree from whole mitochondrial genomes assembled here. Bootstrap values are shown next to the branches. The tree was rooted using the *E. tales* mitochondrial genome. **B** – Proportion of the reference-scaffolded assemblies length anchored to chromosomes. The results are shown for the reference-guided assemblies mapped to the closest reference (either *H. melpomene* or *H. erato demophoon*). For a complete report of the results see Supplemental Fig. S37. **C** – Reference-scaffolded assemblies nuclear genome sizes using either *H. melpomene* (x-axis) or *H. erato demophoon* (y-axis) as the reference. The dashed line represents the expectation if there was a 1:1 correspondence. In all panels, subclade memberships are represented by different colours – *melpomene/silvaniform* (blue), *burneyi+doris* (yellow), *erato*/*sara* (red). Species codes for all the new reference-guided assemblies are as follows: hmel – *H. melpomene*; hcyd – *H. cydno*; htim – *H. timareta*; hbes – *H. besckei*; hnum – *H. numata*; hhec - *H. hecale*; hele – *H. elevatus*; hpar – *H. pardalinus*; hbur – *H. burneyi*; hdor – *H. doris*; hera – *H. erato*; hhimfat – *H. himera*; hhim – *H. himera*; hsia – *H. hecalesia*; htel – *H. telesiphe*; hdem – *H. demeter*; hsar – *H. sara*.

Genome sizes, considering only scaffolds anchored to chromosomes, varied between *ca.* 270-422 Mb (Figure 1C; Supplemental Table S2). Phylogeny is a predictor of genome size: species within the *erato*/*sara* clade have larger genomes (327-422 Mb) than species in the *melpomene*/silvaniform group (270-325 Mb; Figure 1C), while genome sizes of *H. burneyi* and *H. doris* (334 and 371 Mb, respectively) are more typical of those of the *erato*/*sara* group. The genome size of the *H. melpomene* reference-guided assembly (270 Mb; 271 Mb including all scaffolds) is similar to that of the reference assembly (273 Mb; 275 Mb total; Davey et al., 2017), but both are smaller than estimates based on flow cytometry (292 Mb +/− 2.4 Mb; Jiggins et al., 2005). The genome size of the *H. erato demophoon* reference-guided assembly (388 Mb; 391 Mb total) is a little larger than that of the reference assembly (383 Mb; Van Belleghem et al. 2017) but both are smaller than flow cytometry estimates (396-397 Mb, misnamed as *H. e. petiverana*; Tobler et al., 2005). Despite the difference in genome sizes of the reference genome used to guide scaffolding, genome sizes of our assemblies (considering only scaffolds anchored to chromosomes) did not depend strongly on which reference genome was used (Spearman’s Rank correlation test ρ= 0. 99; *P* << 0.01; linear regression slope=0.81; Figure 1C). Likewise, individual chromosome lengths of the species assemblies scaffolded using the two different references differed little and were highly correlated (Spearman’s Rank correlation coefficient, ρ= 0.94-0.99; *P* << 0.01; linear regression slope = 0.80-1.05; Supplemental Fig. S38; Supplemental Table S3).

Assembly completeness was evaluated by the presence of core arthropod genes in BUSCO. The proportion of detected orthologs varied between 98.6 and 99.6%, values similar to those reported by Edelman et al. (2019) for the original *w2rap* genomes (Supplemental Fig. S39; Supplemental Table S4). There are however improvements (1-10% increase) in terms of the percentage of complete single copy BUSCOs and a reduction in complete duplicated, fragmented and missing BUSCOS. These improvements are likely a consequence of the increased contiguity and decreased scaffold redundancy (due to the collapsing of alternative haplotype scaffolds) in the reference-guided assemblies, which allows for better mapping of the core genes.

Gene annotation of *H. melpomene* and *H. erato* demophoon reference genomes was mapped onto the reference-guided assemblies using the annotation lift-over tool Liftoff (Shumate and Salzberg 2020). We considered only transcripts with ORFs (i.e. start and stop codon, no frame-shift mutation and no internal stop codons) as successful mappings. Out of the 21,656 transcripts from 20,096 *H. melpomene* annotated genes and 20,118 transcripts from 13,676 *H. erato demophoon* annotated genes, we were able to successfully map 5,817-14,838 *H. melpomene* genes (6,217-16,007 transcripts) and 4,530-9,780 *H. erato demophoon* genes (6,139-14,472 transcripts) - Supplemental Table S5. The success of the gene annotation lift-over approach decreased with phylogenetic distance to the reference. While some of the genes that were not successfully lifted-over could potentially represent mis-annotations in the reference, this could also reflect differences in the structure of these genes or differences in gene composition between species. In fact, Liftoff is designed to map annotations between assemblies of the same or closely-related species and assumes gene structure is conserved between target and reference assemblies. Species-specific *de-novo* gene annotation using transcriptome data would be needed to obtain a more comprehensive annotation for all species.

### Whole mitochondrial genome assemblies

The *de novo* assembly of *Heliconius* mitochondrial genomes allowed the recovery of partially complete mitochondrial sequences (*ca.* 15-kb, typical of *Heliconius*) for all 16 species, including part of the mitochondrial DNA control region. A genealogy based on these mitochondrial genomes (Figure 1A) did not differ from that for the mitochondrial genomes assembled using reference-aided approaches (Kozak et al. 2015; Massardo et al. 2020), thereby validating our *de novo* approach.

### Improved mapping efficiency using the reference-guided assemblies

Mapping the original *w2rap* Illumina short read sequence data to the reference-guided genome assemblies of their own species resulted in 0.45-12.77% more mapped reads and 0.60-40.77% more properly paired reads than when mapping to the closest reference genome (Supplemental Table S6). These mappings also show an increase of the depth of coverage (1.02-2.29 times the coverage obtained when mapped to the closest reference; Supplemental Table S6), which is also more uniform along chromosomes (Supplemental Figs. S40-S41). The largest increases in depth of coverage were observed for *H. burneyi* and *H. doris*, which are the two sequenced *Heliconius* species phylogenetically most distant to either reference genome. Increases in depth of coverage tend to be larger in species in the *erato*/*sara* clade (1.06 to 1.97 times more coverage) than in species in the *melpomene*/silvaniform clade (1.02 to 1.35 times more coverage). This is expected since species sampled within the *erato*/*sara* clade were typically more divergent from *H. erato* than species in the *melpomene*/silvaniform group are from *H. melpomene*.

Importantly, these results show how studies focusing on *Heliconius* species with deeper divergence to both *H. melpomene* and *H. erato* will benefit from mapping re-sequence data to the reference-guided assemblies generated here. Also, the greater uniformity of coverage along chromosomes when mapping reads to the reference-guided assemblies suggests that they should better capture fine-scale structural variation. This likely reflects the ability of the high sequencing fidelity of the original *w2rap* assemblies to resolve short imperfect repeats (< 500 bp long) (Love et al. 2016; Edelman et al. 2019) that differ between species.

### Genome expansions and gene duplications

Although genome sizes vary among *Heliconius* species, the relative but not absolute sizes of chromosomes were generally conserved (Figure 2A, Supplemental Fig. S42). The three closely related species with the largest genomes in the *melpomene/*silvaniform group (*H. hecale, H. elevatus and H. pardalinus*) are exceptions. Upon closer inspection, the variation in chromosome size in these three species is particularly accentuated on chromosome 9 (Figure 2A; Supplemental Fig. S42). Alignment of reference-guided assemblies of these three species to the *H. melpomene* reference genome suggests that the increase in size of chromosome 9 mainly corresponds to a single genomic region in *H. melpomene* (Hmel209001o:5125000-5450000, Figure 2B). This region is *ca*. 325 kb long in *H. melpomene* but the scaffolds that map to it total over 10x as long (3.350-4.125 Mb) in the *hecale*/*elevatus*/*pardalinus* trio.

**Figure 2.**
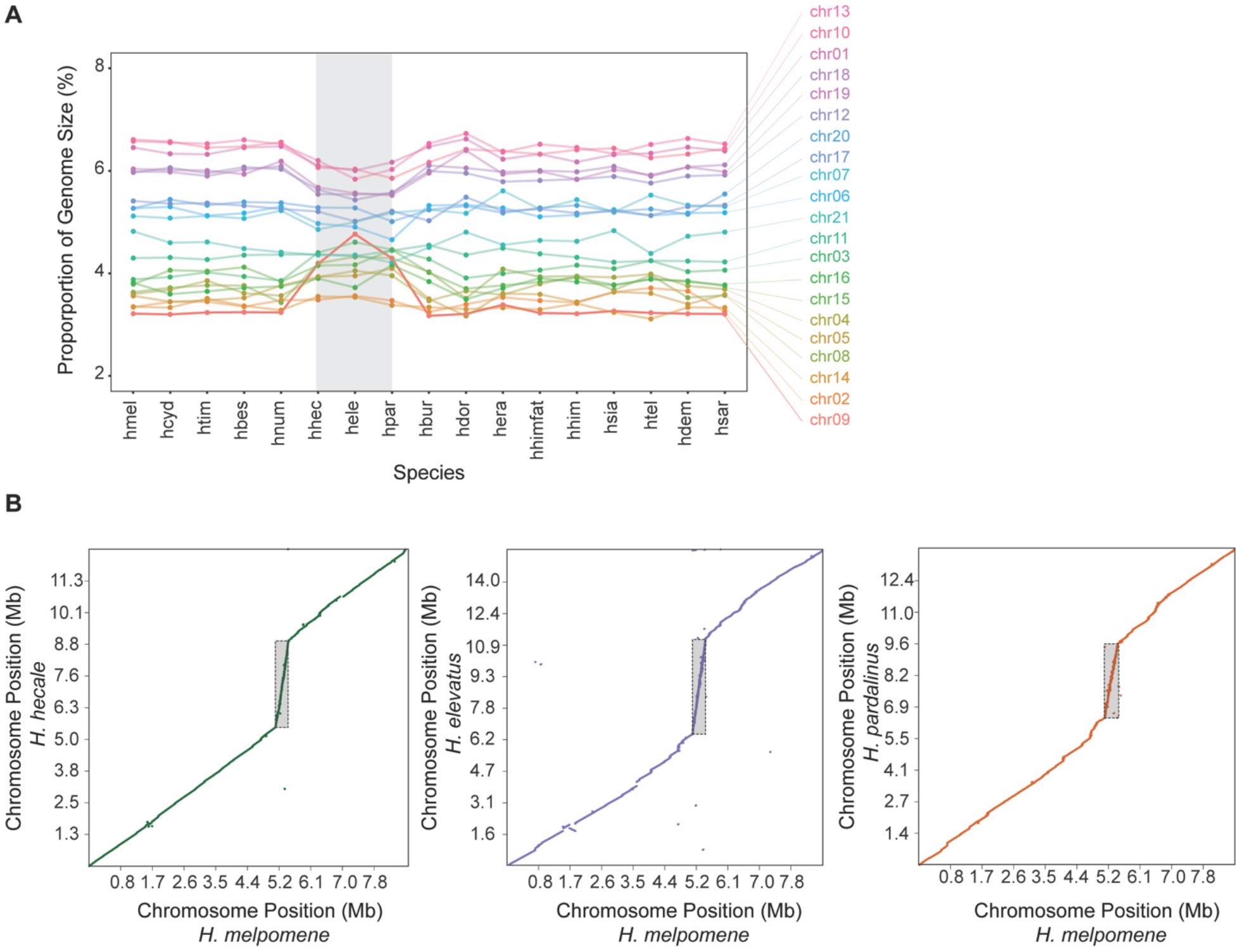
Chromosome size variation and local genomic expansions. **A** – Chromosome sizes in proportion to the genome size across the different species for the reference-guided assemblies mapped to the *H. melpomene* reference genome. Chromosome relative sizes are generally similar across species, with the exception of *H. hecale*, *H. elevatus*, and *H. pardalinus*, particularly chromosome 9. **B** – Genome to genome alignment showing the repeat region on chromosome 9 (highlighted by the grey rectangles) in the species trio: *H. hecale*, *H. elevatus* and *H. pardalinus*.

We investigated whether other genomic regions also underwent an increase in size specifically in these three species. There are four regions that show exceptionally high coverage in these three species (at least 5-fold local increase in the *hecale*/*elevatus*/*pardalinus* trio and less than 2-fold local increase in every other species, in at least two consecutive 25 kb windows). These included the region on chromosome 9 discussed above and three other regions on chromosome 2 (Hmel202001o:4075000-4125000), chromosome 4 (Hmel204001o:5650000-5875000), and chromosome 8 (Hmel208001o:3300000-3475000) (Supplemental Fig. S43, Supplemental Table S7). In contrast, mapping reads onto the reference-guided assemblies resulted in more uniform coverage in these regions (Supplemental Figs. S44-S47). This suggests the repeats are divergent enough so that they could be at least partially resolved in the *w2rap* assemblies.

All repeat regions harbor protein coding genes (Supplemental Table S7), as annotated in the *H. melpomene* reference genome, and thus structural variation in these regions could have resulted in gene copy number variations with potential functional consequences. To address this, we first estimated the copy number of each exon based both on i) the number of valid alignments of *H. melpomene* exon sequences onto the reference-guided assemblies and ii) the normalized mean per base coverage for each exon, mapping *H. hecale*, *H. elevatus* and *H. pardalinus* re-sequencing data to the *H. melpomene* reference. Copy number estimates based on number of exon sequence alignments are generally lower than estimates based on read coverage (Figure 2C; Supplemental Figs. S48-S50). Nevertheless, for both measures, exonic copy number is much larger in *hecale*/*elevatus*/*pardalinus* trio than in *H. melpomene*, suggesting duplications of the corresponding genes. It should also be noted that copy number is variable between exons of the same gene and, and while it can probably be attributed to different alignment efficiency due to variation in exons sequence length (Li 2018), it might also be due to partial duplications of some of these genes. However, given the fragmented nature of the *w2rap* genomes, we could not assess whether genes were wholly or partially duplicated, nor whether the duplications were translocated elsewhere in the genome or are located in the same region as in *H. melpomene*. Long read sequencing data would be required to resolve this.

These gene duplications could result in pseudogenes, in which case we might expect to find stop codons within exons and a relaxation of selection. In general, we find high exon copy numbers even after excluding exon copies with stop codons (10-14% exon copies have a stop codon; Supplemental Figs. S51-S54). Also, dN/dS estimates are overall close to zero, suggestive of purifying selection (Supplemental Figs. S55-S58). RNA-Seq shows a significant correlation between gene copy number and expression levels and that many of these genes have significantly higher expression in *H. pardalinus* than in *H. melpomene* (Figure 2D). Together, these results suggest that many of the gene copies are functional and that CNV at these genes resulted in altered gene dosage.

### Inversions fixed between the two Heliconius major clades

Reference-guided assemblies will inevitably be ineffective at detecting inversions or translocated regions, so it seems important to quantify potential drawbacks of our approach. Previous studies showed that some regions of the genome with unusual phylogenomic patterns in the *erato*/*sara* clade were associated with inversions (Edelman et al. 2019). Here, we make a systematic search for small to medium sized inversion differences among *Heliconius* species, focusing on those 50 kb - 2 Mb long. At the broad scale, the genome structure of the reference-guided assemblies is constrained by the reference genome, so we returned to the *w2rap* scaffolds (after collapsing alternative haplotypes with HaploMerger2), mapping these to the *H. melpomene* and the *H. erato* reference genomes to infer inversion breakpoints. In total, and after filtering, we found 2560 and 3829 scaffolds for which one end aligns to the positive strand of the reference genome and the other end maps to the negative strand, using the *H. melpomene* and *H. erato*, respectively. Of these, 900 and 1786 support inversions 50 kb - 2 Mb long, yielding 345 and 741 unique candidate inversions across all species (mapping to *H. melpomene* and *H. erato demophoon*, respectively), supported by at least one scaffold per species, some of which were shared by multiple species (Supplemental Table S8).

Our systematic search confirmed previous findings of two independent but overlapping introgressed inversions around a color patterning locus on chromosome 15 (one shared by *H. sara*, *H. demeter*, *H. telesiphe* and *H. hecalesia* and the other shared by *H. pardalinus* and *H. numata*) and another inversion on chromosome 2 (shared by *H. erato* and *H. telesiphe*) (Edelman et al. 2019). In addition, we found five moderately large inversions, previously identified as inversion candidates based on alignments between *H. melpomene* and *H. erato* reference genomes (Davey et al. 2017), to be fixed between major branches of the *Heliconius* phylogeny (Figure 4A). Such shared inversions occur on chromosome 2 (Supplemental Fig. S59), chromosome 6 (Supplemental Fig. S60), chromosome 13 (Figure 2C; Supplemental Fig. S61) and the Z chromosome, chromosome 21 (Supplemental Fig. S62; Supplemental Table S9). The two inversions on chromosome 6 occur in tandem and are further supported by linkage maps in *H. melpomene* and *H. erato* (Davey et al. 2017).

The placement of *H. doris* and *H. burneyi* in the *Heliconius* phylogeny remains contentious. These two species have been inferred to be more closely related to the *melpomene/*silvaniform clade than to the *erato*/*sara* clade (see also the mitochondrial tree of Figure 1A), but node supports are relatively weak and the internal branches leading to *H. doris* and *H. burneyi* are short (Kozak et al. 2015). We here test whether homologous inversions can be used as a phylogenetic character to resolve their placement. Figure 1A). Both *H. burneyi* and *H. doris* group with the *melpomene*/silvaniform group based on the orientation of three inversions on chromosomes 2 and 6, but with the *erato*/*sara* clade based on the of homologous inversions on chromosomes 13 and Z. These groupings are further confirmed based on maximum-likelihood (ML) phylogenetic analysis of the inversion regions using a subset of species (Figure 4B). The only exception is that *H. doris* and *H. burneyi* both group with *melpomene*/silvaniform species for the inversion on the Z chromosome in the ML phylogeny, rather than with *erato*/*sara* (Figure 4B), as might be expected solely based on presence/absence of the inversion. (Supplemental Fig. S61). This apparent contradiction can be reconciled if the *melpomene*/silvaniform clade is sister both to *H. doris* and to *H. burneyi*, but the Z chromosome inversion was derived in the *melpomene*/silvaniform ancestor after it split from the *burneyi* and *doris* lineages. In this scenario, the sharing of the inversion between *burneyi*/*doris* and the *erato*/*sara clade* on chromosome 13 must be explained by secondary transfer *via* introgression, perhaps soon after the initial separation of the two major clades, or through incomplete lineage sorting of an inversion polymorphism at the base of *Heliconius*. Previously, reticulation involving *H. burneyi* and *H. doris* and the *erato*/*sara* group had been hypothesized, but different phylogenomic methods gave different results (Kozak et al. 2018).

To test for introgression genome-wide, we used Patterson’s *D*-statistic (Green et al. 2010; Durand et al. 2011). Specifically, we calculated *D* for all possible topologies of the triplets (*H. erato – H. melpomene – H. doris*) and (*H. erato – H. melpomene – H. burneyi*), in each case using *Eueides tales* as an outgroup.

For a given triplet of species, the minimum absolute whole genome Patterson’s *D*-statistic should result for the topology that best describes the relationships between species. We found that this is the case when *H. erato* is the inner outgroup in both triplets, implying that *H. burneyi* and *H. doris* are more closely related to *H. melpomene*. Yet, Patterson’s *D*-statistics are still significantly different from zero (Patterson’s *D* = 0.037 and 0.060 for *H. doris* and *H. burneyi*, respectively) based on block-jackknifing, providing evidence of introgression among lineages leading to *H. burneyi, H. doris,* and *H. erato*. We also used an alternative branch length-based approach, QuIBL (Quantifying Introgression via Branch Lengths; Edelman et al. 2019), which further corroborated these results (Supplemental Table S10). To understand which specific genomic regions were shared by introgression between these species, we estimated the excess of shared derived mutations between *H. doris* and *H. burneyi* with either *H. melpomene* or *H. erato,* using the *f*_*dM*_ statistic (Malinsky et al. 2015). The *f*_*dM*_ estimates in windows overlapping the chromosome 13 inversion show a significant deviation from the genomic average, with an excess of shared variation between *H. erato* and both *H. doris* and *H. burneyi* (Figure 4D; Supplemental Fig. S63). Likewise, relative divergence between *H. erato* to both *H. burneyi* and *H. doris* is significantly reduced in the inversion region (Figure 4D; Supplemental Fig. S64). We also used QuIBL with the triplets (*H. erato – H. melpomene – H. doris*) and (*H. erato – H. melpomene – H. burneyi*) to calculate the likelihood that the discordant phylogenies at the chromosome 13 inversion were due to introgression. For both triplets, the average internal branch of gene trees within the chromosome 13 inversion is larger than the genome-wide average, corresponding to a 90.1% and 86.7% probability of introgression, respectively (Supplemental Fig. S65) We found no significant *f*_*dM*_ or relative divergence estimates for any of the other four inversions, including the Z chromosome inversion. These results strongly support the argument that the chromosome 13 inversion of *H. doris* and *H. burneyi* results from introgression from the common ancestor of the *erato*/*sara* clade.

## Discussion

### Genome assembly improvements and limitations

Here, we implement a purely *in silico* reference-guided scaffolding approach to improve draft genome assemblies of 16 species from across the genus *Heliconius*. The contiguity of our new assemblies is similar to that of the reference genomes. For instance, the *H. melpomene* reference genome assembly has 38 scaffolds anchored to chromosomes (99.1% of the assembly length), and the reference-guided assemblies scaffolded based on this reference have 31-36 scaffolds anchored to chromosomes representing 83.8-99.1% of the total assembly. Similarly, the *H. erato* reference has 195 scaffolds anchored to chromosomes (100% of the assembly length), and the reference-guided assemblies scaffolded based on this reference have 94-168 scaffolds anchored to chromosomes representing 83.2-99.9% of the total assembly.

Our reference-guided assembly strategy assumes that the orientation and order of the new scaffolds in our genomes is the same as the reference. Clearly, it may not fully represent the structure of these genomes. While small genomic rearrangements spanned by the original scaffolds (i.e. rearrangements in relation to the reference present within *w2rap* scaffolds) are recovered in our reference-guided assemblies, larger genomic rearrangements relative to the reference not spanned by a single *w2rap* scaffold can be missed. One such example is the case of the known *ca.* 400 kb inversion around a color pattern locus known from *H. numata* and *H. pardalinus* on chromosome 15 (Jay et al. 2018) which we do not recover in the reference-guided assemblies, in either species. This is also the case for the five large inversions we discovered that are fixed between the two *Heliconius* major clades, depending on the reference genome used to guide scaffolding. For instance, for species in the *melpomene*/silvaniform group, all reference-guided assemblies mapped to the *H. melpomene* reference have the correct orientation for all five inversions, but not when mapped to the *H. erato* reference. The same logic applies for species in the *erato*/*sara* group, when mapped to different references. For *H. burneyi* and *H. doris* however, neither of the two alternative reference-guided assemblies recovers the correct orientation of all five inversions, since these two species share the same orientation as *H. melpomene* for the inversions on chromosome 2 and 6, but not for chromosomes 13 and 21 (for which they have the same orientation as *H. erato*). Long-read sequence data and/or linkage mapping could better resolve the genome structure of species-specific assemblies. Nevertheless, our reference-guided assemblies represent a major improvement over mapping short read data directly to existing reference genomes, and researchers that use these and other reference-guided assemblies for this purpose will see marked improvement in their data quality.

Mapping the original w2rap Illumina reads back to the reference-guided assembly of their own species resulted in more than doubling of the median genomic coverage in some species and in a more uniform depth of coverage along the genome than when mapping to the closest reference genome. Mapping efficiency improves in all species studied here (Supplemental Table S6), but we see the greatest benefits in *H. burneyi* and *H. doris,* the two *Heliconius* species studied here that are most divergent from either reference genome assembly. In these two species, the proportion of properly mapped reads increases from 53.6% and 49.9% (for *H. burneyi* and *H. doris*, respectively) when mapped to the *H. melpomene* reference genome, to 90.7% and 90.6% when mapped to their own reference-guided assembly. In another study (Rosser et al., in preparation), a linkage map produced from backcrosses of F1 male hybrids, between *H. pardalinus butleri* and *H. p. sergestus,* to the parental *H. p. butleri* population contained *ca.* 29% more markers when RADseq data was mapped to the new *H. pardalinus* reference-guided assembly than to the *H. melpomene* reference. The use of reference-guided assemblies of the closest species thus greatly improves the efficiency of mapping resequencing data over mapping to the currently available reference genomes.

The more uniform depth of coverage when mapping to reference-guided assemblies also leads to improvements in discovery of species-specific genomic variation and in resolving imperfect repeat regions. Indeed, given variation in genome sizes among *Heliconius* species (275-418 Mb), the new genomes are helpful in mapping variation that is otherwise lost or mapped to similar but non-orthologous regions of more divergent reference genomes. Variations in depth of coverage along the genome, if not properly filtered, could lead to biased estimates of diversity and divergence. For example, partially divergent repeats mapping to the same region in the reference genome (resulting in unusually high coverage) could inflate local estimates of diversity and thus be spuriously implicated as important sites for species divergence. This is especially likely in studies focusing on *Heliconius* species with larger genomes when mapping reads to the *H. melpomene* reference, the smallest genome assembled here. On the other hand, if regions with abnormal coverage are filtered out, information could be lost by discarding genomic regions with potentially relevant biological signals. For example, highly divergent regions may result in abnormally low coverage, even though such regions could be important for diversification of the group.

Overall, our reference-guided assemblies extend the number of applications for which these genomes can be used. By ordering, orienting and anchoring scaffolds onto chromosomes, the new reference-guided assemblies enable improved chromosome-scale analyses and genome scans.

### Prevalence of structural variants in Heliconius butterflies

Chromosomal rearrangements can play a major role in adaptation and speciation (Wellenreuther and Bernatchez 2018; Feulner and De-Kayne 2017). By reducing recombination, inversions can facilitate the build-up of associations between loci involved in traits responsible for reproductive isolation, and thus could play a role in establishing or reinforcing species barriers (Noor et al. 2001). Inversions can also be favored by selection by maintaining adaptive combinations of locally adapted alleles (Todesco et al. 2020; Faria et al. 2019; Christmas et al. 2019).

In *Heliconius*, a previous study focusing on two closely related species (*H. melpomene* and *H. cydno*) found no evidence for major inversions that might have aided speciation (Davey et al. 2017). Thus, *Heliconius* appeared to have low rates of chromosomal rearrangement, and selection without the help of chromosomal rearrangements was believed to maintain the differences between these two species. In another species, *H. numata*, the tandem inversion complex that forms the supergene locus *P* allows the maintenance of a multi-allele color pattern polymorphism of mimicry morphs (Joron et al. 2011). The first inversion in the tandem supergene was most likely transferred to *H. numata* via introgression from *H. pardalinus* (Jay et al. 2018). An independently derived inversion has since been found for the same colour pattern determination region in four species in the *erato*/*sara* clade (*H. telesiphe*, *H. hecalesia*, *H. demeter* and *H. sara*). This inversion was also inferred to have been shared via introgression, this time between *H. telesiphe* and *H. sara* sub-clades (Edelman et al. 2019). In parallel hybrid zones of *H. erato* and *H. melpomene*, 14 and 19 polymorphic inversions were detected within each species, respectively. Most of these inversion polymorphisms did not differ across the hybrid zones of either species. The frequency of only one inversion on chromosome 2 (different to the inversion on chromosome 2 reported here) differed strongly across the hybrid zone between highland *H. e. notabilis* and lowland *H. e. lativitta* races, and may be associated with ecological adaptation to altitude (Meier et al. 2020).

In the 16 species studied here, we systematically searched for inversions. We found several candidates in all 16 species (17-61 and 40-126 inversions per species, compared with *H. melpomene* and *H. erato*, respectively), including some previously described (Davey et al. 2017; Jay et al. 2018; Edelman et al. 2019). However, the strategy we implemented to search for inversions, i.e. split alignment of *w2rap* scaffolds to forward and reverse strands of the reference genomes, is liable to false positives because small interspersed duplications and translocations (for example due to transposable element activity) might generate a similar signal. This is particularly likely in highly repetitive regions where we find many different, partially overlapping candidate inversions in many or all species (Supplemental Fig. S66). It is thus difficult to assess, solely based on these results, how pervasive inversions are among *Heliconius* species. While it is possible that inversions in this group occur more frequently than earlier studies indicated (Heliconius Genome Consortium 2012; Davey et al. 2017), long-read or linked-read sequencing, preferably with a larger set of individuals per species, will ultimately be needed to answer this question.

However, by focusing on phylogenetically informative inversions, we were able to verify five candidate inversions that occurred deep in the *Heliconius* phylogeny. We searched for inversions fixed between the *melpomene/*silvaniform and *erato*/*sara* groups. We are confident that these were correctly identified for two reasons. First, the inversions are supported in multiple species, with breakpoint coordinates consistent among species. Second, while a mis-assembly in the reference genome could generate a misleading signal of inversion, this is unlikely to happen for the same candidate inversion when mapping to two or more different genomes. All five of these inversions were supported in multiple species when mapping scaffolds to either reference genome, the orientation of the inversion being mirrored depending on the reference used. Furthermore, the inversion orientation shows a phylogenetic signal (fixed between clades) that is unexpected if due to mis-assembly in one of the reference genomes.

The most parsimonious scenario that explains both the orientation and the phylogenetic pattern, taking all five inversions into account, supports the hypothesis that *H. burneyi* and *H. doris* are more closely related to the *melpomene*/silvaniform group than to the *erato*/*sara* group (Figure 4), in line with previous studies (Edelman et al. 2019; Kozak et al. 2018). The relationships of the inversion on chromosome 13, which groups *H. burneyi*, *H. doris* and the *erato*/*sara* group, is then explained by introgression between the ancestor of the latter group and both *H. burneyi* and *H. doris* (Supplemental Figs. S63-S65). Introgression almost certainly occurred from the *erato*/*sara* clade into *H. burneyi* and *H. doris*, since the relative divergence between *H. erato* and both *H. burneyi* and *H. doris* is reduced at the chromosome 13 inversion when compared to the rest of the genome (Figure 4E), but not between *H. erato* and *H. melpomene* as expected if introgression took place in the other direction (Supplemental Fig. S67). Interestingly, *H. burneyi* has been inferred to be on a separate branch from *H. doris*, although the two branches were connected by introgression (Kozak et al. 2015, 2018). This suggests that introgression of the chromosome 13 inversion occurred twice. Either there were two separate introgression events from the *erato*/*sara* ancestor to *H. burneyi* and to *H. doris*, or the inversion first passed from the *erato*/*sara* ancestor to one of these two species which then passed it to the other. Altogether, and in line with previous studies (Edelman et al. 2019; Kozak et al. 2018), this inversion supports a hypothesis that hybridization and introgression among species occurred early in the radiation of *Heliconius*, as well as later, between more closely related species extant within each major subgroup. Alignment issues have previously made it hard to interpret evidence for introgression so deep in the phylogeny. Although we still do not know whether it has functional implications, our finding of transfer of this chromosome 13 inversion provides stronger support for introgression deeper in the Heliconiini tree than hitherto.

Species may also differ in gene copy number. Copy number can affect the phenotype by altering gene dosage, altering the protein sequence, or by creating paralogs that can diverge and gain new functions (Iskow et al. 2012). Copy number variation has been implicated in ecological adaptation – e.g. insecticide resistance in *Anopheles* mosquitoes (Lucas et al. 2019), climate adaptation in white spruce (Prunier et al. 2017) and polar bears (Rinker et al. 2019), and resistance to malaria in humans (Leffler et al. 2017). Gene copy number may also be involved in reproductive barriers among species – e.g. hybrid lethality in *Mimulus* sympatric species (Zuellig and Sweigart 2018). Gene duplications within specific gene families in the branch leading to *Heliconius* have been linked to evolution of visual complexity, development, immunity (Heliconius Genome Consortium 2012) and female oviposition behavior (Briscoe et al. 2013). Within the genus, gene copy number variation is plausibly associated with species divergence between *H. melpomene* and *H. cydno* (Pinharanda et al. 2017).

Here we show that the genomes of different *Heliconius* species vary in size, with each chromosome typically showing similar directional changes in size between species. Thus, genome expansions and reductions in size seem typically to involve all chromosomes, so that the relative sizes of chromosomes are conserved. Our study of the *Heliconius* butterfly radiation conforms, on a much more restricted phylogenetic scale, to the pattern of relative chromosome size across eukaryotes: across many orders of magnitude of genome size, relative chromosome sizes can be predicted based on chromosome number and are almost always between ~0.4x and ~1.9x the mean (Li et al. 2011).

We find that, in *Heliconius*, genomic expansion is at least partially driven by small genomic regions that became hotspots of repeat accumulation. Amplified regions tend to be conserved among closely related species and are more frequent towards chromosome ends (Supplemental Fig. S68). However, in a subclade of three closely related species (*H. hecale*, *H. elevatus* and *H. pardalinus*), we found four small genomic regions with highly aberrant increases in size and exon copy number compared to related species. These three are therefore exceptions to more or less orderly pattern across chromosomes in the rest of the genus. Our approach for detecting exceptional repeat regions relies on the *H. melpomene* genomic arrangement as a backbone. Hence, we do not know whether the additional copies we found were translocated to other regions of the genome of these three species, or whether they remained clustered as tandem copies at a single genomic location. By aligning the reference guided-assemblies to the *H. erato demophoon* reference, we found a signal of local expansion in chromosome 9 (Supplemental Figures S25-27) which would support that the repeats occur in tandem. However, we could not assess whether this was also the case for the repeat regions in the three other chromosomes. Transposable element activity is one possible mechanism responsible for these repeats (Bourque et al. 2018), and rapid divergent transposable element evolution has already been found among *Heliconius* species (Ray et al. 2019). Hybridization could also spread variation in copy number among the species. *H. hecale*, *H. elevatus* and *H. pardalinus* are sympatric in the Amazon where they are known to hybridize occasionally (Mallet et al. 2007; Rosser et al. 2019). We found significantly higher copy numbers in the Amazon than in extra-Amazonian populations of these species (Supplemental Fig. S69). The correlations of copy number among species in an area suggests that hybridization might indeed have been involved.

Genes within highly amplified regions had significantly higher expression levels in *H. pardalinus* than in *H. melpomene* (Figure 3B), which suggests that this gene copy variation could have functional significance. An examination of genes within these regions shows that orthologs of these genes in *Drosophila* are involved in important functions such as cytoskeletal processes and oogenesis (i.e. *Dhc64C*, *sima*, *shotgun*, and *capicua*; Supplemental Table S7). Evaluating how variation in these critical genes impacts phenotypes in *H. pardalinus*, *H. elevatus*, and *H. hecale* will advance our understanding of the role of copy number variation in evolution.

**Figure 3.**
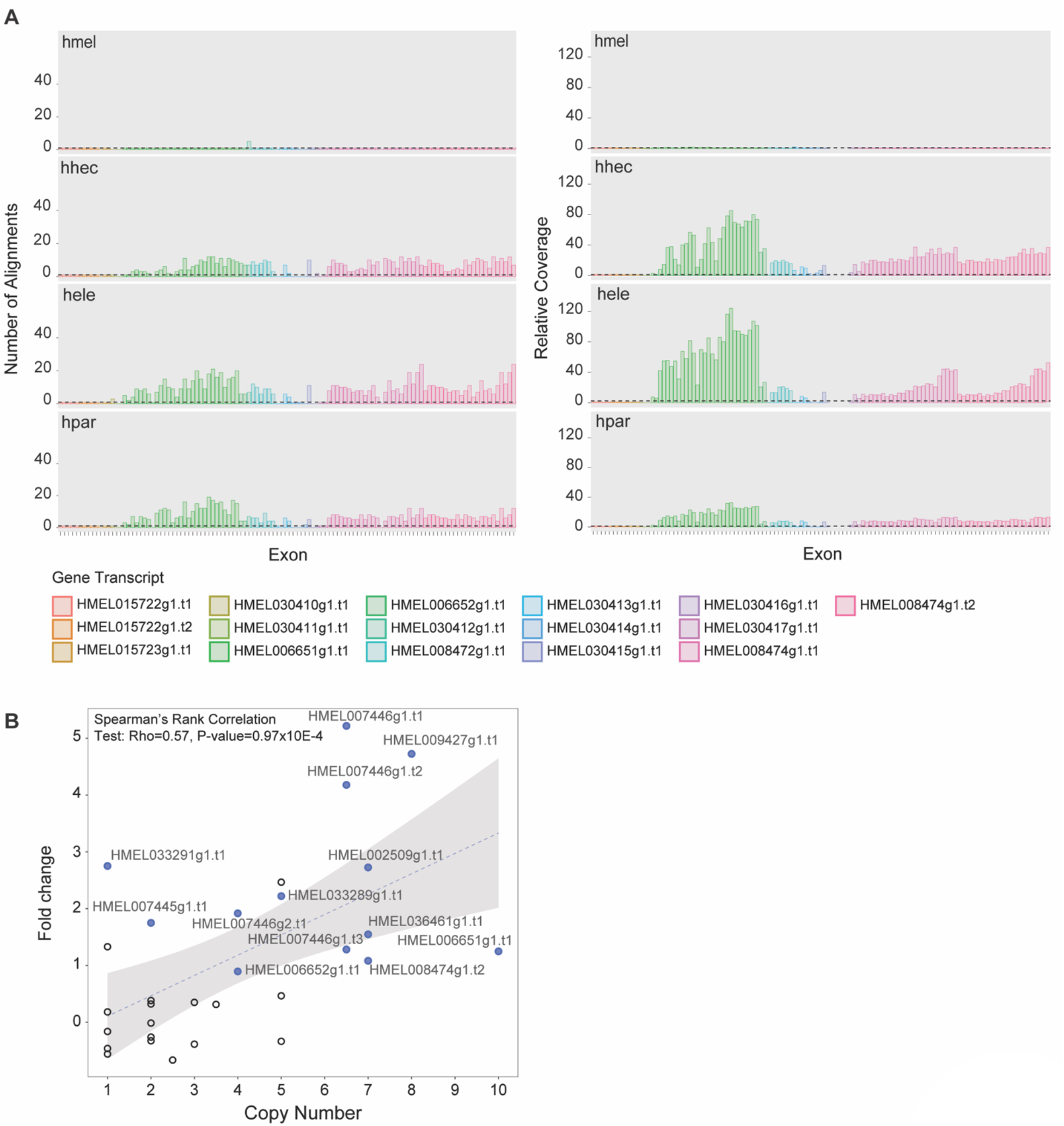
Copy number variation and increased expression levels of genes in the repeat region on chromosome 9. **A** – Exon copy number variation for genes in the chromosome 9 repeat region. The number of alignments (left panel) and relative coverage (right panel) were used as proxies of copy number. Relative coverage was calculated by dividing exon coverage by the median genomic coverage, based on mappings to the *H. melpomene* reference. On the left panel, coloured bars depict the number of alignments to the expected chromosome. Dashed horizontal lines on both plots represent a copy number of one*. Our new H. melpomene* assembly was also included as a control. **B** – Fold change in expression level in *H. pardalinus* compared to *H. melpomene* (y-axis) as a function of *H. pardalinus* transcript copy number (x-axis). For each transcript, copy number was calculated as the median number of alignments across exons for the *H. pardalinus* sample. Full blue circles represent transcripts for which the levels of expression in *H. pardalinus* were significantly higher than in *H. melpomene*. The best fit linear model regression line and confidence intervals are depicted by the dashed line and grey band, respectively. Species codes are as in Figure 1.

**Figure 4.**
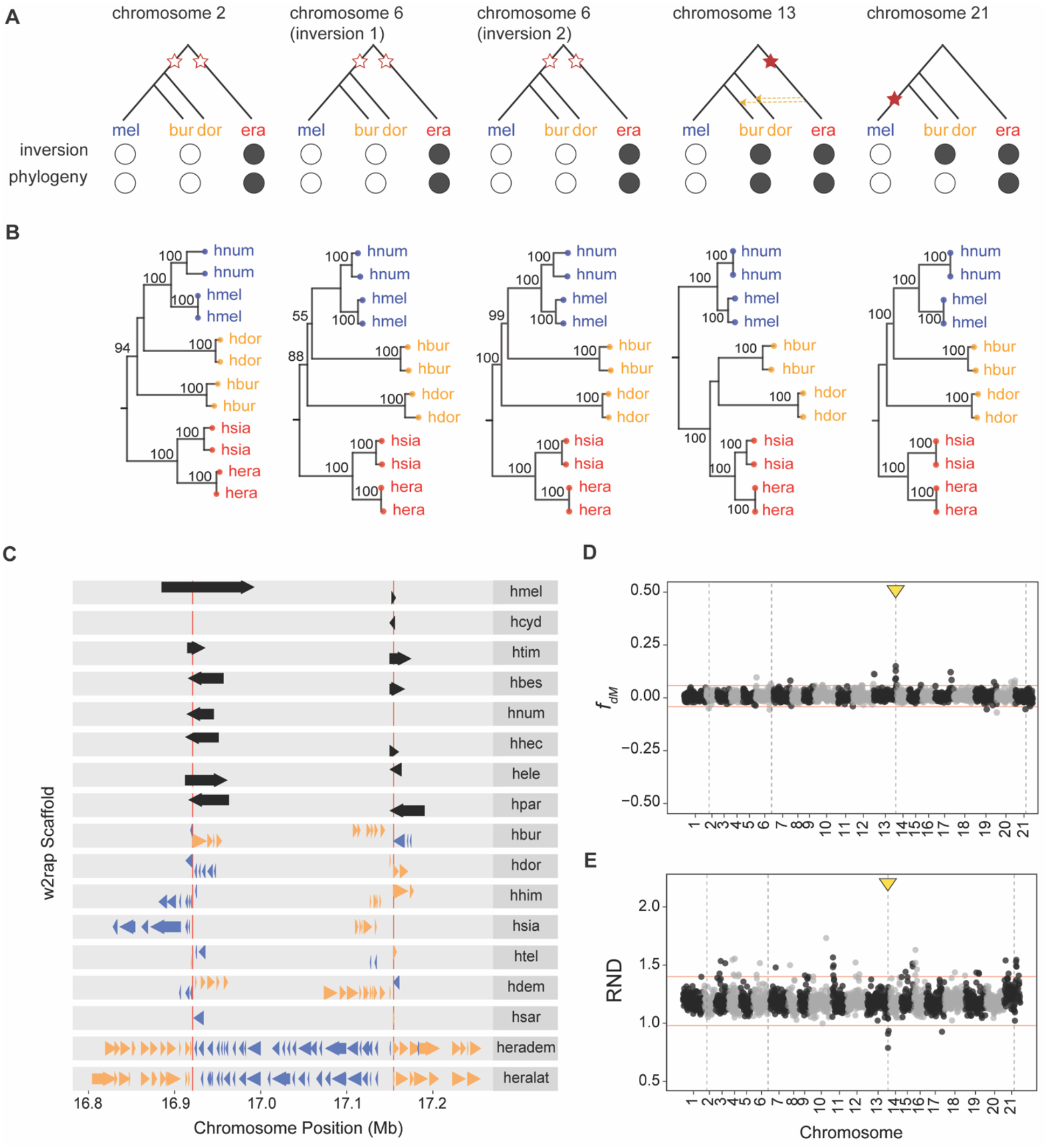
Large Inversions fixed between the *melpomene/* silvaniform and *erato*/ *sara* clades. **A** – Possible scenario for the origin and sharing of the inversion. Stars represent inversions and the branch in which these likely took place. Empty stars are used when the inversion is equally likely to have occurred in two branches. Introgression between branches are represented by arrows, the direction of the arrow indicating directionality. Full and empty circles represent the orientation of the inversion and the groupings based on the phylogeny. **B** – ML phylogenies of each major inversion estimated using IQ-TREE, based on mapping of re-sequence data to the *H. melpomene* reference. **C** – Alignments of scaffolds to the *H. melpomene* reference genome supporting the inversion on chromosome 13. Inversion breakpoints are depicted by the vertical red lines. Scaffold alignments are shown represented by the arrows, the direction and colour of the arrows representing whether the alignments are to the forward strand (blue rightwards arrows) or the reverse strand (yellow leftwards arrows). Black arrows represent alignments spanning the inversion breakpoints. **D,E** – *f*_*dM*_ and relative node depth (RND) statistics along the genome. Both statistics were calculated in 25 kb non-overlapping windows across the genome, based on mapping of re-sequencing data to the *H. melpomene*. Chromosomes are shown with alternating grey and black colours. The location of inversions is given by the dashed vertical lines while horizontal red lines represent +/−3 SD from the mean *f*_*dM*_ and RND values. Outlier windows overlapping the chromosome 13 inversion are indicated by the yellow arrows. Positive *f*_*dM*_ values (and lowered RND) indicate an excess of shared variation between *H. burneyi* with *H. erato* and negative values of *f*_*dM*_ represent an excess of shared with *H. melpomene*. In this test, *H. melpomene* and *H. burneyi* were considered to be the ingroup species and *H. erato* the inner outgroup. Derived alleles were determined using *E. tales*. Species codes are as in Figure 1.

The full extent to which inversions and copy number variation play a role in the evolution of *Heliconius* butterflies remains to be examined. However, the current work suggests that the types of structural variation examined here could be relevant to diversification. The characterization of intra- and interspecific structural variation in this group could thus be an especially promising avenue for future studies particularly now that improvements in sequencing technology allow for more detailed, rigorous and cost-effective detection of structural variants (Wellenreuther et al. 2019; Logsdon et al. 2020).

## Methods

### Genome merging and scaffolding

We used the draft genome scaffolder MEDUSA (Bosi et al. 2015) for reference-aided assembly of the existing DISCOVAR *de novo*/*w2rap* genomes (Edelman et al. 2019). MEDUSA relies on reference genomes from closely related species to determine the correct order and orientation of the draft genome scaffolds, assuming collinearity between reference and the lower contiguity genome. The *w2rap* genome assemblies of 16 *Heliconius* species produced by Edelman et al. (2019) - Supplemental Table S1 - and high-quality reference genome assemblies of two *Heliconius* species - *H. melpomene* (Hmel2.5) and *H. erato demophoon* (Heliconius_erato_demophoon_v1) - were downloaded from lepbase.org. Before the reference-scaffolding step, alternative haplotypes present in the *w2rap* assemblies were collapsed using the HaploMerger2 pipeline (version 20180603; Huang, Kang, & Xu, 2017). Repetitive elements and low complexity regions in the w*2rap* assemblies were first soft-masked using WindowMasker (Morgulis et al. 2006) with default settings. A score matrix for LASTZ (used within HaploMerger) was generated for each *w2rap* assembly. This was done using the lastz_D_Wrapper.pl script with identity = 90 and splitting the *w2rap* assemblies into two sets of scaffolds (scaffolds greater or smaller than 150kb). HaploMerger2 batch scripts A and B were then run using default settings. Finally, MEDUSA was used with default parameters to place and orient the *w2rap* assembly scaffolds based on either of the two reference genomes, placing 100 Ns between adjacent pairs of scaffolds mapping to the same reference chromosome/scaffold. This resulted in two scaffolded assemblies per species (one based on mapping to *H. melpomene* and another based on mapping to *H. erato demophoon* reference genomes).

Reference-guided assemblies were then re-aligned to the *H. melpomene* and *H. erato* reference genomes using the Mashmap aligner as implemented in D-GENIES v1.2.0 online tool (Cabanettes and Klopp 2018) to assess collinearity. Scaffolds in the reference-guided assemblies aligning to reference assembly scaffolds anchored to chromosomes were renamed to reflect their association to chromosomes and order within chromosomes (as in the reference genomes). Also, when necessary, scaffold sequences were reverse-complemented to maintain the same orientation as in the reference.

### Mitochondrial genome assembly

To assemble the mitochondrial genomes the 16 *Heliconius* species analyzed here, we first subsampled 1 million read pairs from the original reads used to produce the *w2rap* assemblies. We then used ABySS 2.0 (Jackman et al. 2017) to assemble the reads, using 5 different *k*-mer sizes (64, 80, 96, 112 and 128-bp) and requiring a minimum mean unitig *k*-mer coverage of 10. All other parameters were left as default. Because of the higher number of mtDNA copies relative to nuclear DNA, resulting in higher mtDNA coverage, we were able to recover the mitochondrial genome as a single large contig (about the size of the complete mitogenome) while any nuclear contigs should be small. In *Heliconius*, the sizes of the mitogenomes sequenced so far are *ca.* 15,300-bp, thus only contigs larger than 15 kb were retained. These where then blasted to the NCBI Nucleotide collection (nr/nt) to confirm that they corresponded to the mitochondrial genome. Finally, for each species, only the largest contig (after removing Ns) was retained. The mitochondrial sequences were aligned using MAFFT v7.407 (Katoh and Standley 2013), with default parameters and a maximum-likelihood (ML) tree was estimated using IQ-TREE v1.6.10 (Nguyen et al. 2015) – Figure 1A. Model selection was performed using ModelFinder (Kalyaanamoorthy et al. 2017) and branch support was assessed with 1000 ultra-fast bootstraps (Hoang et al. 2018), as implemented in IQ-TREE. We also used this approach to recover the mitogenome of *Eueides tales* (Accession number: SRS4612550) to use as an outgroup.

### Scaffolded assemblies quality assessment

Basic statistics (e.g. scaffold N50, cumulative length, proportion of missing sequence) of the reference-guided scaffolded genome assemblies were calculated using QUAST v5.0.2 (Gurevich et al. 2013). Assembly completeness was assessed using BUSCO_V3 (Simão et al. 2015), which looks for the presence (complete, partial or duplicated) or absence (missing) of core arthropod genes (arthropoda-odb9 dataset; https://busco.ezlab.org/datasets/arthropoda_odb9.tar.gz).

### Gene annotation

We used the Liftoff tool (Shumate and Salzberg 2020) to lift gene annotations from the reference genomes to the new reference-guided assemblies. We used either the *H. melpomene* (Hmel2.5.gff3) or *H. erato demophoon* (Heliconius_erato_v1_-_genes.gff.gz) gene annotations (downloaded from www.butterflygenome.org and lepbase.org, respectively), depending on the reference genome used for the scaffolding of the reference-guided assemblies. We ran Liftoff setting the maximum distance between two nodes to be either i) twice the distance between two nodes in the reference genome (i.e. distance scaling factor of 2) or ii) 20 kb distance between in the target, depending on which of these distances is greater. In order to improve mapping of exons at the ends of genes we extended gene sequences by 20% of the gene length, to include flanking sequences on each side (-flank 0.2). Given the *w2rap* scaffolds were ordered, oriented and anchored to chromosomes using the reference genomes as the backbone, and thus we know the association between scaffolds in the reference genomes and in the reference-guided assemblies, we have also enabled the option to first align genes chromosome by chromosome. All other parameters were set as default.

### Mapping and genotype calling of re-sequencing data

Mapping efficiency of the original *w2rap* reads to the reference-guided assemblies was compared with mapping efficiency of the same reads to the reference genomes. Reads were first filtered for Illumina adapters using cutadapt v1.8.1 (Martin 2011) and then mapped to their respective reference-guided genome assemblies, the *H. melpomene* and *H. erato demophoon* reference genomes using BWA mem v0.7.15 (Li 2013), with default parameters and marking short split hits as secondary. Mapped reads were sorted and duplicate reads removed using sambamba v0.6.8 (Tarasov et al. 2015). Realignment around indels was performed with the Genome Analysis Toolkit (GATK) v3.8 RealignerTargetCreator and IndelRealigner modules (McKenna et al. 2010; DePristo et al. 2011), in order to reduce the number of indel miscalls. Mapping statistics and mean read depth were calculated in non-overlapping sliding windows of 25 kb using the *flagstat* and *depth* modules implemented in sambamba v0.6.8, respectively.

Genotype calling was also performed for reads mapped to either of the two reference genomes and for each individual separately with bcftools v1.5 (Li et al. 2009) *mpileup* and *call* modules (Li 2011), using the multiallelic-caller model (call -m) and requiring a minimum base and mapping qualities of 20. Genotypes were filtered using the bcftools *filter* module. Both invariant and variant sites were required to have a minimum quality score (QUAL) of 20. Furthermore, individual genotypes were filtered to have a depth of coverage (DP) >= 8 (except for the Z-chromosome of females for which the minimum required depth was 4) and genotype quality (GQ) >= 20. All genotypes not fulfilling these requirements or within 5-bp of an indel (--SnpGap) were recoded as missing data.

### Copy number variation and selection tests

Copy number variation (CNV) of genes within repeat regions of interest was estimated using two different approaches. The first relies on mapping exonic sequences of genes annotated in the *H. melpomene* reference within regions of interest onto the reference-guided assemblies. The reference-guided assemblies were split back into the original scaffolds by breaking apart regions separated by 100 consecutive Ns, in order to avoid potential mis-mappings over scaffold breakpoints. Exon sequences were mapped to these scaffolds using minimap2 v2.9 (Li 2018), with default settings (except that, as we were interested in repeats, we allowed a much larger threshold of up to 1000 different alignments). Only alignments for which >= 50% of the length of the exon was mapped were considered. Copy number of each exon was then estimated based on the number of alignments to these genomes. The second approach is based on read coverage of the original *w2rap* read data, mapped to the *H. melpomene* reference genome using BWA as described above. For each species, the mean read coverage within an exon (based on the coordinates of exons as annotated in *H. melpomene*) was calculated using the sambamba v0.6.8 ‘*depth’* module (Tarasov et al. 2015). Exon coverage was then normalized dividing by the median genomic coverage (calculated in non-overlapping windows of 25 kb along the genome as described above) to estimate copy number. This second approach was also used to estimate CNV in Amazon and extra-Amazonian populations of *H. hecale*, *H. elevatus* and *H. pardalinus* (Supplemental Table S11).

We further investigated whether CNV in specific genes resulted in potentially functional copies or pseudogenization by analyzing signals of codon-based selection and looking for the presence or absence of stop codons. For each gene we examine each exon independently since different exons can show different copy number. Sequences of the different putative copies were extracted from the reference-guided assemblies, based on the coordinates obtained by aligning the reference *H. melpomene* exon sequences to the reference-guided assemblies (as described above in this section). When shorter than the exon length, coordinates were extended to match the total exon length. Exon sequences including 10 consecutive Ns (introduced during the *w2rap* assembly process) were excluded from this analysis to avoid artificial sequence frameshifts. The remaining exonic sequences of all species were then aligned to the *H. melpomene* reference genome using MAFFT v7.407 (Katoh and Standley 2013), with default parameters and allowing reverse complementing of sequences when necessary. Bases before the start and after the end of the *H. melpomene* reference sequence were removed from the alignment since these could have been erroneously included when extending sequences to match the total exon length (see above). Also, alignments including frameshift mutations (determined based on the *H. melpomene* sequence) were excluded. We then calculated the ratio of non-synonymous versus synonymous changes (dN/dS) for each pairwise comparison between exon copies detected in the reference-guided assemblies and the reference *H. melpomene* sequence, using Li’s (1993) method implemented in the ‘*seqinr*’ package in R. Finally, we checked for the presence of stop codons using a custom script.

### Detection of inversions in the w2rap assemblies

In order to detect potential inversions in relation to the reference genomes, we mapped the *w2rap* scaffolds (after filtering with HaploMerger2; see above) onto the reference genomes. Scaffolds of at least 5 kb were mapped to the *H. m. melpomene* and the *H. erato demophoon* reference genomes using minimap2 (Li 2018) with default settings. Only primary alignments (tp:A:P), at least 1 kb long, with mapping quality >= 60 and with less than 25% approximate per-base sequence divergence (dv) to the reference were kept. Mappings of scaffolds spanning inversion breakpoints in the reference genome should result in split alignments to different strands. We thus considered scaffolds as potentially informative for inversions if they had at least two alignments to the same chromosome (split-alignments) and at least one alignment to each strand as potentially informative for inversions. Same-scaffold alignments mapping to the same strand, partially overlapping or not more than 50 kb apart were concatenated. If less than 20% of the length of the scaffold aligned to the reference, the scaffold was excluded. Furthermore, any scaffolds for which both forward and reverse alignments to the reference i) come from overlapping scaffold regions (overlap greater than 5 kb), ii) overlap in the reference by more than 5 kb or iii) in which the alignment in one strand is completely within the alignment to the other strand, were removed as these likely represent spurious alignments, perhaps due to repeats. Candidate inversions less than 50 kb from scaffold boundaries within chromosomes of the reference genome were also excluded. Finally, we considered any two informative scaffolds to support the same candidate inversion if they overlapped by at least 75% of the maximum length of the two. We also mapped the two reference genomes against each other (and also the *H. erato lativitta* onto both) using minimap2 and inferred candidate inversions by looking for alignments, within a scaffold, to the reverse strand. Only alignments with a MQ >= 10 and to the same chromosome in the reference were considered. Entire scaffolds aligning to the reverse strand are possibly mis-oriented and were not considered to be inversions.

For each candidate inversion we made sequence alignments for a subset of species (*H. melpomene*, *H. numata*, *H. doris*, *H. burneyi*, *H. erato* and *H. hecalesia*, using *Eueides tales* as an outgroup) based on the original *w2rap* sequencing data mapped to both *H. melpomene* and *H. erato* reference genomes. We then estimated maximum-likelihood (ML) trees for these candidate regions using IQ-TREE v1.6.10 (Nguyen et al. 2015). Model selection was performed using ModelFinder (Kalyaanamoorthy et al. 2017) and branch support was assessed with 1000 ultra-fast bootstraps (Hoang et al. 2018), as implemented in IQ-TREE.

We used the Patterson’s *D* statistic (Green et al. 2010; Durand et al. 2011) to test i) which branching pattern best describes the relationships between *H. doris, H. burney*, the *erato*/*sara* and the *melpomene/*silvaniform groups and ii) whether the alternative clustering of *H. doris* and *H. burney* with either of the two groups (both patterns were observed in the inversions) could be explained by introgression. We used the ABBABABAwindows.py script (available from github.com/simonhmartin/genomics_general) to estimate the *D* statistic in non-overlapping windows of 1 Mb, discarding all windows with fewer than 100 informative sites. The mean and variance of the *D statistic* were calculated using a 1-Mb block jackknifing approach, allowing to test whether *D* differed significantly from zero. We have also used the internal branch length based approach QuIBL (Edelman et al. 2019), which uses the distribution of internal branch lengths and calculates the likelihood that the triplet topologies discordant from the species tree are due to introgression rather than ILS alone. For this analysis, we sampled 10 kb windows along the genome (50 kb apart) and for each we estimated maximum-likelihood trees using the phyml_sliding_windows.py (available from github.com/simonhmartin/genomics_general). Only alignments with less than 5% of the sites genotyped were discarded. We then ran QuIBL on the filtered dataset with default parameters and adjusting the number of steps to 50. In both Patterson’s *D* and QuIBL analyses, *Eueides tales* was used as outgroup.

In order to detect local signals of introgression we also calculated the *f*_*dM*_ statistic (Malinsky et al. 2015), which, like the *f*_*d*_ statistic (Martin et al. 2015), checks for imbalance in the number of shared variants between the inner outgroup population and one of two ingroup populations, and was developed specifically to investigate introgression of small genomic regions. Unlike the *f*_*d*_ statistic, it simultaneously tests for an excess of shared variation between the inner outgroup population and either ingroup population, at each genomic window. Again, we used the ABBABABAwindows.py script (available from github.com/simonhmartin/genomics_general) to estimate the *f*_*dM*_ in non-overlapping windows of 100 kb, discarding all windows with fewer than 100 informative sites. Because a local excess of derived alleles could also be explained by retention of ancestral polymorphism (incomplete lineage sorting - ILS), we calculated the divergence (*D*_*XY*_) between both *H. doris* and *H. burneyi* to *H. erato*, normalized by divergence to *H. melpomene* (i.e. Relative Node Depth, RND), to control for variation in substitution rate across the genome. *D*_*XY*_ was calculated in 100 kb non-overlapping windows using the popgenWindows.py script (available from github.com/simonhmartin/genomics_general). Finally, we also used QuIBL to estimate the probability that gene trees within the chromosome 13 inversion were generated by introgression.

### Gene expression analyses

Ovaries were dissected from adult females of *H. melpomene rosina* and *H. pardalinus butleri* at two weeks post-eclosion, divided into developmental stages, and stored in RNALater. Ovaries were blotted dry with kimWipes to remove excess RNALater solution. Tissue was then transferred to TRIZOL and homogenized with the PRO200 tissue homogenizer (PRO Scientific). RNA was extracted with the Direct-zol RNA miniprep kit (Zymo R2051). mRNA libraries were prepared by the Harvard University Bauer Core with the KAPA mRNA HyperPrep kit, with mean fragment insert sizes of 200-300bp. mRNA was sequenced with the NovaSeq S2, producing an average of 49 million paired-end, 50 bp reads.

RNASeq reads were mapped to the *H. melpomene* v2.5 transcriptome (Pinharanda et al. 2019) using kallisto (Bray et al. 2016). Analysis was carried out in R using the Sleuth package (Pimentel et al. 2017). Significant differences in expression levels between *H. melpomene* and *H. pardalinus* were assessed with a likelihood ratio test, comparing expression as a function of developmental stage to expression as a function of developmental stage + species identity.

## Data access

On publication, the reference-guided assemblies and gene annotations generated in this study will have been made available in Zenodo and all custom scripts used in this study will be made available on the GitHub repository https://github.com/FernandoSeixas/HeliconiusReferenceGuidedAssemblies.

## Acknowledgements

We thank the Harvard FAS Research Computing team for their support, A. Shumate for her guidance using the Liftoff software, J. Davey and D. Ray for their valuable inputs to our thinking around genome structural variation, and N. Rosser for the helpful discussions on *Heliconius*. This project was funded by a SPARC Grant from the Broad Institute of Harvard and MIT and funds from Harvard University

## Bibliography

Alonge M, Soyk S, Ramakrishnan S, Wang X, Goodwin S, Sedlazeck FJ, Lippman ZB, Schatz MC. 2019. RaGOO: fast and accurate reference-guided scaffolding of draft genomes. Genome Biol 20: 224.

Benevenuto J, Ferrão LF V., Amadeu RR, Munoz P. 2019. How can a high-quality genome assembly help plant breeders? Gigascience 8: 1–4.

Bosi E, Donati B, Galardini M, Brunetti S, Sagot M-F, Lió P, Crescenzi P, Fani R, Fondi M. 2015. MeDuSa: a multi-draft based scaffolder. Bioinformatics 31: 2443–2451.

Bourque G, Burns KH, Gehring M, Gorbunova V, Seluanov A, Hammell M, Imbeault M, Izsvák Z, Levin HL, Macfarlan TS, et al. 2018. Ten things you should know about transposable elements. Genome Biol 19: 199.

Bray NL, Pimentel H, Melsted P, Pachter L. 2016. Near-optimal probabilistic RNA-seq quantification. Nat Biotechnol 34: 525–527.

Briscoe AD, Macias-Muñoz A, Kozak KM, Walters JR, Yuan F, Jamie GA, Martin SH, Dasmahapatra KK, Ferguson LC, Mallet J, et al. 2013. Female Behaviour Drives Expression and Evolution of Gustatory Receptors in Butterflies. PLoS Genet 9: e1003620.

Cabanettes F, Klopp C. 2018. D-GENIES: dot plot large genomes in an interactive, efficient and simple way. PeerJ 6: e4958.

Christmas MJ, Wallberg A, Bunikis I, Olsson A, Wallerman O, Webster MT. 2019. Chromosomal inversions associated with environmental adaptation in honeybees. Mol Ecol 28: 1358–1374.

Clavijo B, Accinelli GG, Wright J, Heavens D, Barr K, Yanes L, Di-Palma F. 2017. W2RAP: a pipeline for high quality, robust assemblies of large complex genomes from short read data. bioRxiv.

Coyne JA. 2018. “Two Rules of Speciation” revisited. Mol Ecol 27: 3749–3752.

Coyne JA, Orr AH. 1989. Two rules of speciation. In *Speciation and its Consequences* (eds. D. Otte and J.A. Endler), pp. 180–207, Sinauer Associates, Sunderland, MA.

Davey JW, Barker SL, Rastas PM, Pinharanda A, Martin SH, Durbin R, McMillan WO, Merrill RM, Jiggins CD. 2017. No evidence for maintenance of a sympatric *Heliconius* species barrier by chromosomal inversions. Evol Lett 1: 138–154.

DePristo MA, Banks E, Poplin R, Garimella KV, Maguire JR, Hartl C, Philippakis AA, del Angel G, Rivas MA, Hanna M, et al. 2011. A framework for variation discovery and genotyping using next-generation DNA sequencing data. Nat Genet 43: 491–8.

Deschamps S, Zhang Y, Llaca V, Ye L, Sanyal A, King M, May G, Lin H. 2018. A chromosome-scale assembly of the sorghum genome using nanopore sequencing and optical mapping. Nat Commun 9: 4844.

Durand EY, Patterson N, Reich D, Slatkin M. 2011. Testing for Ancient Admixture between Closely Related Populations. Mol Biol Evol 28: 2239–2252.

Edelman NB, Frandsen PB, Miyagi M, Clavijo B, Davey JW, Dikow RB, García-Accinelli G, Van Belleghem SM, Patterson N, Neafsey DE, et al. 2019. Genomic architecture and introgression shape a butterfly radiation. Science 366: 594–599.

Ellegren H, Smeds L, Burri R, Olason PI, Backström N, Kawakami T, Künstner A, Mäkinen H, Nadachowska-Brzyska K, Qvarnström A, et al. 2012. The genomic landscape of species divergence in *Ficedula* flycatchers. Nature 491: 756–760.

Faria R, Chaube P, Morales HE, Larsson T, Lemmon AR, Lemmon EM, Rafajlović M, Panova M, Ravinet M, Johannesson K, et al. 2019. Multiple chromosomal rearrangements in a hybrid zone between *Littorina saxatilis* ecotypes. Mol Ecol 28: 1375–1393.

Feulner PGD, De-Kayne R. 2017. Genome evolution, structural rearrangements and speciation. J Evol Biol 30: 1488–1490.

Fontaine MC, Pease JB, Steele A, Waterhouse RM, Neafsey DE, Sharakhov IV, Jiang X, Hall AB, Catteruccia F, Kakani E, et al. 2015. Extensive introgression in a malaria vector species complex revealed by phylogenomics. Science 347: 1258524.

Ghurye J, Pop M. 2019. Modern technologies and algorithms for scaffolding assembled genomes. PLOS Comput Biol 15: e1006994.

Gopalakrishnan S, Samaniego Castruita JA, Sinding M-HS, Kuderna LFK, Räikkönen J, Petersen B, Sicheritz-Ponten T, Larson G, Orlando L, Marques-Bonet T, et al. 2017. The wolf reference genome sequence (*Canis lupus lupus*) and its implications for Canis spp. population genomics. BMC Genomics 18: 495.

Green RE, Krause J, Briggs AW, Maricic T, Stenzel U, Kircher M, Patterson N, Li H, Zhai W, Fritz MH-Y, et al. 2010. A draft sequence of the Neandertal genome. Science 328: 710–22.

Gurevich A, Saveliev V, Vyahhi N, Tesler G. 2013. QUAST: quality assessment tool for genome assemblies. Bioinformatics 29: 1072–5.

Hoang DT, Chernomor O, von Haeseler A, Minh BQ, Vinh LS. 2018. UFBoot2: Improving the Ultrafast Bootstrap Approximation. Mol Biol Evol 35: 518–522.

Huang S, Kang M, Xu A. 2017. HaploMerger2: rebuilding both haploid sub-assemblies from high-heterozygosity diploid genome assembly. Bioinformatics 33: 2577–2579.

Iskow RC, Gokcumen O, Lee C. 2012. Exploring the role of copy number variants in human adaptation. Trends Genet 28: 245–257.

Jackman SD, Vandervalk BP, Mohamadi H, Chu J, Yeo S, Hammond SA, Jahesh G, Khan H, Coombe L, Warren RL, et al. 2017. ABySS 2.0: resource-efficient assembly of large genomes using a Bloom filter. Genome Res 27: 768–777.

Jay P, Whibley A, Frézal L, Rodríguez de Cara MÁ, Nowell RW, Mallet J, Dasmahapatra KK, Joron M. 2018. Supergene Evolution Triggered by the Introgression of a Chromosomal Inversion. Curr Biol 28: 1839–1845.

Jiggins CD, Mavarez J, Beltrán M, McMillan WO, Johnston JS, Bermingham E. 2005. A genetic linkage map of the mimetic butterfly *Heliconius melpomene*. Genetics 171: 557–570.

Joron M, Frezal L, Jones RT, Chamberlain NL, Lee SF, Haag CR, Whibley A, Becuwe M, Baxter SW, Ferguson L, et al. 2011. Chromosomal rearrangements maintain a polymorphic supergene controlling butterfly mimicry. Nature 477: 203–206.

Kalyaanamoorthy S, Minh BQ, Wong TKF, von Haeseler A, Jermiin LS. 2017. ModelFinder: fast model selection for accurate phylogenetic estimates. Nat Methods 14: 587–589.

Katoh K, Standley DM. 2013. MAFFT Multiple Sequence Alignment Software Version 7: Improvements in Performance and Usability. Mol Biol Evol 30: 772–780.

Kozak KM, McMillan O, Joron M, Jiggins CD. 2018. Genome-wide admixture is common across the *Heliconius* radiation. bioRxiv doi: 10.1101/414201.

Kozak KM, Wahlberg N, Neild AFE, Dasmahapatra KK, Mallet J, Jiggins CD. 2015. Multilocus Species Trees Show the Recent Adaptive Radiation of the Mimetic *Heliconius* Butterflies. Syst Biol 64: 505–524.

Leffler EM, Band G, Busby GBJ, Kivinen K, Le QS, Clarke GM, Bojang KA, Conway DJ, Jallow M, Sisay-Joof F, et al. 2017. Resistance to malaria through structural variation of red blood cell invasion receptors. Science 356: 1140–1152.

Lewis JJ, Reed RD. 2019. Genome-Wide Regulatory Adaptation Shapes Population-Level Genomic Landscapes in *Heliconius*. Mol Biol Evol 36: 159–173.

Lewis JJ, van der Burg KRL, Mazo-Vargas A, Reed RD. 2016. ChIP-Seq-Annotated *Heliconius erato* Genome Highlights Patterns of cis-Regulatory Evolution in Lepidoptera. Cell Rep 16: 2855–2863.

Li H. 2011. A statistical framework for SNP calling, mutation discovery, association mapping and population genetical parameter estimation from sequencing data. Bioinformatics 27: 2987–2993.

Li H. 2013. Aligning sequence reads, clone sequences and assembly contigs with BWA-MEM. arXiv Prepr. arXiv 0: 3.

Li H. 2018. Minimap2: pairwise alignment for nucleotide sequences. Bioinformatics 34: 3094–3100.

Li H, Handsaker B, Wysoker A, Fennell T, Ruan J, Homer N, Marth G, Abecasis G, Durbin R. 2009. The Sequence Alignment/Map format and SAMtools. Bioinformatics 25: 2078–2079.

Li W-H. 1993. Unbiased estimation of the rates of synonymous and nonsynonymous substitution. J Mol Evol 36: 96–99.

Li X, Zhu C, Lin Z, Wu Y, Zhang D, Bai G, Song W, Ma J, Muehlbauer GJ, Scanlon MJ, et al. 2011. Chromosome Size in Diploid Eukaryotic Species Centers on the Average Length with a Conserved Boundary. Mol Biol Evol 28: 1901–1911.

Logsdon GA, Vollger MR, Eichler EE. 2020. Long-read human genome sequencing and its applications. Nat Rev Genet 21: 597–614.

Love RR, Weisenfeld NI, Jaffe DB, Besansky NJ, Neafsey DE. 2016. Evaluation of DISCOVAR de novo using a mosquito sample for cost-effective short-read genome assembly. BMC Genomics 17: 187.

Lucas ER, Miles A, Harding NJ, Clarkson CS, Lawniczak MKN, Kwiatkowski DP, Weetman D, Donnelly MJ. 2019. Whole-genome sequencing reveals high complexity of copy number variation at insecticide resistance loci in malaria mosquitoes. Genome Res 29: 1250–1261.

Malinsky M, Challis RJ, Tyers AM, Schiffels S, Terai Y, Ngatunga BP, Miska EA, Durbin R, Genner MJ, Turner GF. 2015. Genomic islands of speciation separate cichlid ecomorphs in an East African crater lake. Science 350: 1493–1498.

Mallet J, Beltrán M, Neukirchen W, Linares M. 2007. Natural hybridization in heliconiine butterflies: the species boundary as a continuum. BMC Evol Biol 7: 28.

Markelz RJC, Covington MF, Brock MT, Devisetty UK, Kliebenstein DJ, Weinig C, Maloof JN. 2017. Using RNA-Seq for genomic scaffold placement, correcting assemblies, and genetic map creation in a common *Brassica rapa* mapping population. G3 Genes, Genomes, Genet 7: 2259–2270.

Martin M. 2011. Cutadapt removes adapter sequences from high-throughput sequencing reads. EMBnet.journal 17: 10.

Martin SH, Davey JW, Jiggins CD. 2015. Evaluating the Use of ABBA–BABA Statistics to Locate Introgressed Loci. Mol Biol Evol 32: 244–257.

Martin SH, Davey JW, Salazar C, Jiggins CD. 2019. Recombination rate variation shapes barriers to introgression across butterfly genomes. PLOS Biol 17: e2006288.

Masly JP, Presgraves DC. 2007. High-Resolution Genome-Wide Dissection of the Two Rules of Speciation in *Drosophila*. PLoS Biol 5: e243.

Massardo D, VanKuren NW, Nallu S, Ramos RR, Ribeiro PG, Silva-Brandão KL, Brandão MM, Lion MB, Freitas AVL, Cardoso MZ, et al. 2020. The roles of hybridization and habitat fragmentation in the evolution of Brazil’s enigmatic longwing butterflies, *Heliconius nattereri* and *H. hermathena*. BMC Biol 18: 84.

McKenna A, Hanna M, Banks E, Sivachenko A, Cibulskis K, Kernytsky A, Garimella K, Altshuler D, Gabriel S, Daly M, et al. 2010. The genome analysis toolkit: A MapReduce framework for analyzing next-generation DNA sequencing data. Genome Res 20: 1297–1303.

Meier JI, Salazar PA, Ku M, Davies RW, Dréau A, Aldás I, Power OB, Nadeau NJ, Bridle JR, Rolian C, et al. 2020. Haplotype tagging reveals parallel formation of hybrid races in two butterfly species. bioRxiv doi: 10.1101/2020.05.25.113688.

Morgulis A, Michael Gertz E, Schä AA, Agarwala R. 2006. WindowMasker: window-based masker for sequenced genomes. 22: 134–141.

Nguyen L-T, Schmidt HA, von Haeseler A, Minh BQ. 2015. IQ-TREE: A Fast and Effective Stochastic Algorithm for Estimating Maximum-Likelihood Phylogenies. Mol Biol Evol 32: 268–274.

Noor MAF, Grams KL, Bertucci LA, Reiland J. 2001. Chromosomal inversions and the reproductive isolation of species. Proc Natl Acad Sci 98: 12084–12088.

Pimentel H, Bray NL, Puente S, Melsted P, Pachter L. 2017. Differential analysis of RNA-seq incorporating quantification uncertainty. Nat Methods 14: 687–690.

Pinharanda A, Martin SH, Barker SL, Davey JW, Jiggins CD. 2017. The comparative landscape of duplications in *Heliconius melpomene* and *Heliconius cydno*. Heredity 118: 78–87.

Pinharanda A, Rousselle M, Martin SH, Hanly JJ, Davey JW, Kumar S, Galtier N, Jiggins CD. 2019. Sexually dimorphic gene expression and transcriptome evolution provide mixed evidence for a fast-Z effect in Heliconius. J Evol Biol 32: 194–204.

Prowell PD. 1998. Sex linkage and speciation in Lepidoptera. In *Endless Forms. Species and Speciation* (eds. D.J. Howard and S.H. Berlocher), pp. 309–319, Oxford University Press, New York.

Prüfer K, Stenzel U, Hofreiter M, Pääbo S, Kelso J, Green RE. 2010. Computational challenges in the analysis of ancient DNA. Genome Biol 11: R47.

Prunier J, Caron S, Lamothe M, Blais S, Bousquet J, Isabel N, MacKay J. 2017. Gene copy number variations in adaptive evolution: The genomic distribution of gene copy number variations revealed by genetic mapping and their adaptive role in an undomesticated species, white spruce (*Picea glauca*). Mol Ecol 26: 5989–6001.

Ray DA, Grimshaw JR, Halsey MK, Korstian JM, Osmanski AB, Sullivan KAM, Wolf KA, Reddy H, Foley N, Stevens RD, et al. 2019. Simultaneous TE Analysis of 19 Heliconiine Butterflies Yields Novel Insights into Rapid TE-Based Genome Diversification and Multiple SINE Births and Deaths. Genome Biol Evol 11: 2162–2177.

Rice ES, Green RE. 2019. New Approaches for Genome Assembly and Scaffolding. Annu Rev Anim Biosci 7: 17–40.

Rinker DC, Specian NK, Zhao S, Gibbons JG. 2019. Polar bear evolution is marked by rapid changes in gene copy number in response to dietary shift. Proc Natl Acad Sci 116: 13446–13451.

Rosser N, Queste LM, Cama B, Edelman NB, Mann F, Mori Pezo R, Morris J, Segami C, Velado P, Schulz S, et al. 2019. Geographic contrasts between pre- and postzygotic barriers are consistent with reinforcement in *Heliconius* butterflies. Evolution 73: 1821–1838.

Schumer M, Xu C, Powell DL, Durvasula A, Skov L, Holland C, Blazier JC, Sankararaman S, Andolfatto P, Rosenthal GG, et al. 2018. Natural selection interacts with recombination to shape the evolution of hybrid genomes. Science 3684: eaar3684.

Seixas FA, Boursot P, Melo-Ferreira J. 2018. The genomic impact of historical hybridization with massive mitochondrial DNA introgression. Genome Biol 19: 91.

Shumate A, Salzberg SL. 2020. Liftoff: an accurate gene annotation mapping tool. bioRxiv doi: 2020.06.24.169680.

Simão FA, Waterhouse R, Ioannidis P, Kriventseva E V., Zdobnov EM. 2015. BUSCO: Assessing genome assembly and annotation completeness with single-copy orthologs. Bioinformatics 31: 3210–3212.

Tarasov A, Vilella AJ, Cuppen E, Nijman IJ, Prins P. 2015. Genome analysis Sambamba: fast processing of NGS alignment formats. Bioinformatics 31: 2032–2034.

The *Heliconius* Genome Consortium. 2012. Butterfly genome reveals promiscuous exchange of mimicry adaptations among species. Nature 487: 94–98.

Tobler A, Kapan D, Flanagan NS, Gonzalez C, Peterson E, Jiggins CD, Johntson JS, Heckel DG, McMillan WO. 2005. First-generation linkage map of the warningly colored butterfly *Heliconius erato*. Heredity 94: 408–417.

Todesco M, Owens GL, Bercovich N, Légaré JS, Soudi S, Burge DO, Huang K, Ostevik KL, Drummond EBM, Imerovski I, et al. 2020. Massive haplotypes underlie ecotypic differentiation in sunflowers. Nature.

Van Belleghem SM, Rastas P, Papanicolaou A, Martin SH, Arias CF, Supple MA, Hanly JJ, Mallet J, Lewis JJ, Hines HM, et al. 2017. Complex modular architecture around a simple toolkit of wing pattern genes. Nat Ecol Evol 1: 0052.

Wei S, Yang Y, Yin T. 2020. The chromosome-scale assembly of the willow genome provides insight into Salicaceae genome evolution. Hortic Res 7: 45.

Weisenfeld NI, Yin S, Sharpe T, Lau B, Hegarty R, Holmes L, Sogoloff B, Tabbaa D, Williams L, Russ C, et al. 2014. Comprehensive variation discovery in single human genomes. Nat Genet 46: 1350–1355.

Wellenreuther M, Bernatchez L. 2018. Eco-Evolutionary Genomics of Chromosomal Inversions. Trends Ecol Evol 33: 427–440.

Wellenreuther M, Mérot C, Berdan E, Bernatchez L. 2019. Going beyond SNPs: The role of structural genomic variants in adaptive evolution and species diversification. Mol Ecol 28: 1203–1209.

Yang J, Wan W, Xie M, Mao J, Dong Z, Lu S, He J, Xie F, Liu G, Dai X, et al. 2020. Chromosome-level reference genome assembly and gene editing of the dead-leaf butterfly *Kallima inachus*. Mol Ecol Resour 20: 1080–1092.

Yu A, Li F, Xu W, Wang Z, Sun C, Han B, Wang Y, Wang B, Cheng X, Liu A. 2019. Application of a high-resolution genetic map for chromosome-scale genome assembly and fine QTLs mapping of seed size and weight traits in castor bean. Sci Rep 9: 1–11.

Zuellig MP, Sweigart AL. 2018. Gene duplicates cause hybrid lethality between sympatric species of *Mimulus*. PLoS Genet 14: 1–20.

