## Supplemental Material for "Synteny-based genome assembly for 16 species of *Heliconius* butterflies, and an assessment of structural variation across the genus"

**Table of Contents:**

Supplemental Tables Legends

Supplemental Figures

### Supplemental Tables

The supplemental tables are included in the online Supplemental Material as a single .xlsx file. Each supplemental table legend is provided below.

**Supplemental Table S1** – Samples information for the reference-guided assemblies.

**Supplemental Table S2** – Assembly Statistics.

**Supplemental Table S3** – Reference-guided assemblies chromosome lengths.

**Supplemental Table S4** – BUSCO results for each of the reference-guided assemblies mapped to *H. melpomene* and *H. erato*. Column codes represent: complete single-copy BUSCOs (S); complete duplicated BUSCOs (D); fragmented BUSCOs (F); and missing BUSCOs (M).

**Supplemental Table S5** – Gene annotation results for the reference-guided assemblies mapped to *H. melpomene* and *H. erato*. For each reference-guided assembly, gene annotations were lifted-over from the reference genome used to guide the scaffolding of the *w2rap* assemblies.

**Supplemental Table S6** – Mapping statistics. The original *w2rap* reads were mapped either to the two reference genomes (hmelv25 – *H. melpomene*; heradem – *H. erato demophoon*) or to the new reference-guided assemblies of their own species (MEDUSA-to-hmelv25 – scaffolding using the *H. melpomene* reference; MEDUSA-to-heradem – scaffolded using the *H. erato demophoon* reference).

**Supplemental Table S7** – List of genes in the four repeat-rich regions specific to the *H. hecale*, *H. elevatus* and *H. pardalinus* trio.

**Supplemental Table S8** – List of candidate Inversions.

**Supplemental Table S9** – List of genes within candidate inversions fixed between the two *Heliconius* major clades.

**Supplemental Table S10** – QuIBL results for the triplets (*H. erato – H. melpomene – H. doris*) and (*H. erato – H. melpomene – H. burneyi*). For each trio, the three possible topologies are presented (the last being the species trees). C1,C2: Inferred species tree branch length for the ILS case (C1) and the non-ILS case (C2). The ILS case is forced to be 0, as all lineages must be in the same population. Topology proportions: Inferred mixture proportion for the ILS and non-ILS distributions. These values sum to 1. Trees: Frequency of the topology in the sample. BIC: Raw BIC values for each model. ΔBIC: difference in BIC value between the models. ΔBIC < -10 implies that the ILS+introgression model is a better fit for the data. Species codes: hmel - *H. melpomene*; hera - *H. erato*; hbur - *H. burneyi*; hdor - *H. doris*.

**Supplemental Table S11** – Samples information for the population-based copy number variation inference analysis.

**Supplemental Figures**

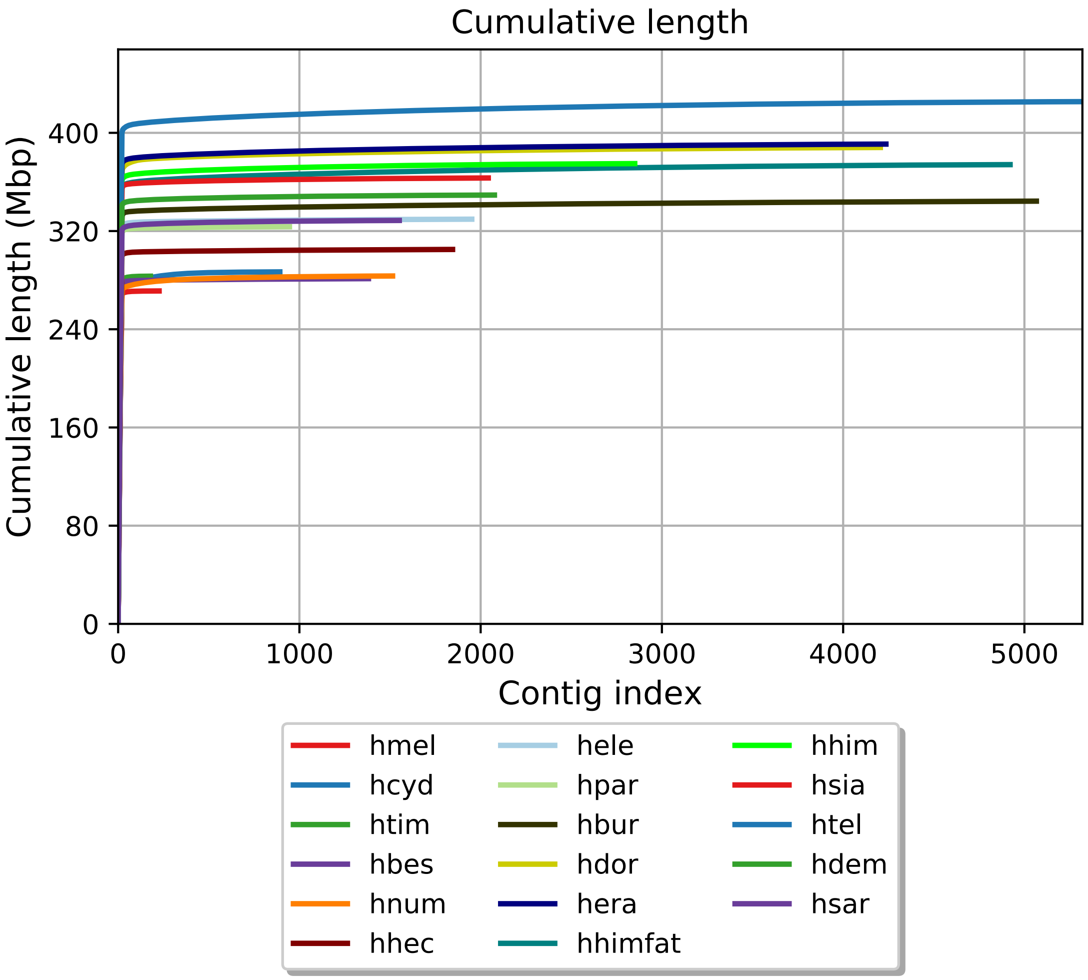


**Supplemental Fig. S1** - Cumulative length distribution of the reference-guided scaffold assemblies scaffolds using *H. melpomene* reference genome. In all the reference-guided assemblies, the cumulative length distribution rapidly plateaus showing that most of the genome is included in the first few super-scaffolds. Species codes are as in Figure 1.

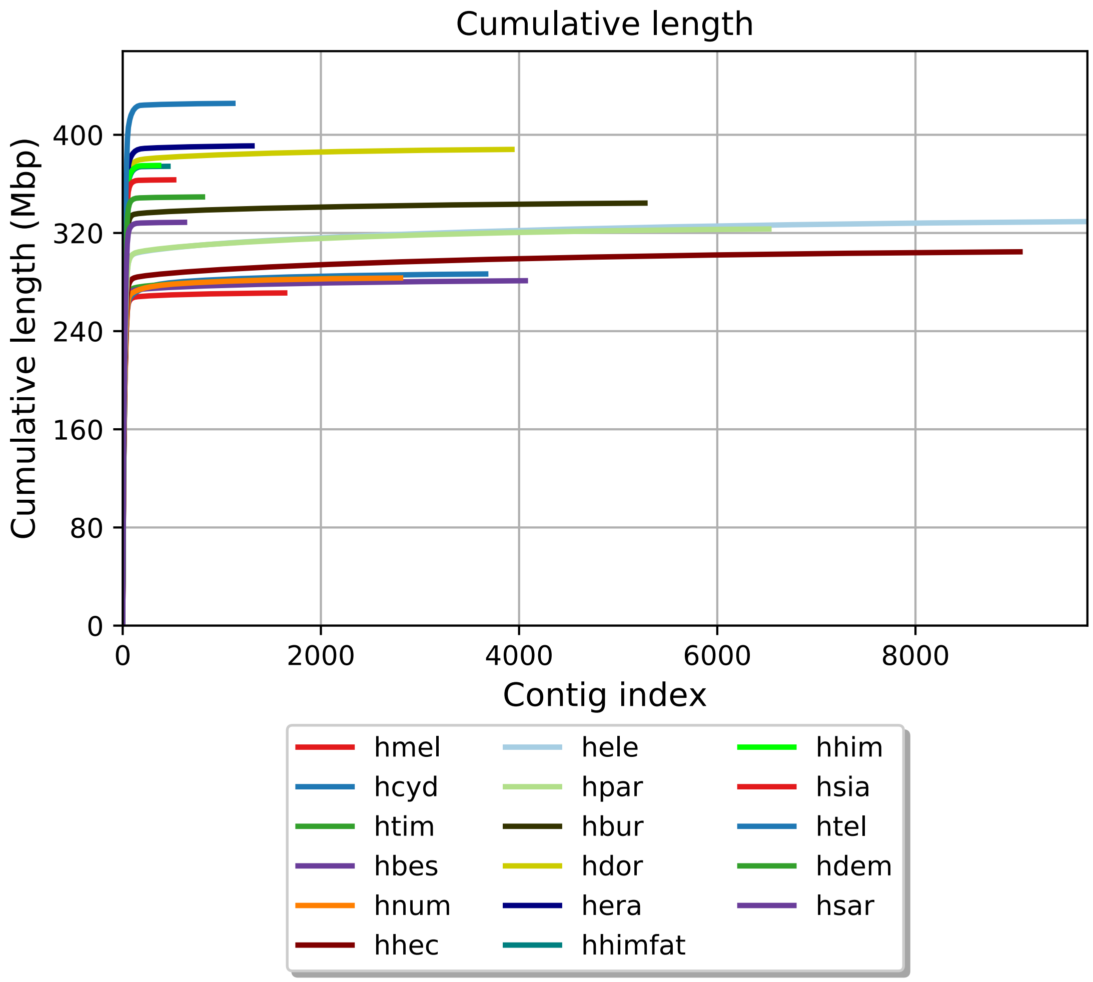


**Supplemental Fig. S2** - Cumulative length distribution of the reference-guided scaffold assemblies scaffolds using *H. erato demophoon* reference genome. In all the reference-guided assemblies, the cumulative length distribution rapidly plateaus showing that most of the genome is included in the first few super-scaffolds. Species codes are as in Figure 1.

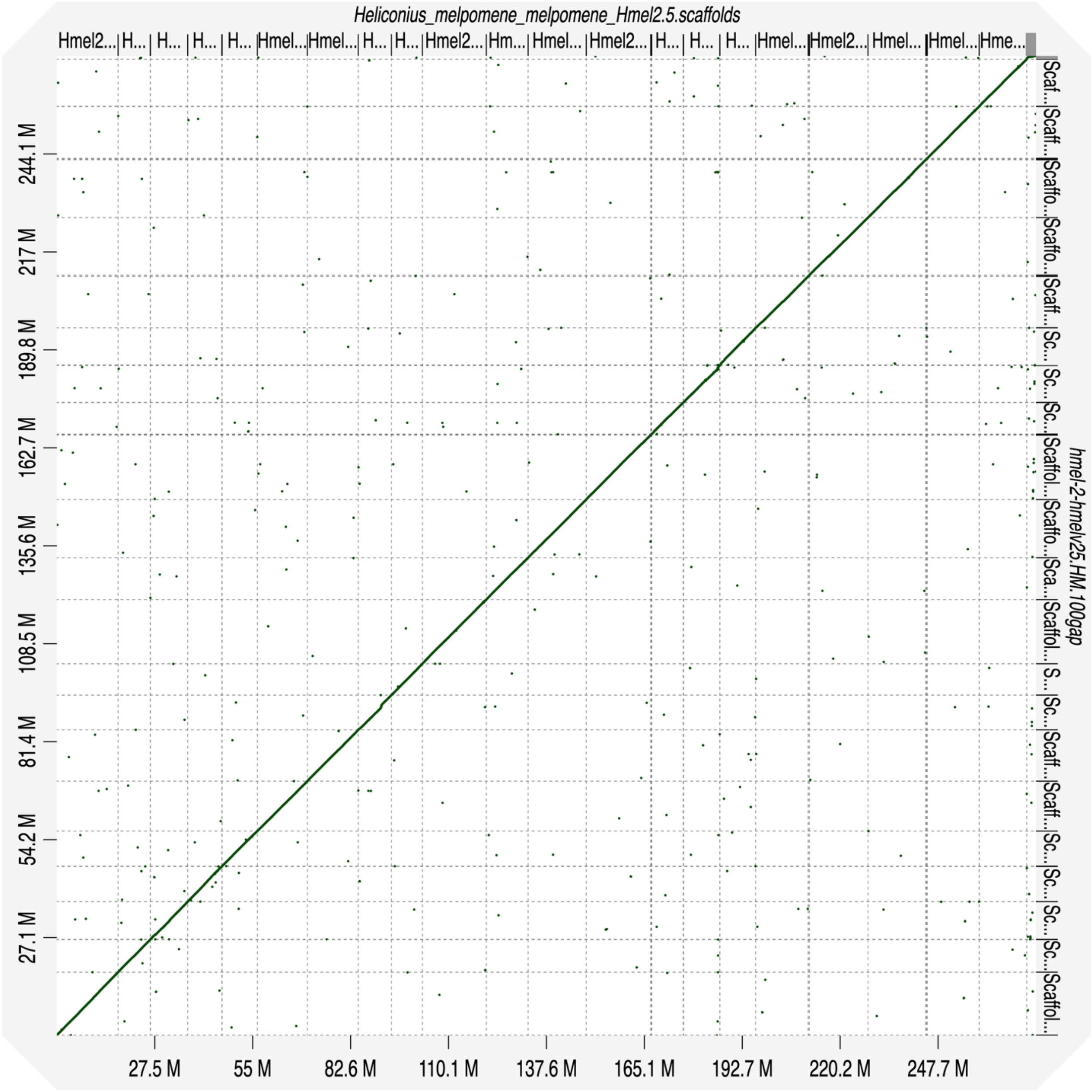


**Supplemental Fig. S3** – Dot plot showing the alignment of the reference-guided *H. melpomene* genome assembly (y-axis) to the *H. melpomene* reference genome (x-axis), used as reference to guide the scaffolding process. *H. melpomene* reference genome scaffolds are ordered, from chromosome 1 to chromosome 21 and including unanchored chromosomes at the end (right).

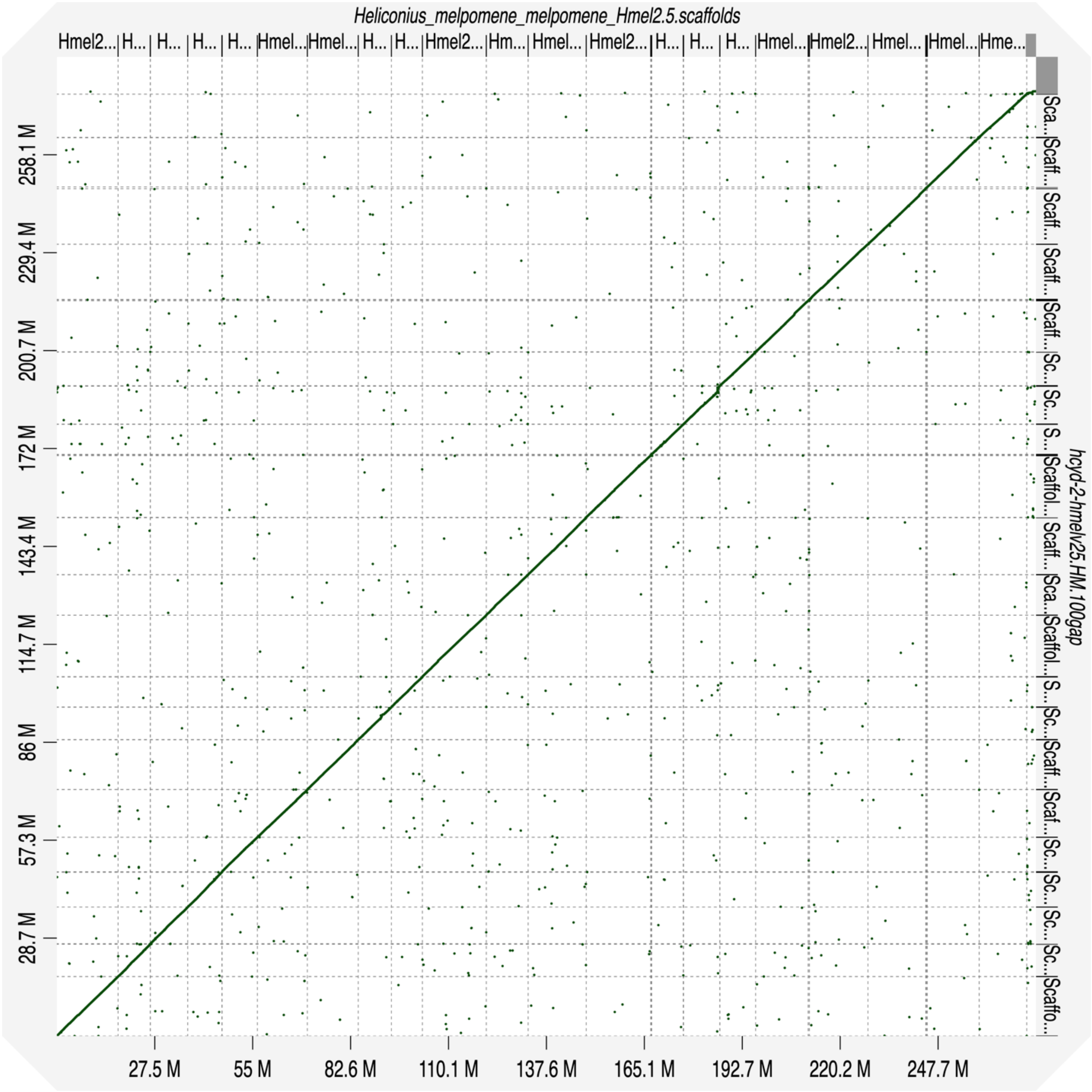


**Supplemental Fig. S4** - Dot plot showing the alignment of the reference-guided *H. cydno* genome assembly (y-axis) to the *H. melpomene* reference genome (x-axis), used as reference to guide the scaffolding process. *H. melpomene* reference genome scaffolds are ordered, from chromosome 1 to chromosome 21 and including unanchored chromosomes at the end (right).

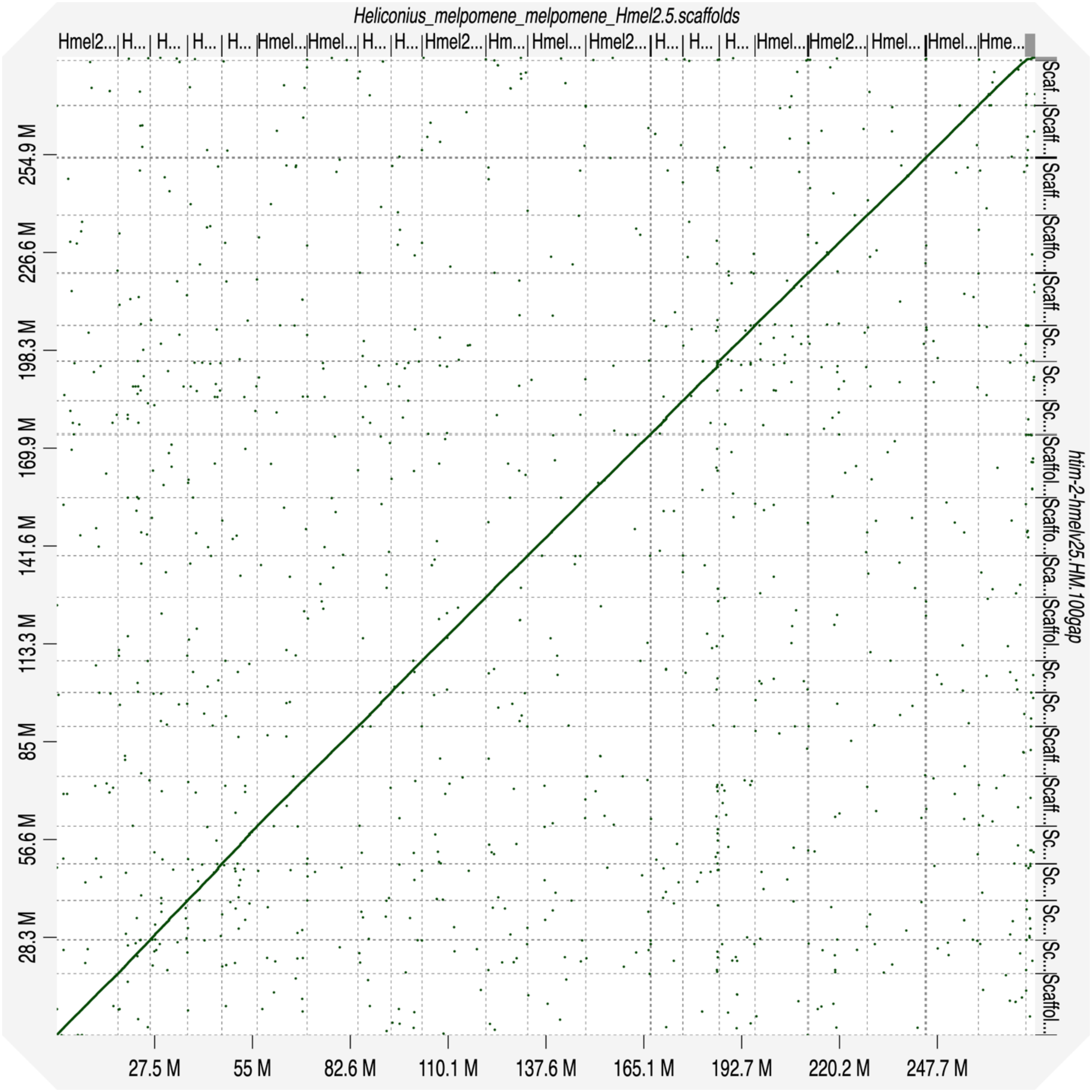


**Supplemental Fig. S5** - Dot plot showing the alignment of the reference-guided *H. timareta* genome assembly (y-axis) to the *H. melpomene* reference genome (x-axis), used as reference to guide the scaffolding process. *H. melpomene* reference genome scaffolds are ordered, from chromosome 1 to chromosome 21 and including unanchored chromosomes at the end (right).

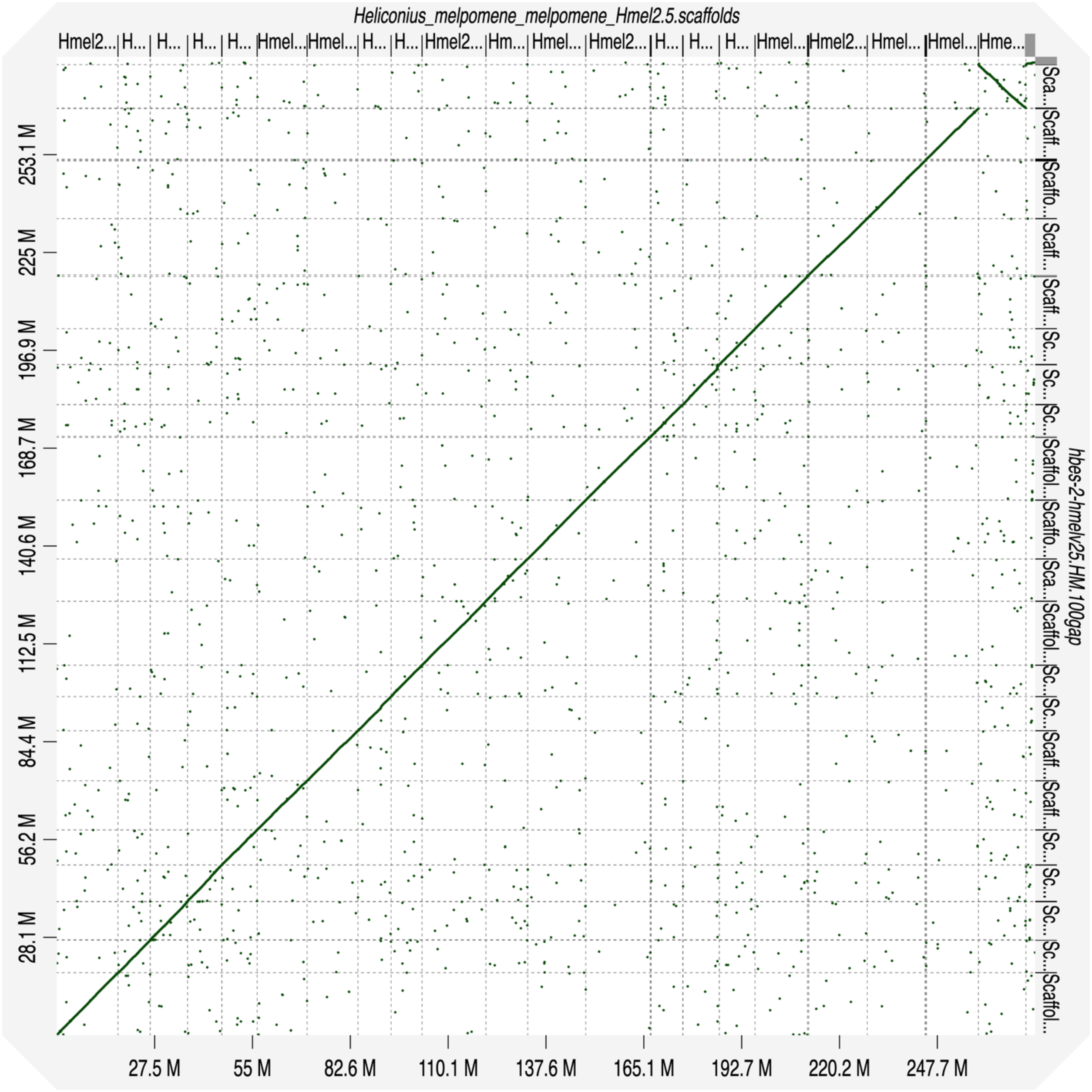


**Supplemental Fig. S6** - Dot plot showing the alignment of the reference-guided *H. besckei* genome assembly (y-axis) to the *H. melpomene* reference genome (x-axis), used as reference to guide the scaffolding process. *H. melpomene* reference genome scaffolds are ordered, from chromosome 1 to chromosome 21 and including unanchored chromosomes at the end (right).

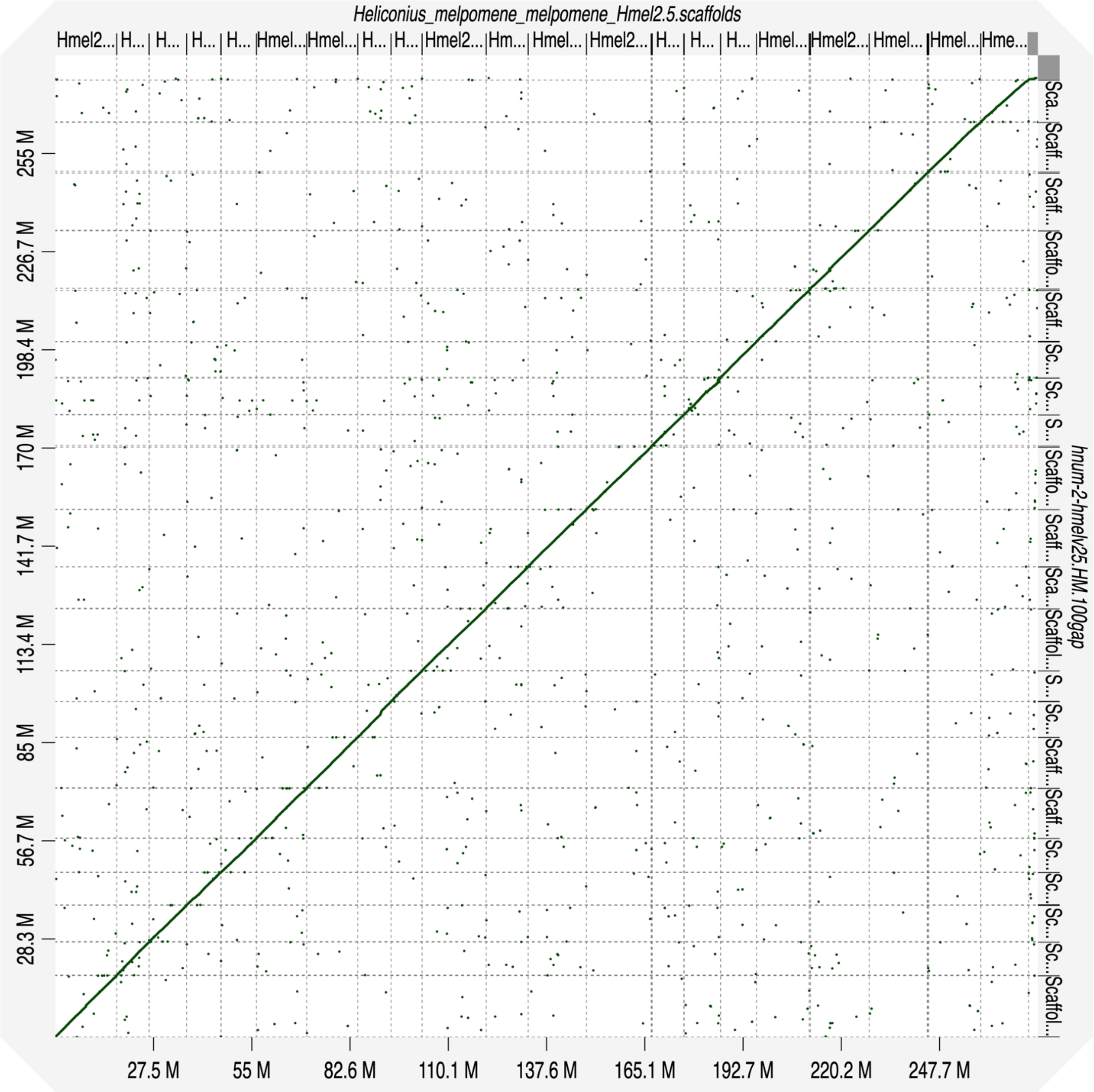


**Supplemental Fig. S7** - Dot plot showing the alignment of the reference-guided *H. numata* genome assembly (y-axis) to the *H. melpomene* reference genome (x-axis) , used as reference to guide the scaffolding process. *H. melpomene* reference genome scaffolds are ordered, from chromosome 1 to chromosome 21 and including unanchored chromosomes at the end (right).

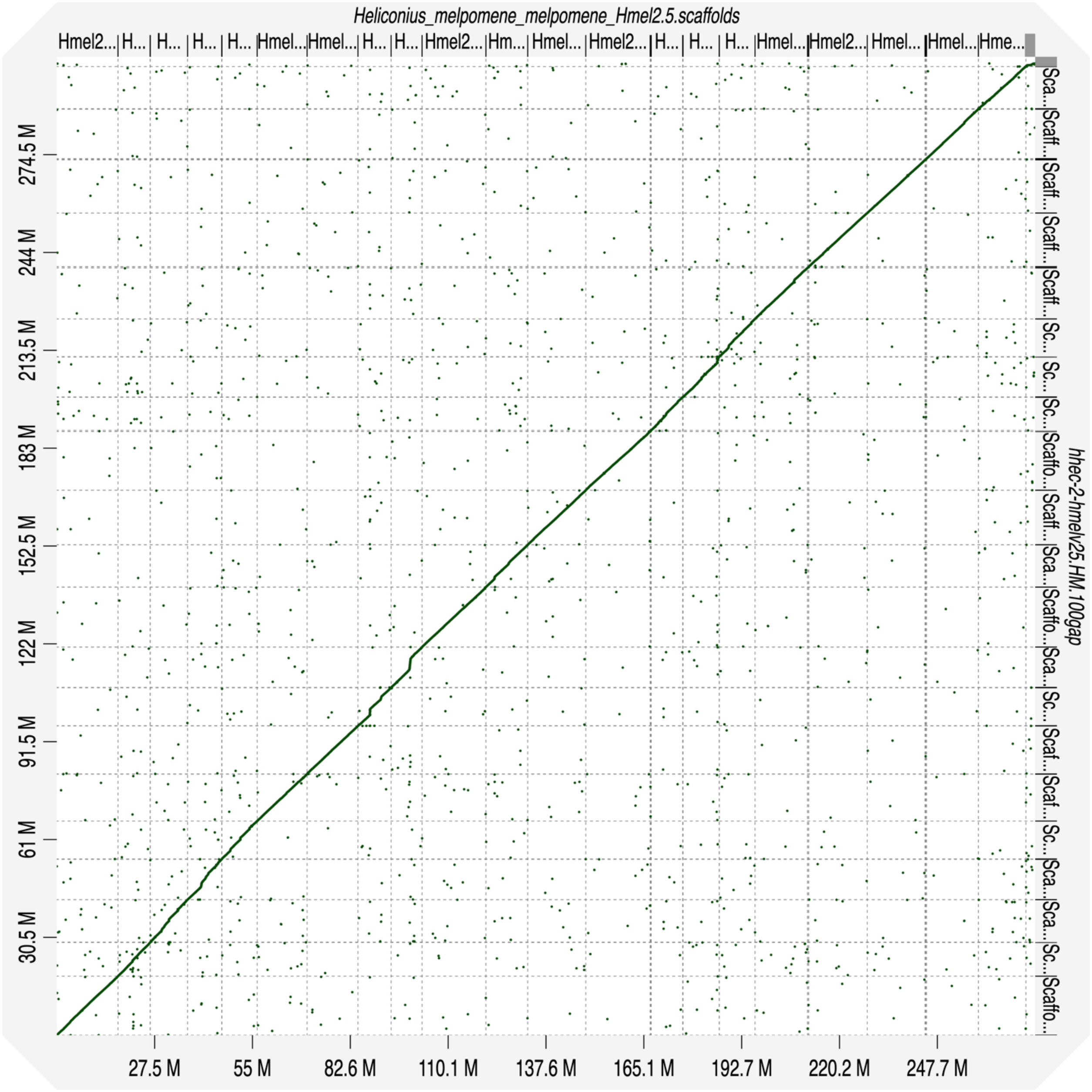


**Supplemental Fig. S8** - Dot plot showing the alignment of the reference-guided *H. hecale* genome assembly (y-axis) to the *H. melpomene* reference genome (x-axis), used as reference to guide the scaffolding process. *H. melpomene* reference genome scaffolds are ordered, from chromosome 1 to chromosome 21 and including unanchored chromosomes at the end (right).

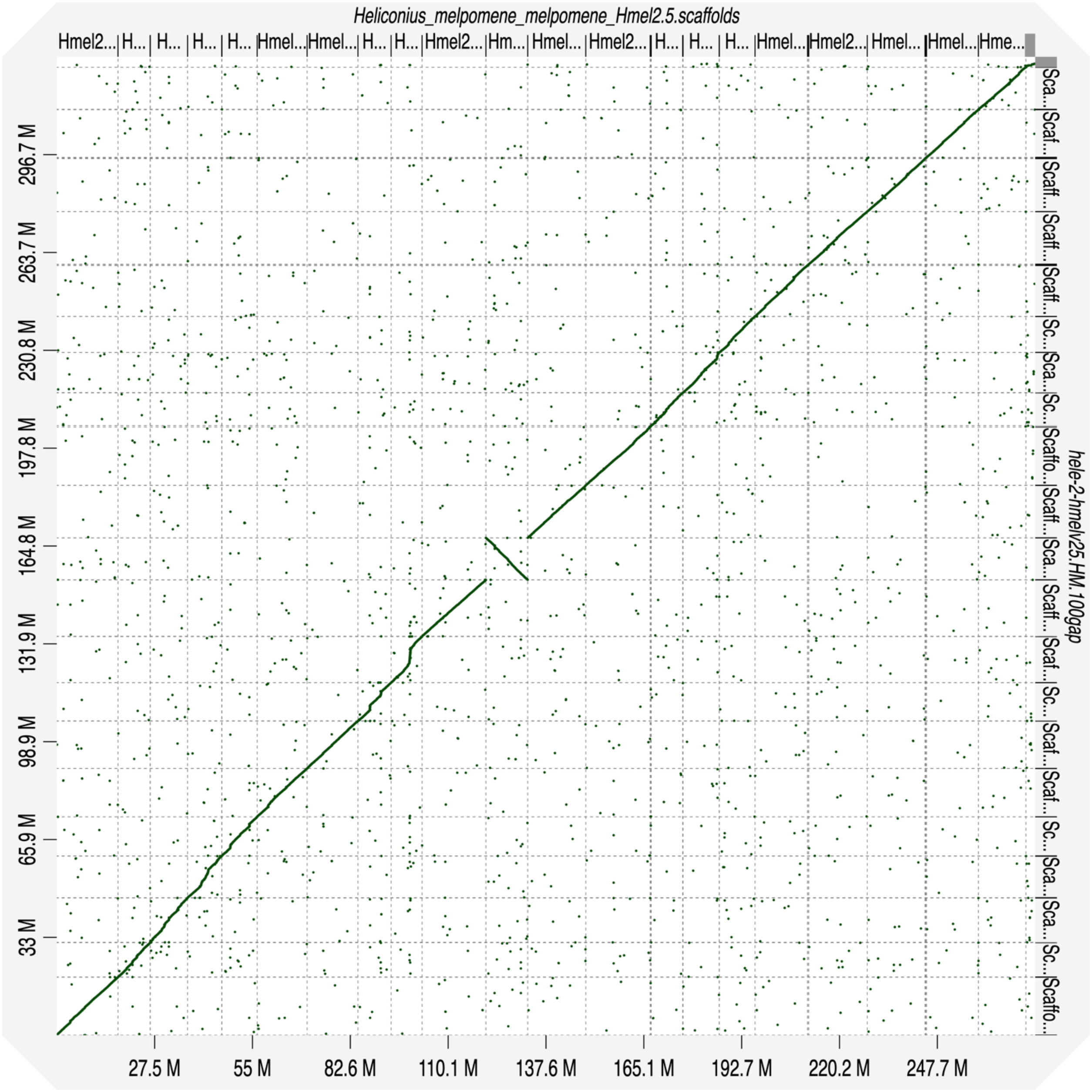


**Supplemental Fig. S9** - Dot plot showing the alignment of the reference-guided *H. elevatus* genome assembly (y-axis) to the *H. melpomene* reference genome (x-axis), used as reference to guide the scaffolding process. *H. melpomene* reference genome scaffolds are ordered, from chromosome 1 to chromosome 21 and including unanchored chromosomes at the end (right).

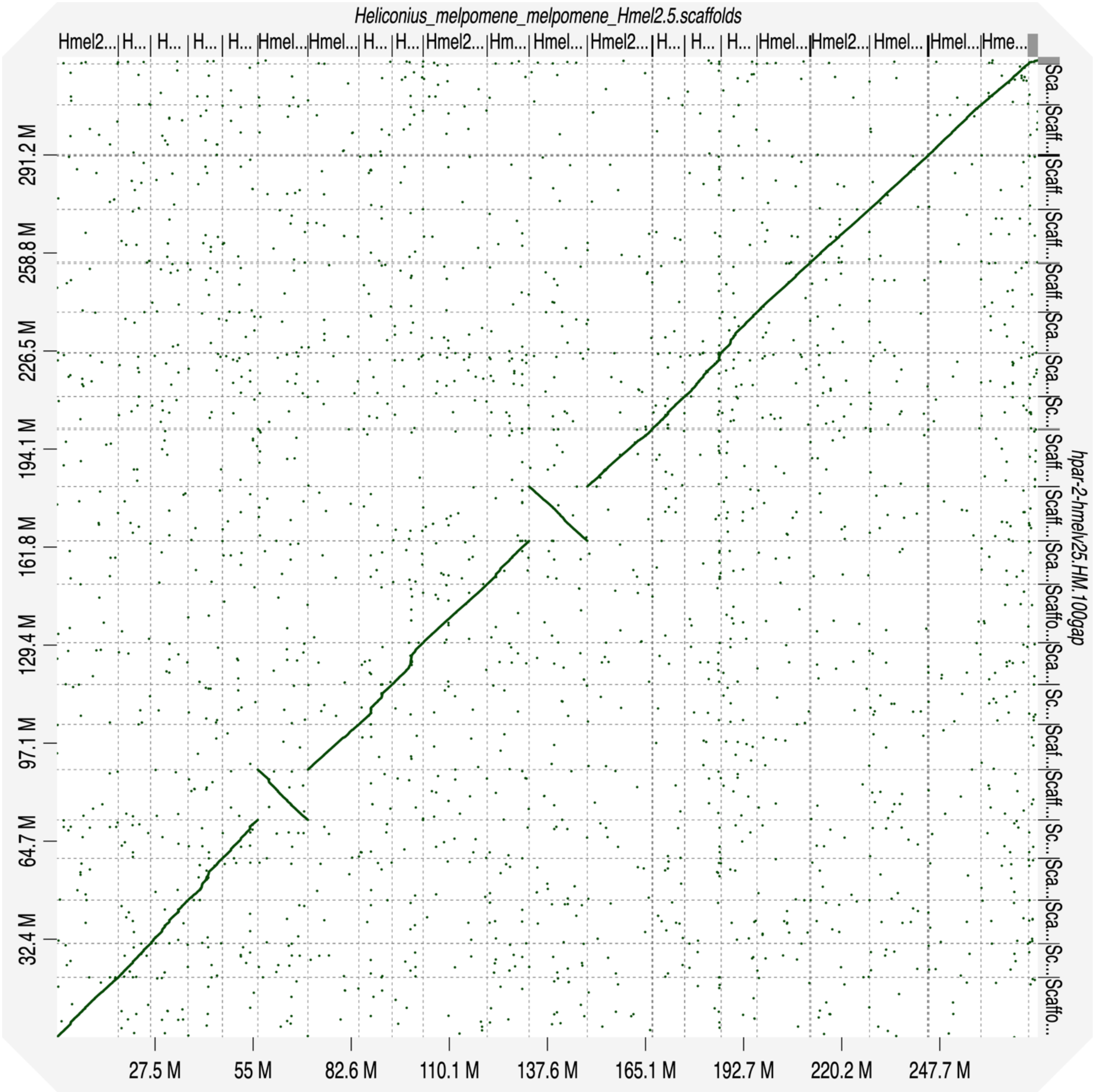


**Supplemental Fig. S10** - Dot plot showing the alignment of the reference-guided *H. pardalinus* genome assembly (y-axis) to the *H. melpomene* reference genome (x-axis), used as reference to guide the scaffolding process. *H. melpomene* reference genome scaffolds are ordered, from chromosome 1 to chromosome 21 and including unanchored chromosomes at the end (right).

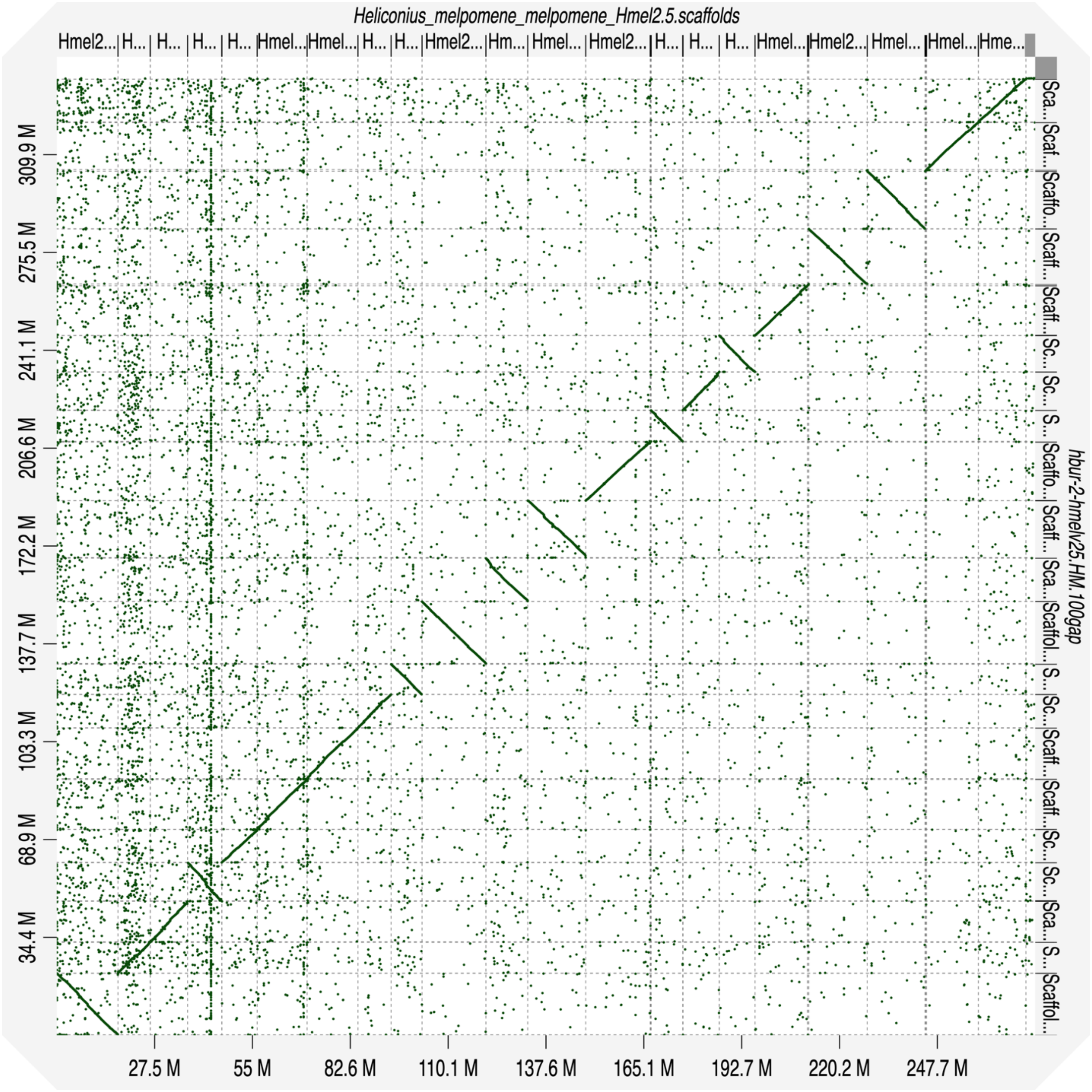


**Supplemental Fig. S11** - Dot plot showing the alignment of the reference-guided *H. burneyi* genome assembly (y-axis) to the *H. melpomene* reference genome (x-axis), used as reference to guide the scaffolding process. *H. melpomene* reference genome scaffolds are ordered, from chromosome 1 to chromosome 21 and including unanchored chromosomes at the end (right).

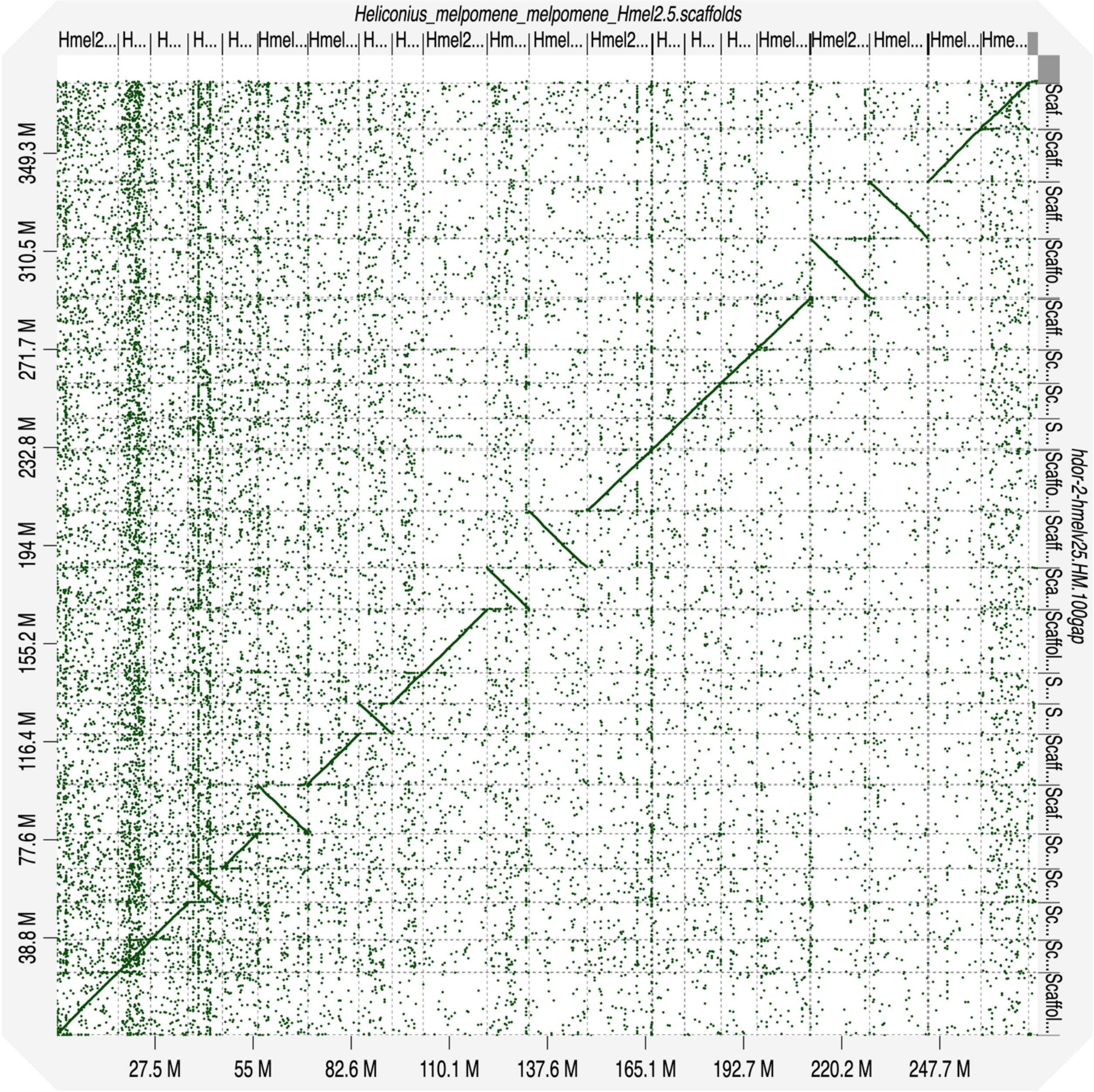


**Supplemental Fig. S12** - Dot plot showing the alignment of the reference-guided *H. doris* genome assembly (y-axis) to the *H. melpomene* reference genome (x-axis), used as reference to guide the scaffolding process. *H. melpomene* reference genome scaffolds are ordered, from chromosome 1 to chromosome 21 and including unanchored chromosomes at the end (right).

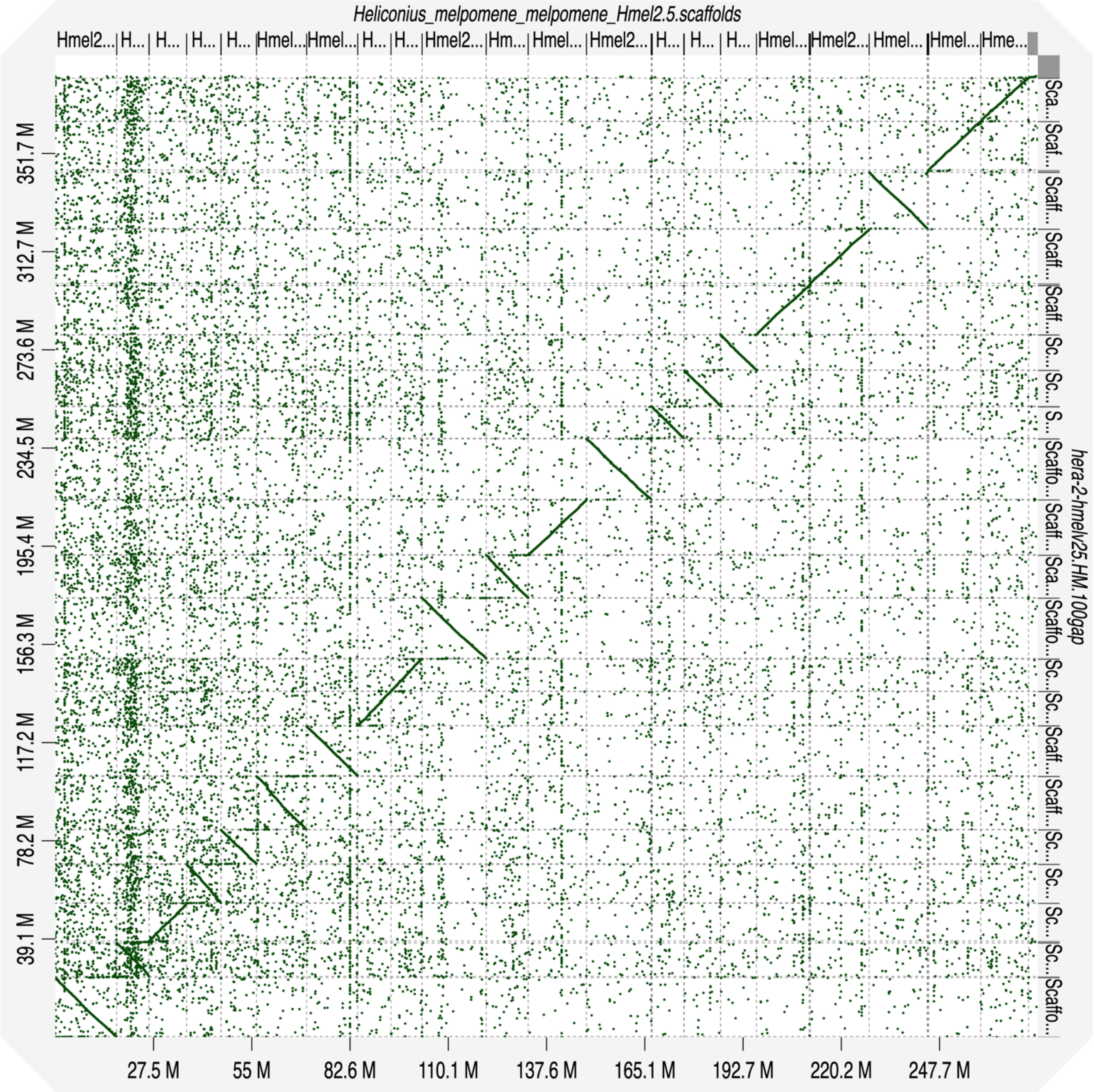


**Supplemental Fig. S13** - Dot plot showing the alignment of the reference-guided *H. erato* genome assembly (y-axis) to the *H. melpomene* reference genome (x-axis), used as reference to guide the scaffolding process. *H. melpomene* reference genome scaffolds are ordered, from chromosome 1 to chromosome 21 and including unanchored chromosomes at the end (right).

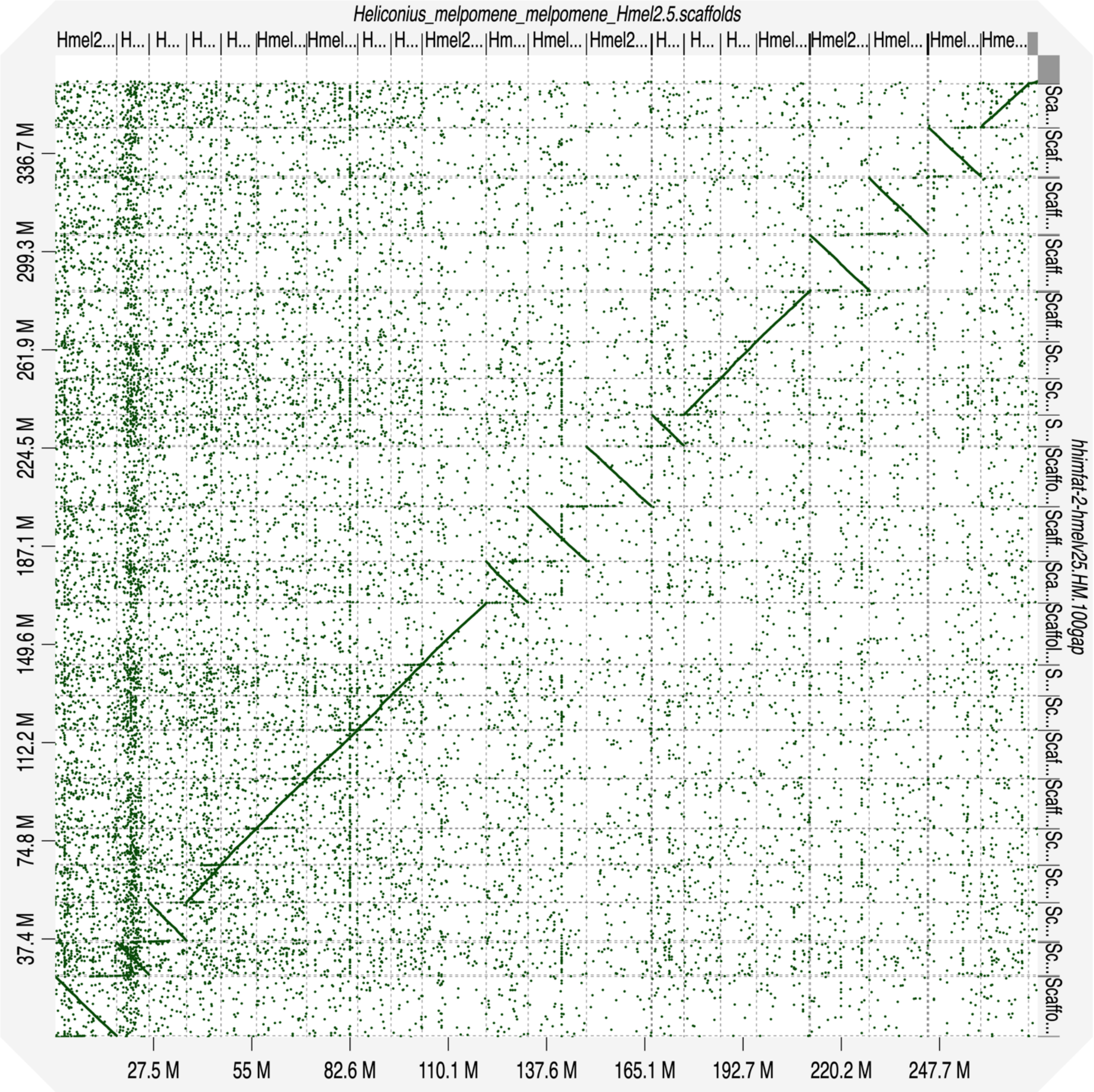


**Supplemental Fig. S14** - - Dot plot showing the alignment of the reference-guided *H. himera* (father) genome assembly (y-axis) to the *H. melpomene* reference genome (x-axis), used as reference to guide the scaffolding process. *H. melpomene* reference genome scaffolds are ordered, from chromosome 1 to chromosome 21 and including unanchored chromosomes at the end (right).

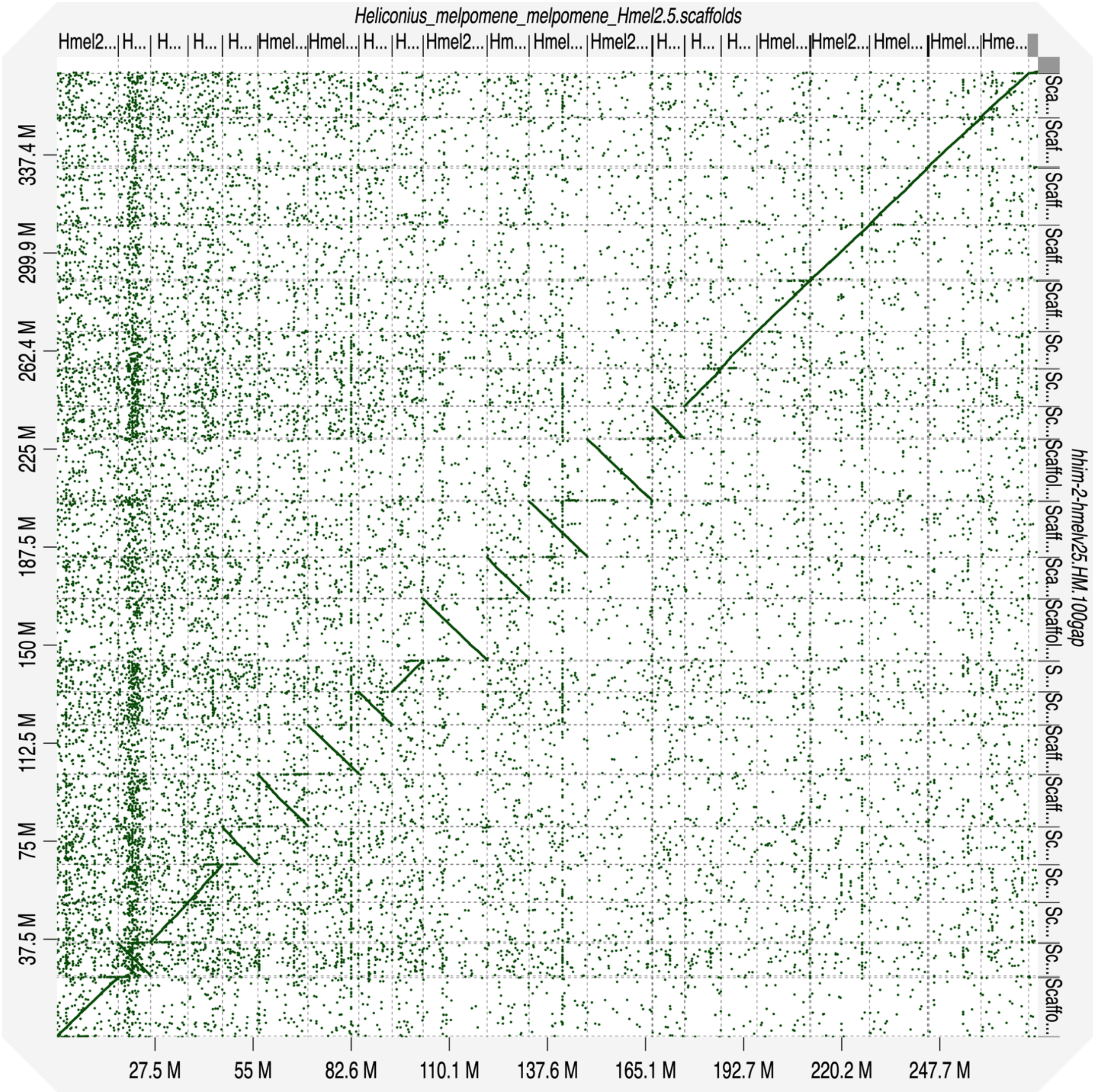


**Supplemental Fig. S15** - Dot plot showing the alignment of the reference-guided *H. himera* genome assembly (y-axis) to the *H. melpomene* reference genome (x-axis), used as reference to guide the scaffolding process. *H. melpomene* reference genome scaffolds are ordered, from chromosome 1 to chromosome 21 and including unanchored chromosomes at the end (right).

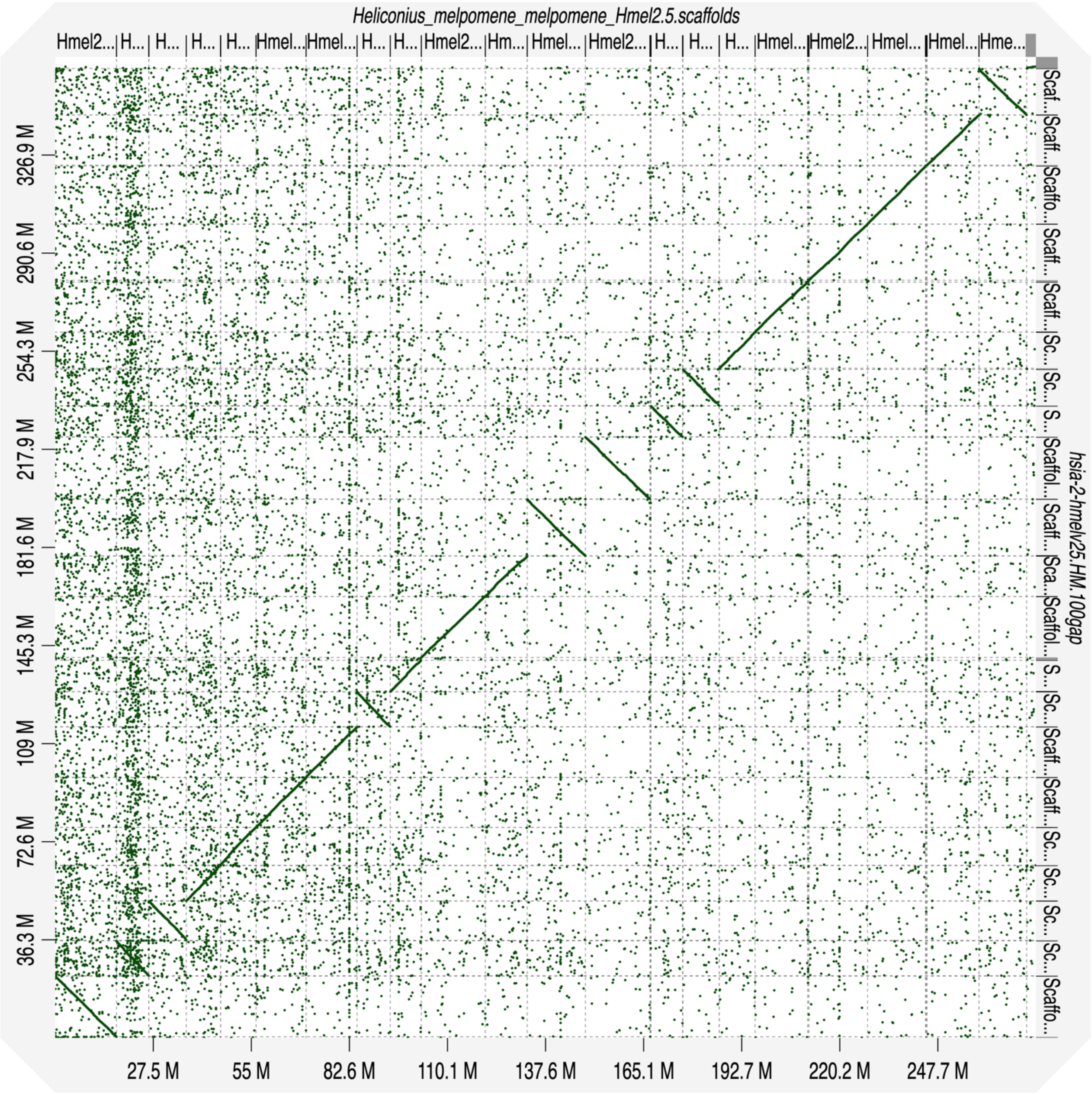


**Supplemental Fig. S16** - Dot plot showing the alignment of the reference-guided *H. hecalesia* genome assembly (y-axis) to the *H. melpomene* reference genome (x-axis), used as reference to guide the scaffolding process. *H. melpomene* reference genome scaffolds are ordered, from chromosome 1 to chromosome 21 and including unanchored chromosomes at the end (right).

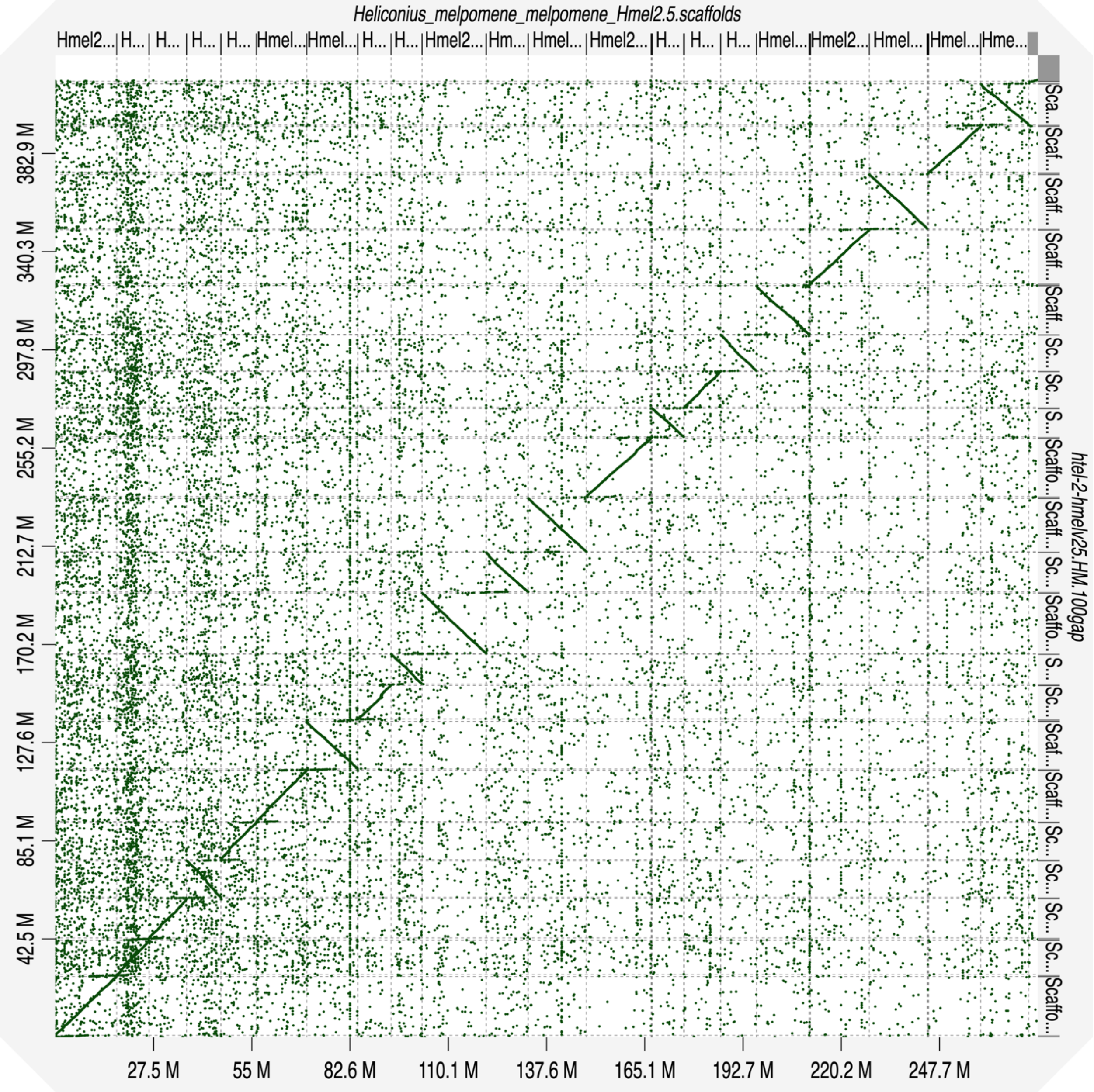


**Supplemental Fig. S17** - Dot plot showing the alignment of the reference-guided *H. telesiphe* genome assembly (y-axis) to the *H. melpomene* reference genome (x-axis), used as reference to guide the scaffolding process. *H. melpomene* reference genome scaffolds are ordered, from chromosome 1 to chromosome 21 and including unanchored chromosomes at the end (right).

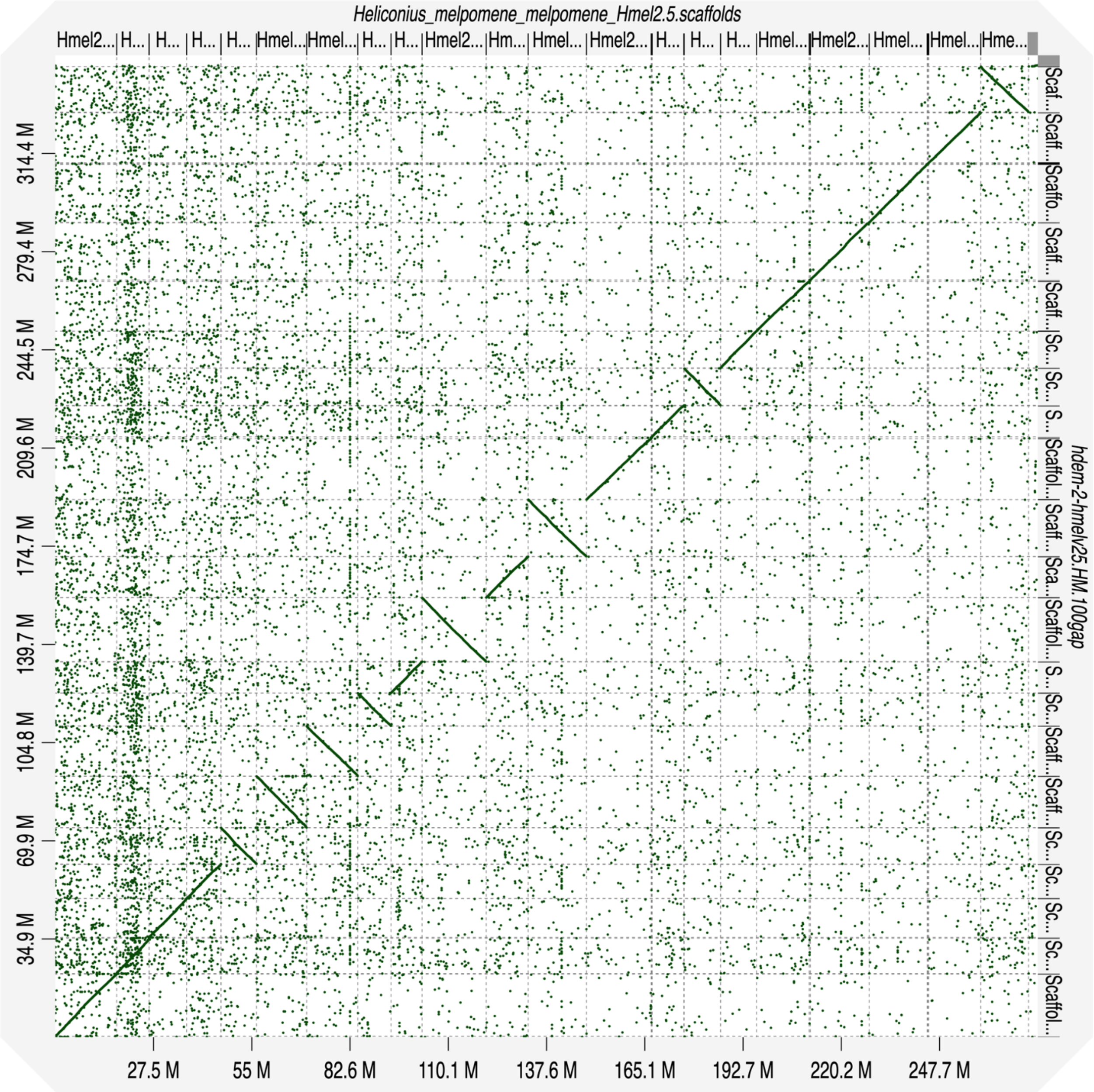


**Supplemental Fig. S18** - Dot plot showing the alignment of the reference-guided *H. demeter* genome assembly (y-axis) to the *H. melpomene* reference genome (x-axis), used as reference to guide the scaffolding process. *H. melpomene* reference genome scaffolds are ordered, from chromosome 1 to chromosome 21 and including unanchored chromosomes at the end (right).

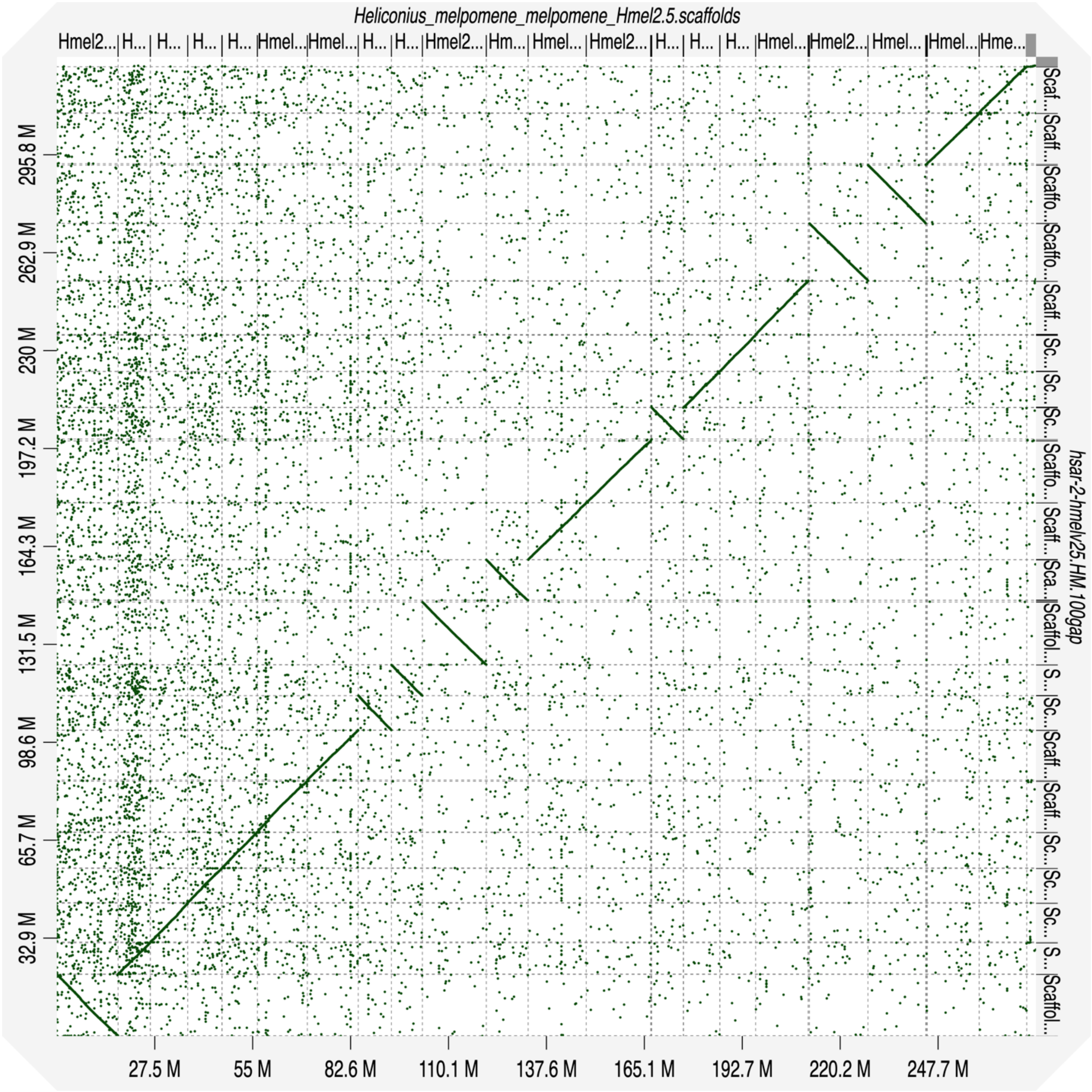


**Supplemental Fig. S19** - Dot plot showing the alignment of the reference-guided *H. sara* genome assembly (y-axis) to the *H. melpomene* reference genome (x-axis), used as reference to guide the scaffolding process. *H. melpomene* reference genome scaffolds are ordered, from chromosome 1 to chromosome 21 and including unanchored chromosomes at the end (right).

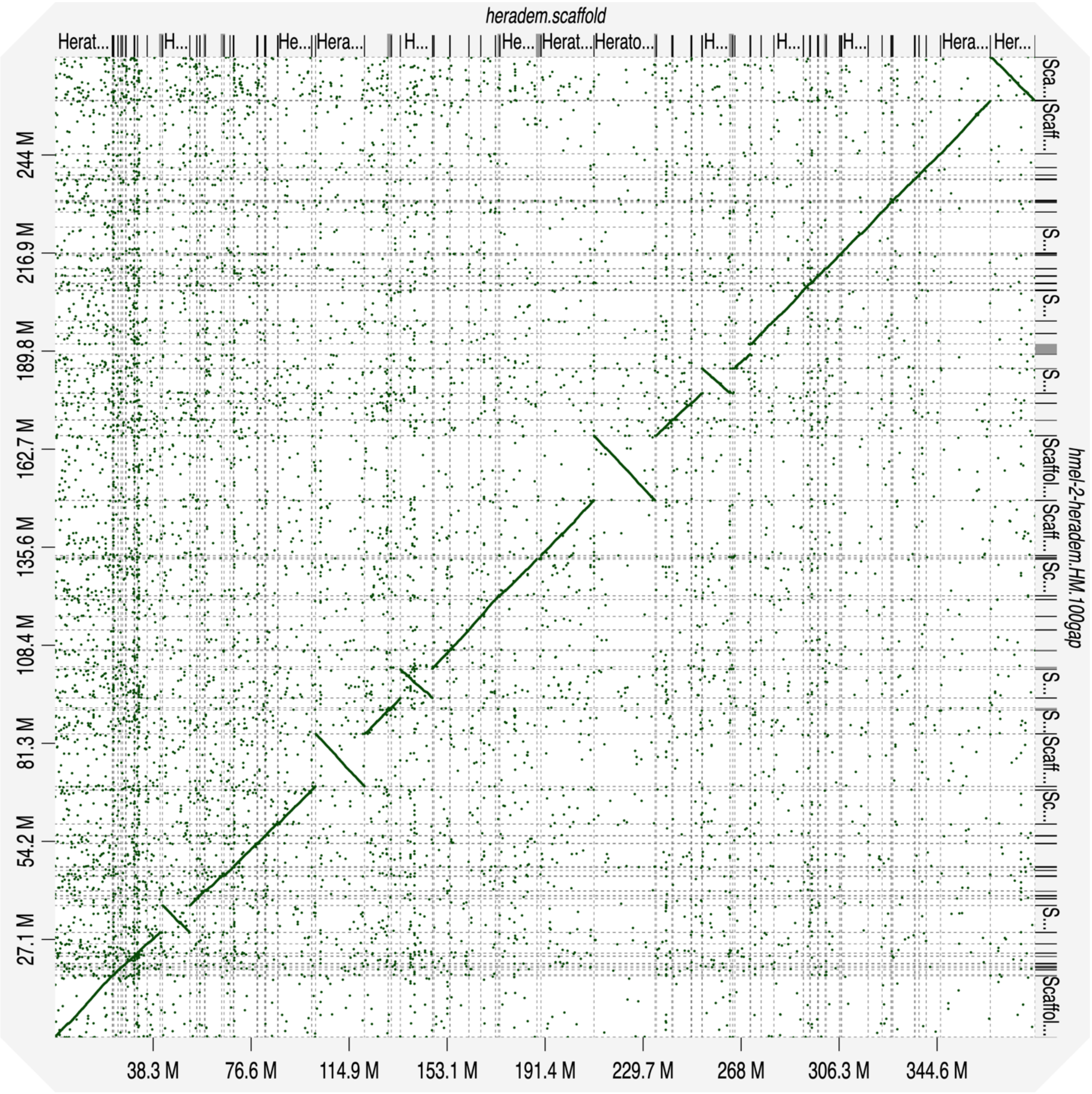


**Supplemental Fig. S20** - Dot plot showing the alignment of the reference-guided *H. melpomene* genome assembly (y-axis) to the *H. erato demophoon* reference genome (x-axis), used as reference to guide the scaffolding process. *H. erato* reference genome scaffolds are ordered, from chromosome 1 to chromosome 21.

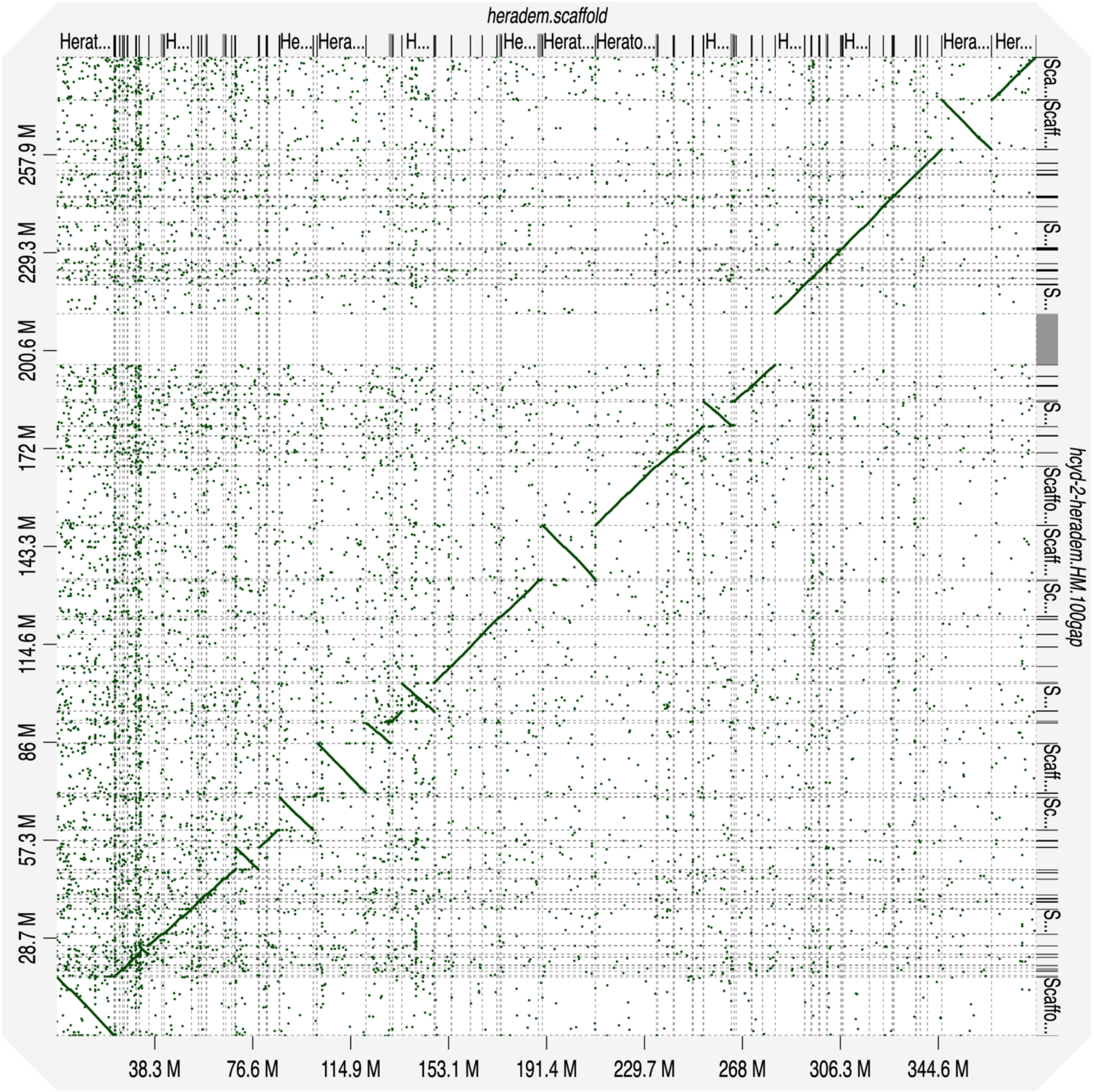


**Supplemental Fig. S21** - Dot plot showing the alignment of the reference-guided *H. cydno* genome assembly (y-axis) to the *H. erato demophoon* reference genome (x-axis), used as reference to guide the scaffolding process. *H. erato* reference genome scaffolds are ordered, from chromosome 1 to chromosome 21.

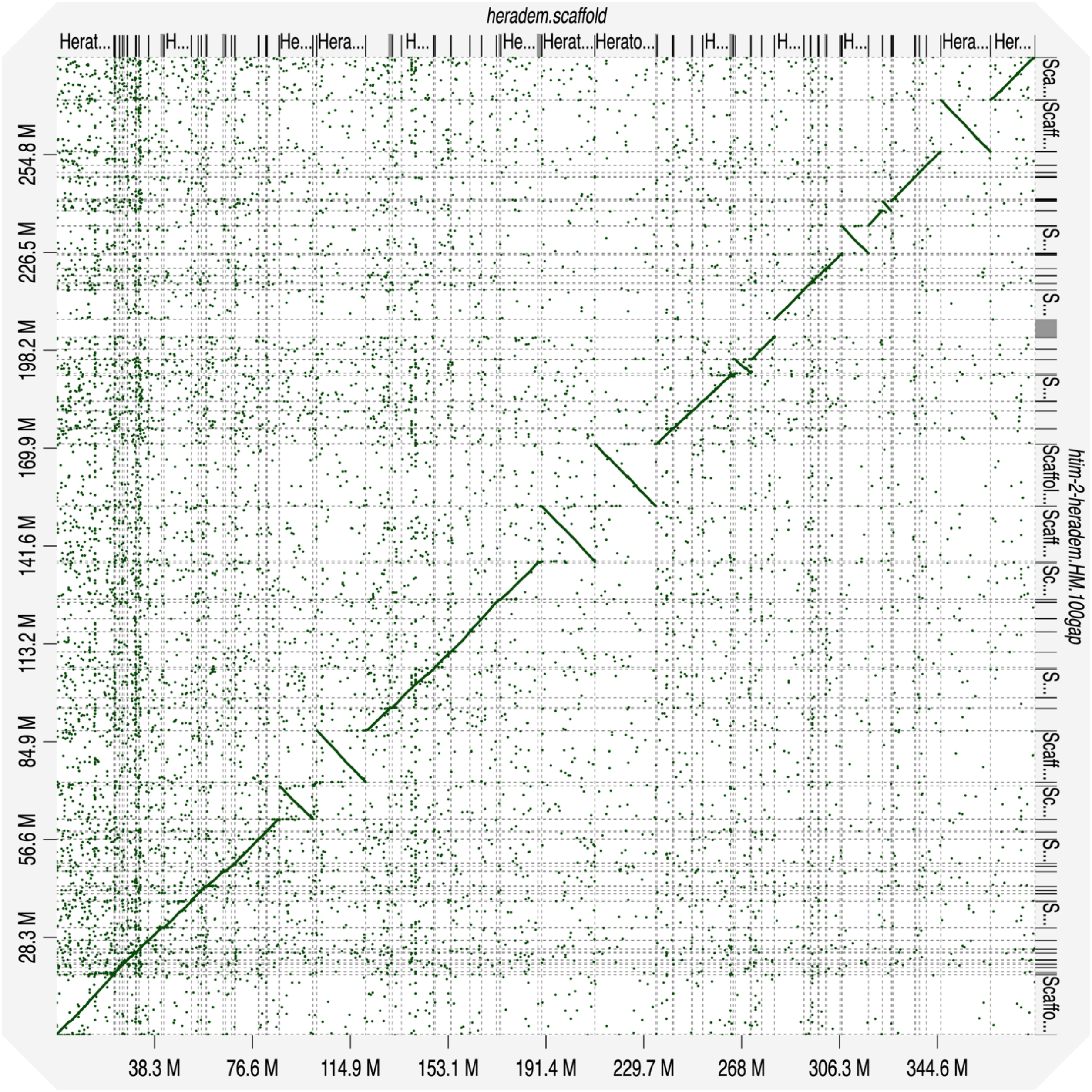


**Supplemental Fig. S22** - Dot plot showing the alignment of the reference-guided *H. timareta* genome assembly (y-axis) to the *H. erato demophoon* reference genome (x-axis), used as reference to guide the scaffolding process. *H. erato* reference genome scaffolds are ordered, from chromosome 1 to chromosome 21.

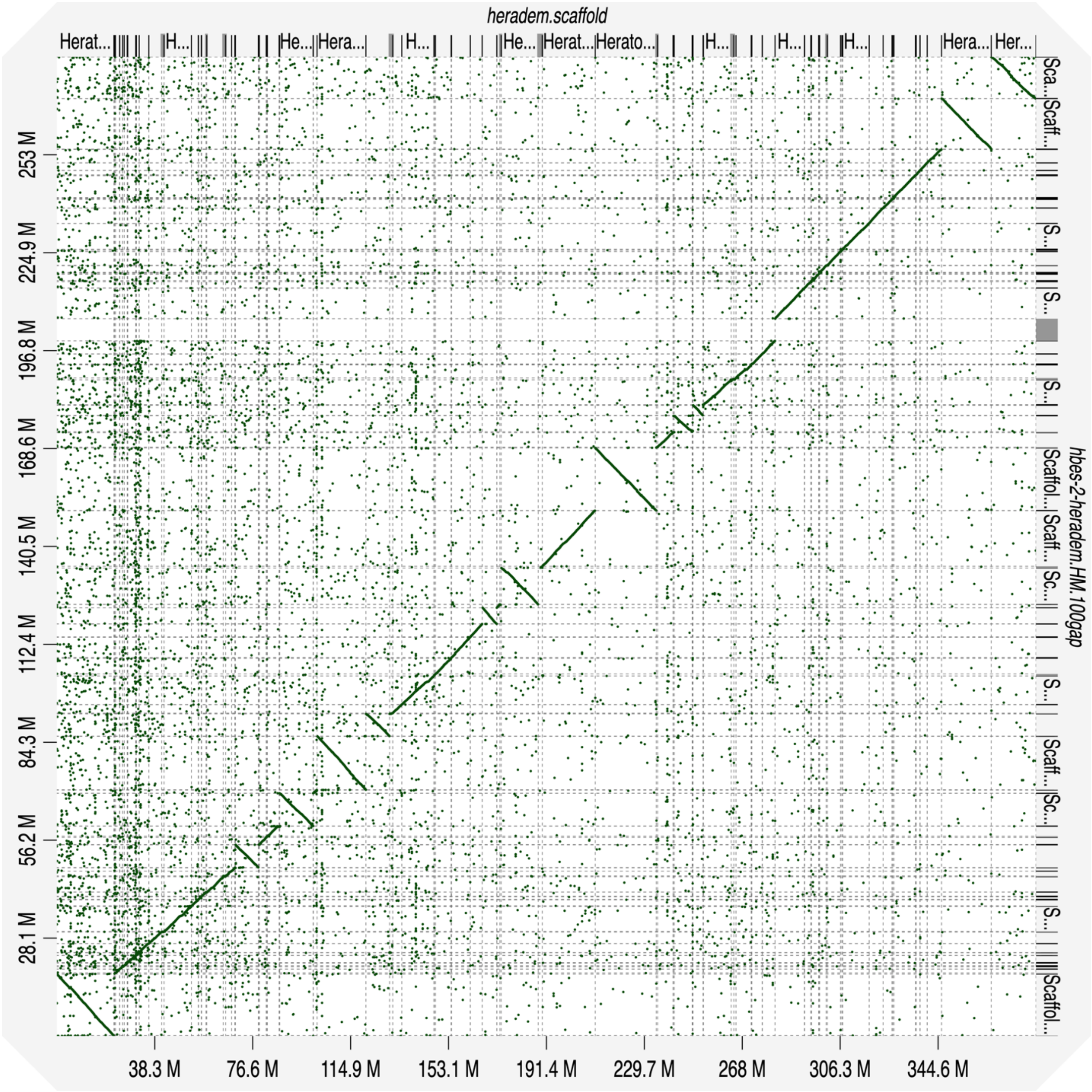


**Supplemental Fig. S23** - Dot plot showing the alignment of the reference-guided *H. besckei* genome assembly (y-axis) to the *H. erato demophoon* reference genome (x-axis), used as reference to guide the scaffolding process. *H. erato* reference genome scaffolds are ordered, from chromosome 1 to chromosome 21.

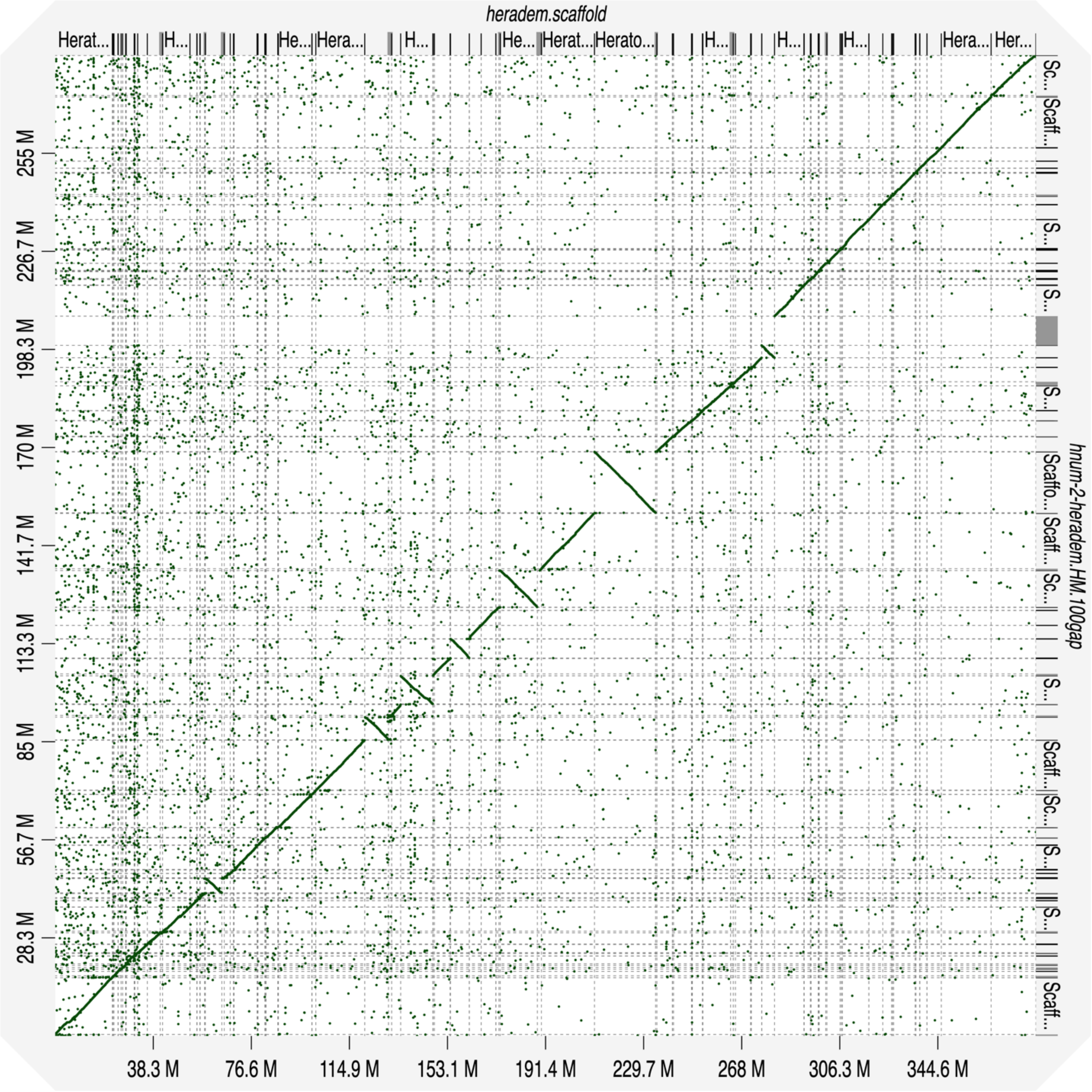


**Supplemental Fig. S24** - Dot plot showing the alignment of the reference-guided *H. numata* genome assembly (y-axis) to the *H. erato demophoon* reference genome (x-axis), used as reference to guide the scaffolding process. *H. erato* reference genome scaffolds are ordered, from chromosome 1 to chromosome 21.

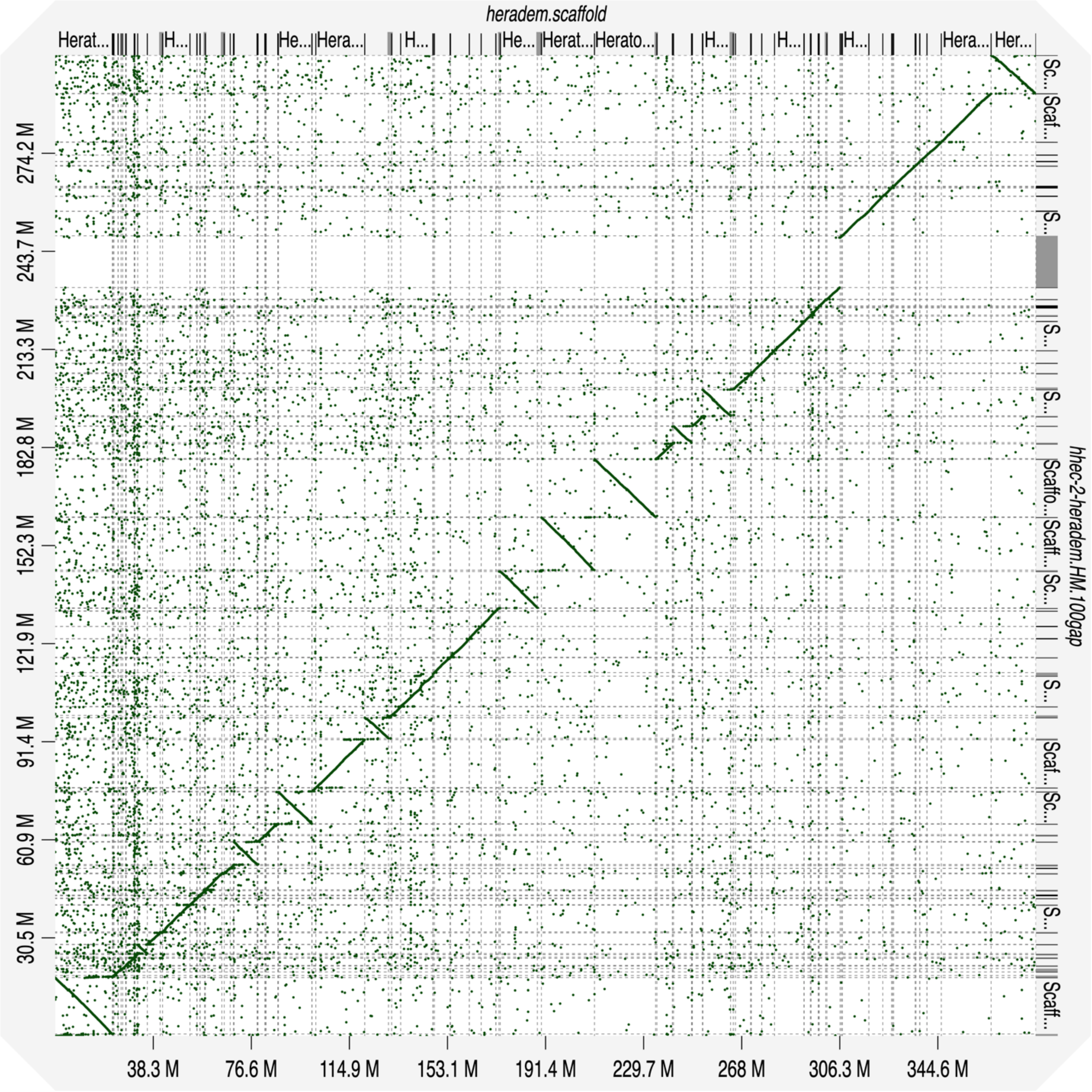


**Supplemental Fig. S25** - Dot plot showing the alignment of the reference-guided *H. hecale* genome assembly (y-axis) to the *H. erato demophoon* reference genome (x-axis), used as reference to guide the scaffolding process. *H. erato* reference genome scaffolds are ordered, from chromosome 1 to chromosome 21.

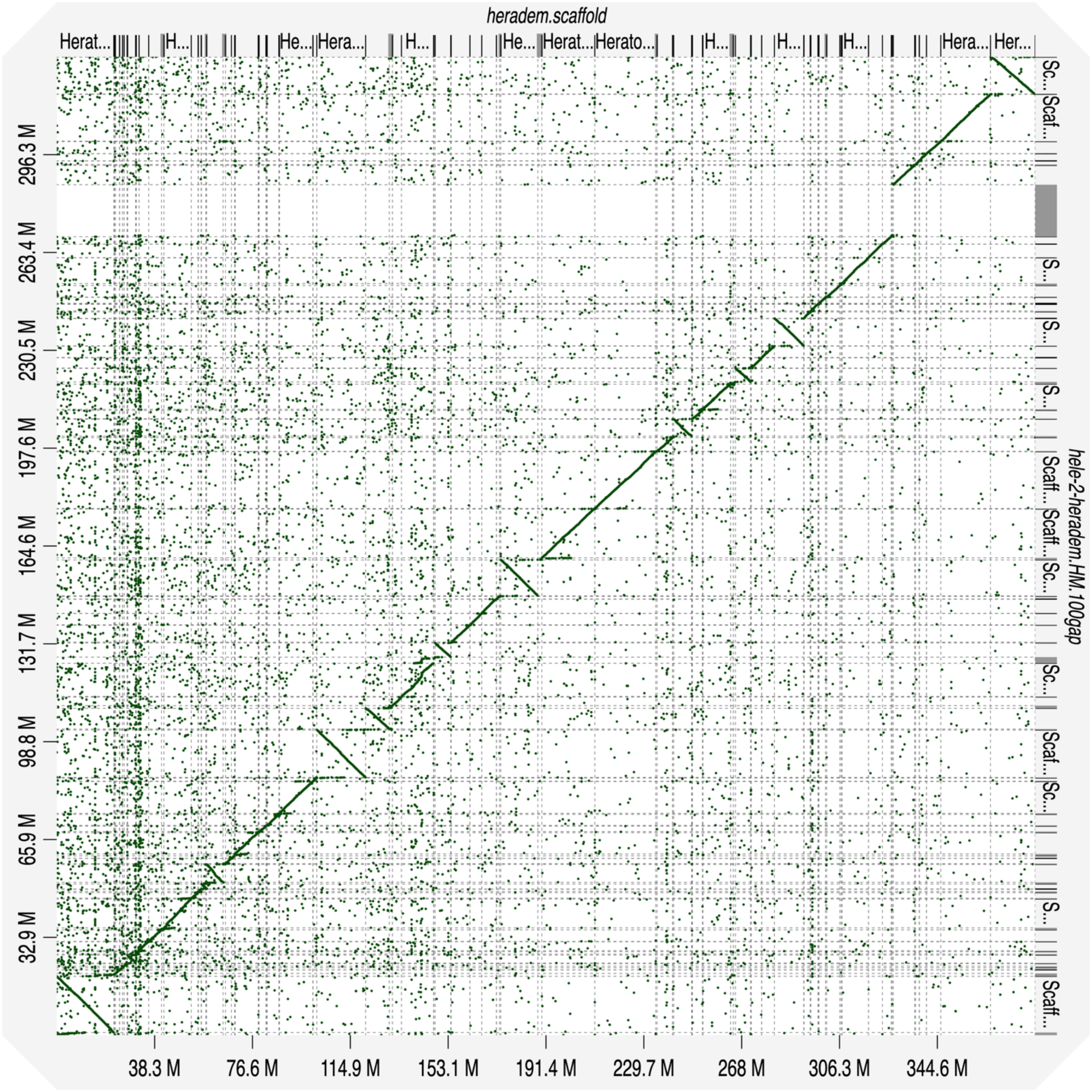


**Supplemental Fig. S26** - Dot plot showing the alignment of the reference-guided *H. elevatus* genome assembly (y-axis) to the *H. erato demophoon* reference genome (x-axis), used as reference to guide the scaffolding process. *H. erato* reference genome scaffolds are ordered, from chromosome 1 to chromosome 21.

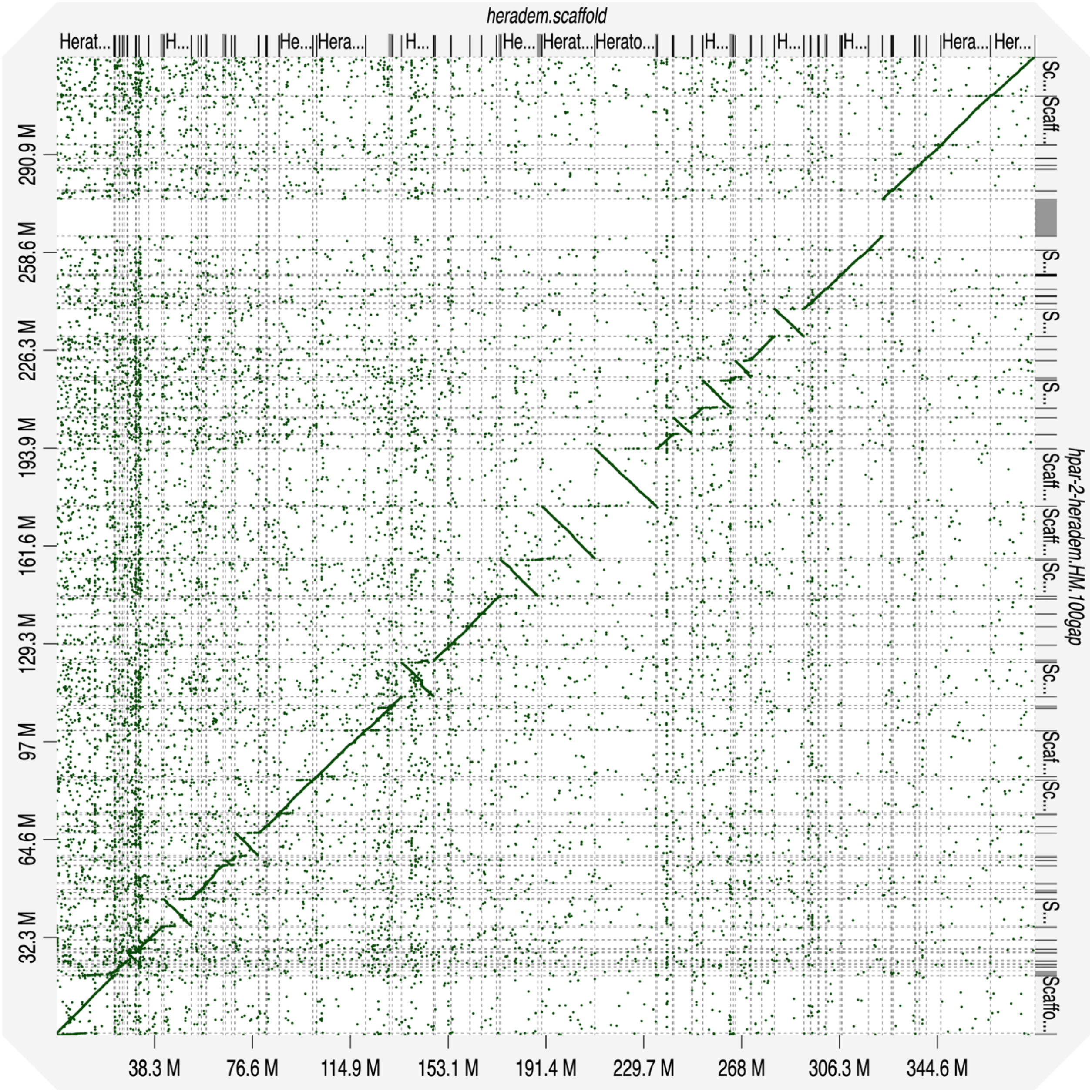


**Supplemental Fig. S27** - Dot plot showing the alignment of the reference-guided *H. pardalinus* genome assembly (y-axis) to the *H. erato demophoon* reference genome (x-axis), used as reference to guide the scaffolding process. *H. erato* reference genome scaffolds are ordered, from chromosome 1 to chromosome 21.

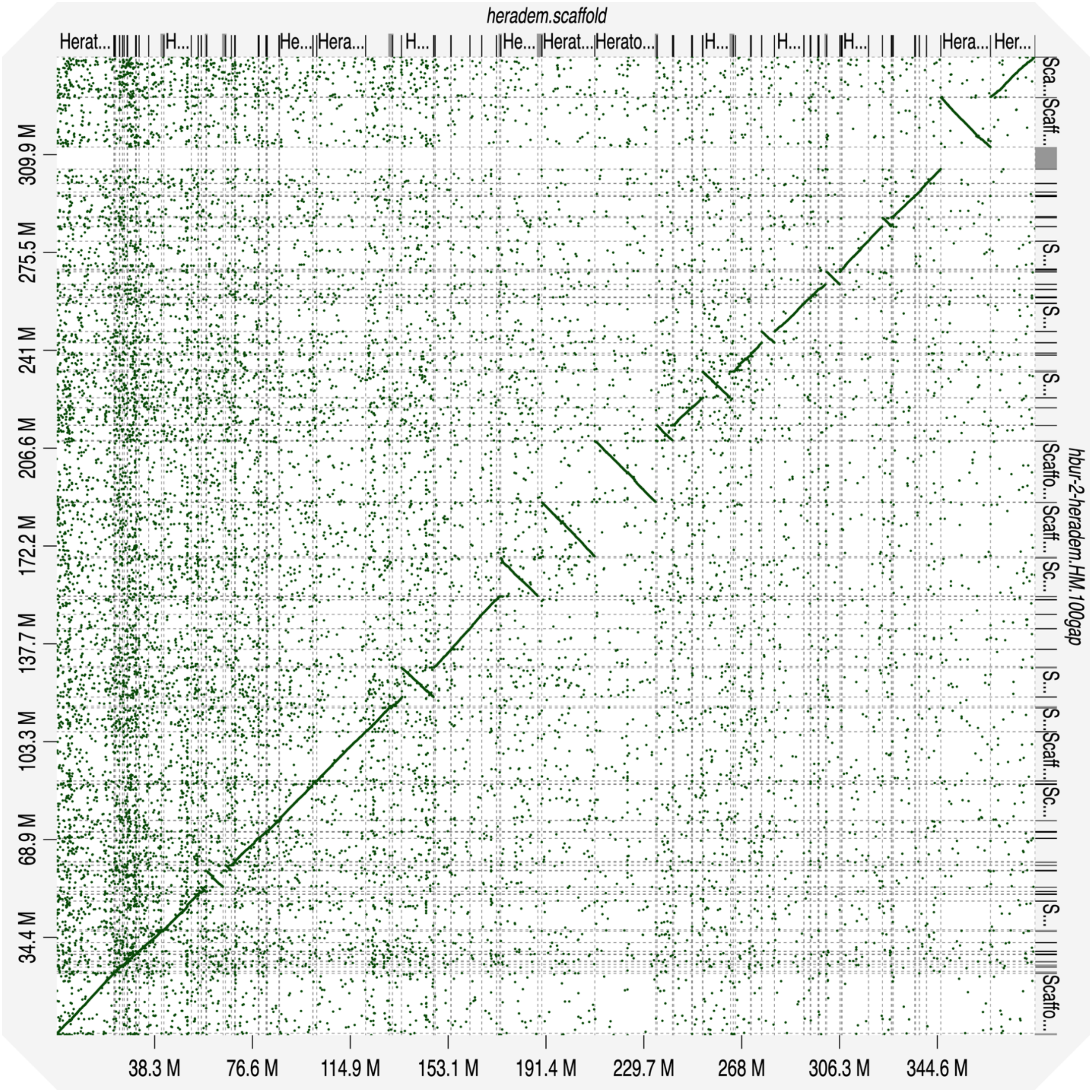


**Supplemental Fig. S28** - Dot plot showing the alignment of the reference-guided *H. burneyi* genome assembly (y-axis) to the *H. erato demophoon* reference genome (x-axis), used as reference to guide the scaffolding process. *H. erato* reference genome scaffolds are ordered, from chromosome 1 to chromosome 21.

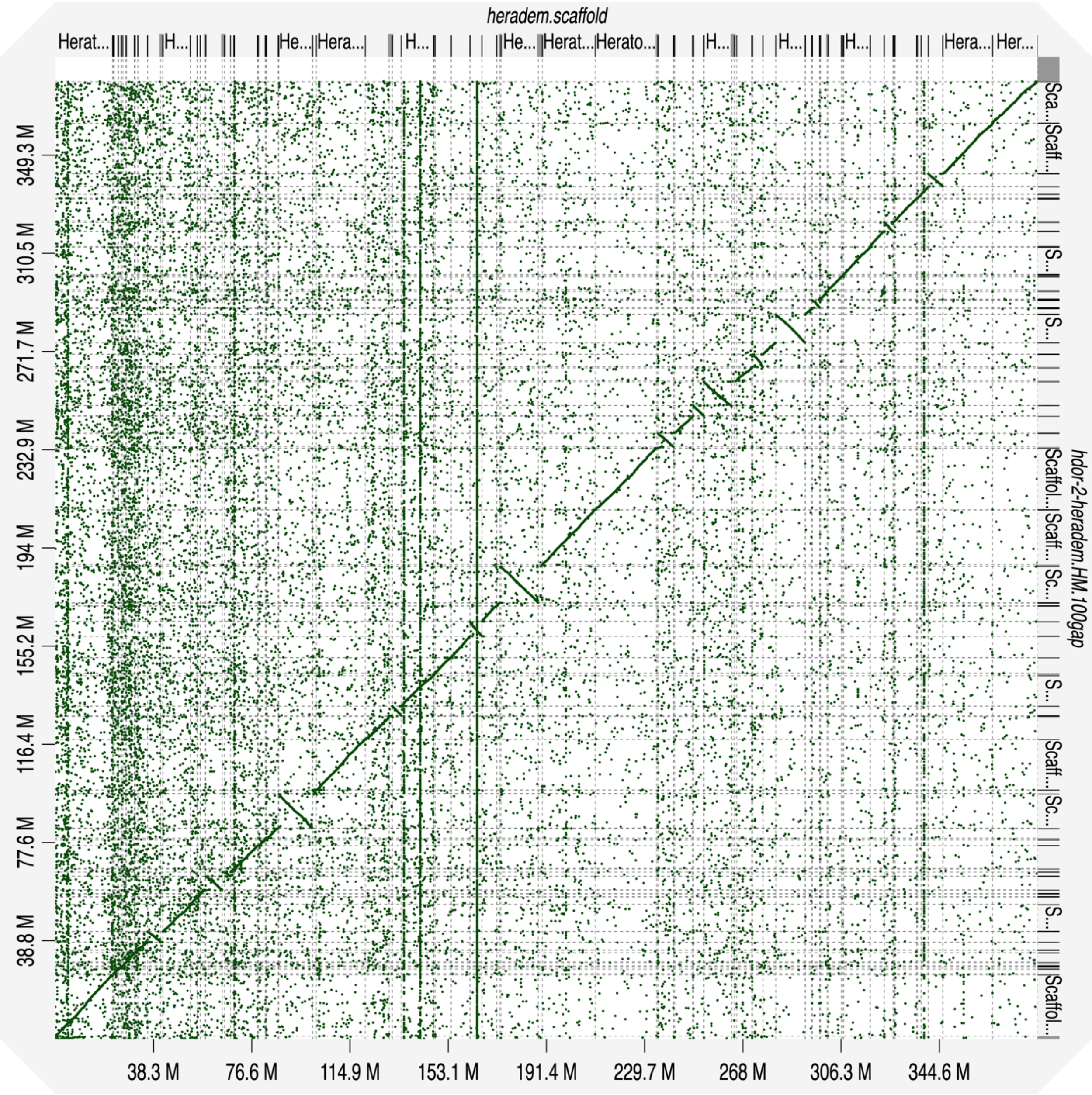


**Supplemental Fig. S29** - Dot plot showing the alignment of the reference-guided *H. doris* genome assembly (y-axis) to the *H. erato demophoon* reference genome (x-axis), used as reference to guide the scaffolding process. *H. erato* reference genome scaffolds are ordered, from chromosome 1 to chromosome 21.

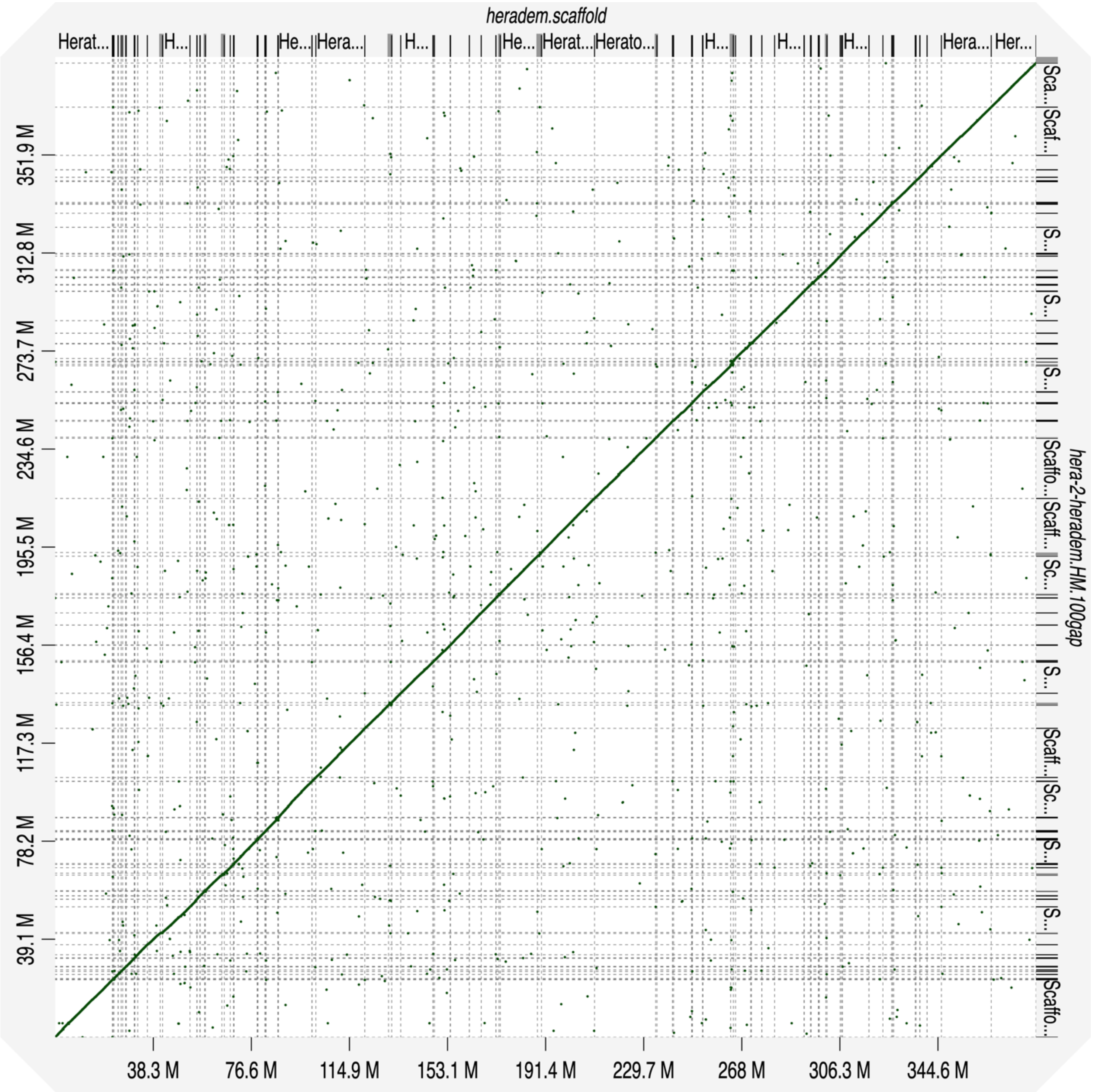


**Supplemental Fig. S30** - Dot plot showing the alignment of the reference-guided *H. erato* genome assembly (y-axis) to the *H. erato demophoon* reference genome (x-axis), used as reference to guide the scaffolding process. *H. erato* reference genome scaffolds are ordered, from chromosome 1 to chromosome 21.

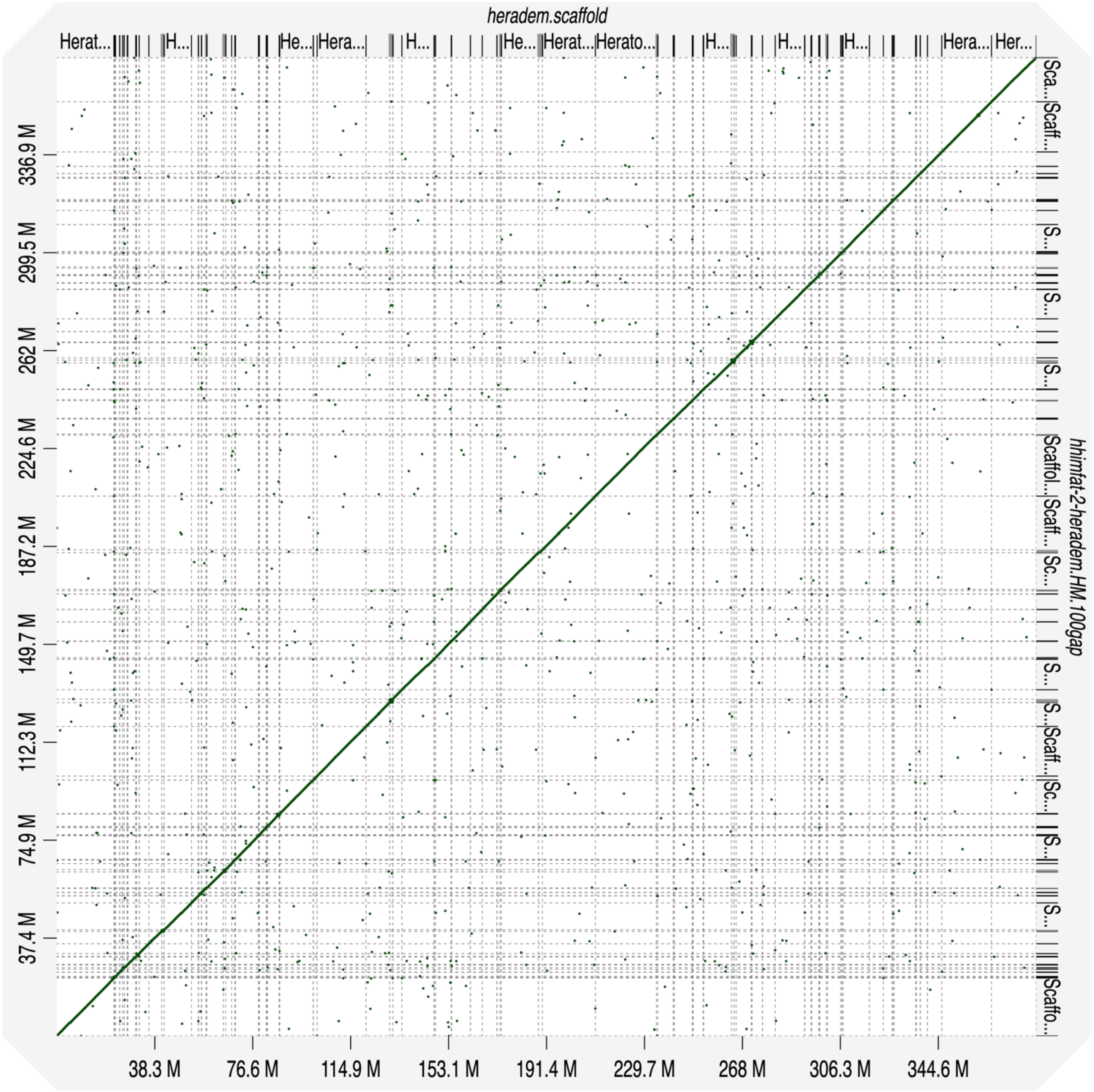


**Supplemental Fig. S31** - Dot plot showing the alignment of the reference-guided *H. himera* (father) genome assembly (y-axis) to the *H. erato demophoon* reference genome (x-axis), used as reference to guide the scaffolding process. *H. erato* reference genome scaffolds are ordered, from chromosome 1 to chromosome 21.

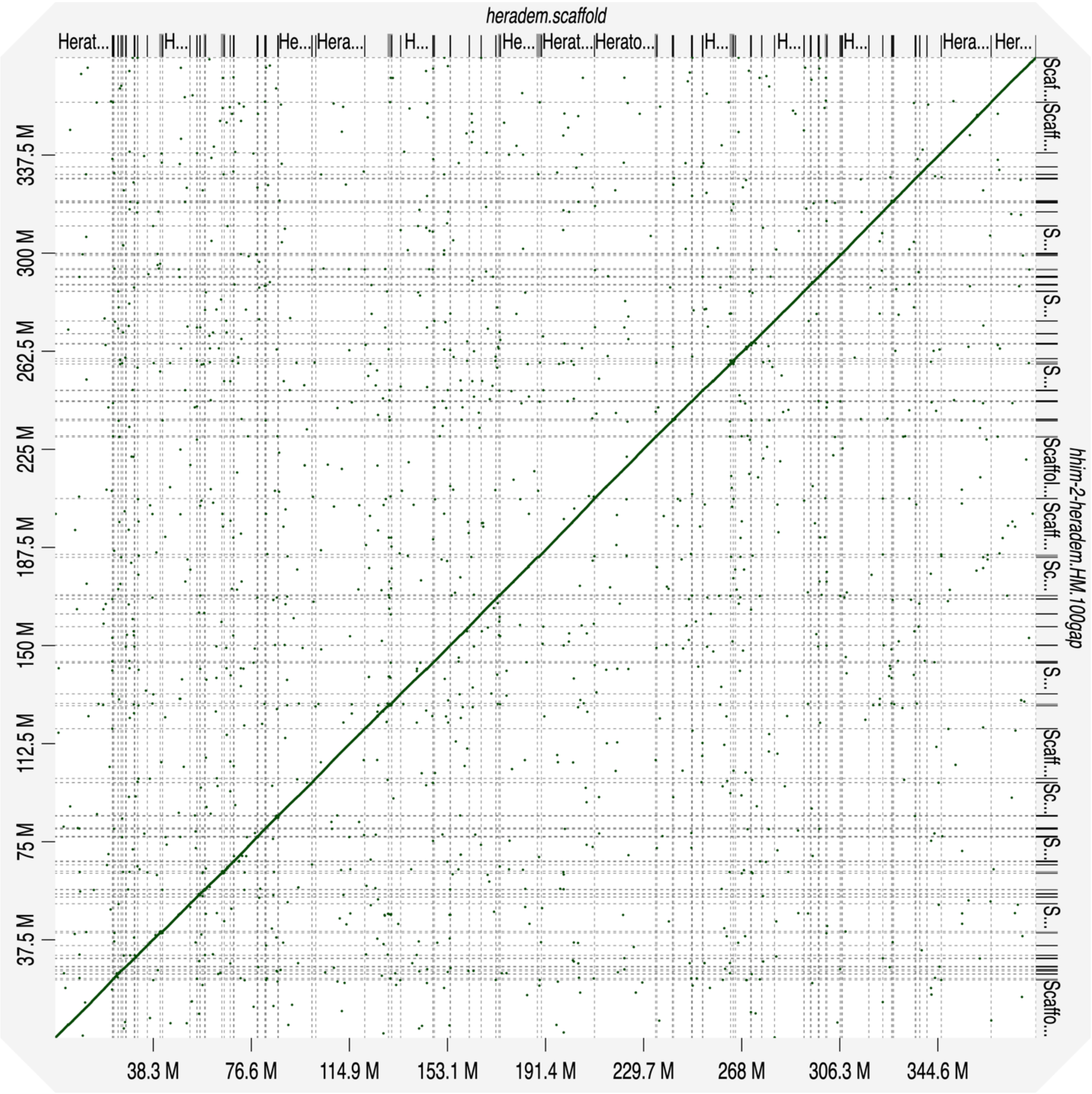


**Supplemental Fig. S32** - Dot plot showing the alignment of the reference-guided *H. himera* genome assembly (y-axis) to the *H. erato demophoon* reference genome (x-axis), used as reference to guide the scaffolding process. *H. erato* reference genome scaffolds are ordered, from chromosome 1 to chromosome 21.

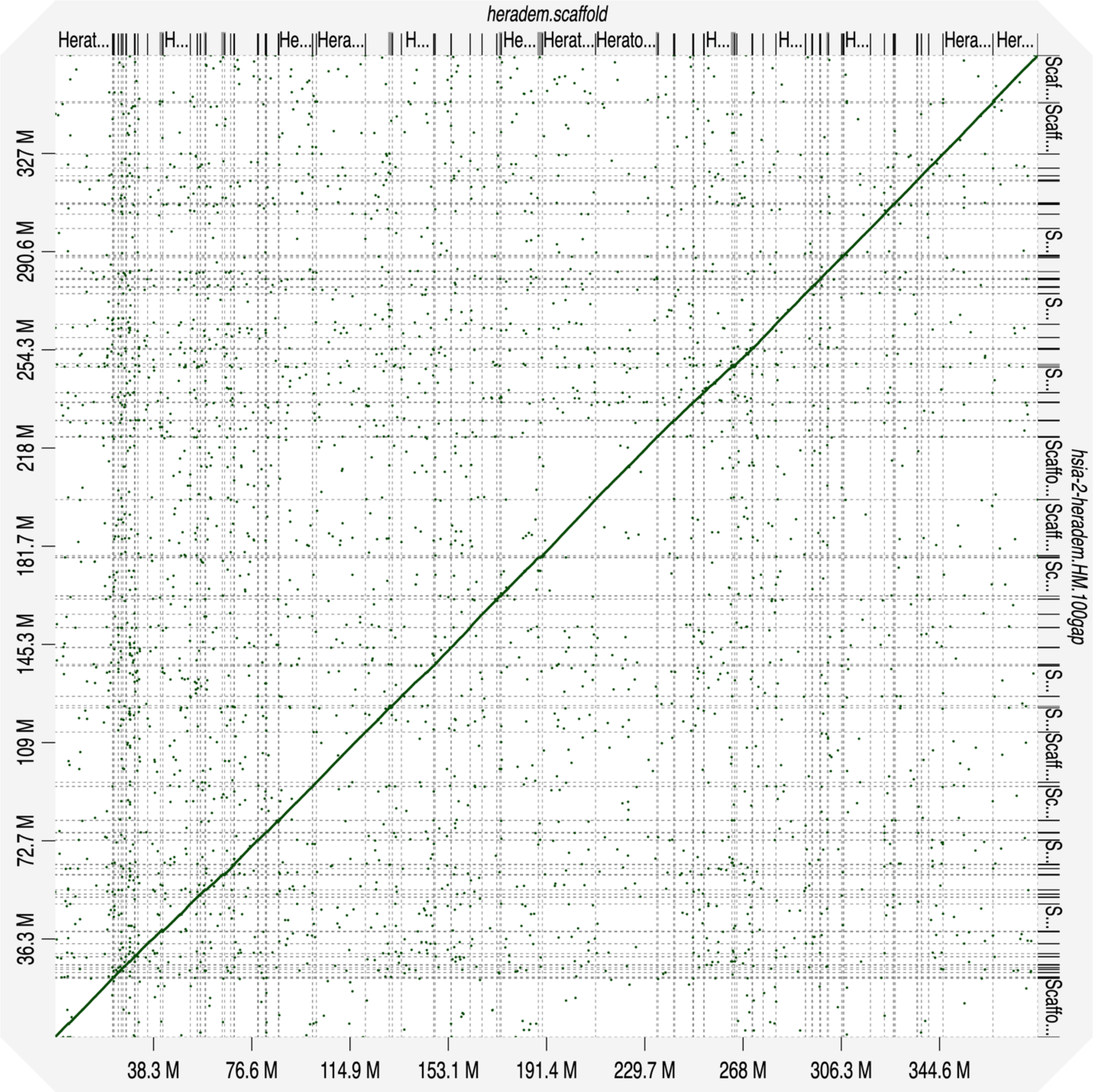


**Supplemental Fig. S33** - Dot plot showing the alignment of the reference-guided *H. hecalesia* genome assembly (y-axis) to the *H. erato demophoon* reference genome (x-axis), used as reference to guide the scaffolding process. *H. erato* reference genome scaffolds are ordered, from chromosome 1 to chromosome 21.

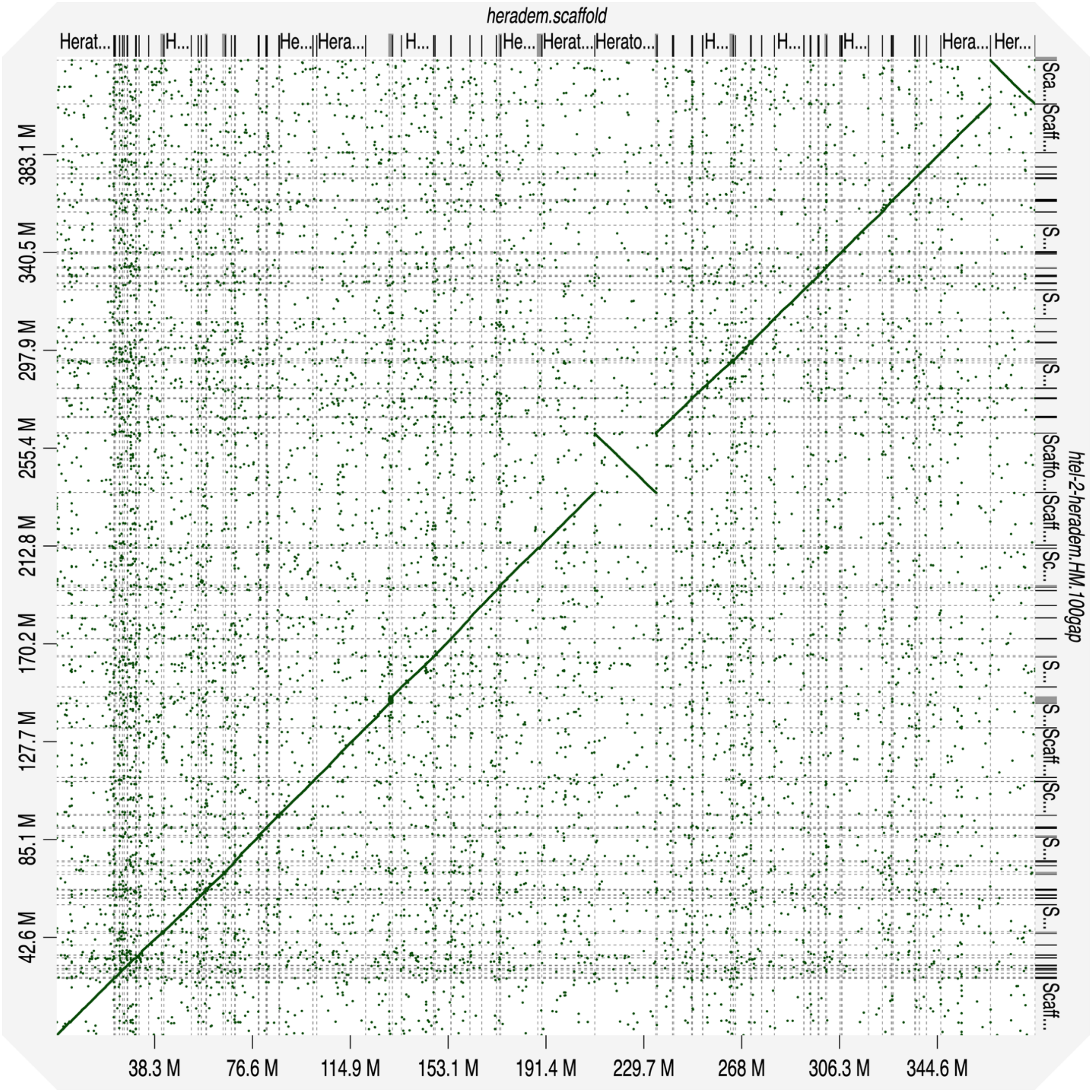


**Supplemental Fig. S34** - Dot plot showing the alignment of the reference-guided *H. telesiphe* genome assembly (y-axis) to the *H. erato demophoon* reference genome (x-axis), used as reference to guide the scaffolding process. *H. erato* reference genome scaffolds are ordered, from chromosome 1 to chromosome 21.

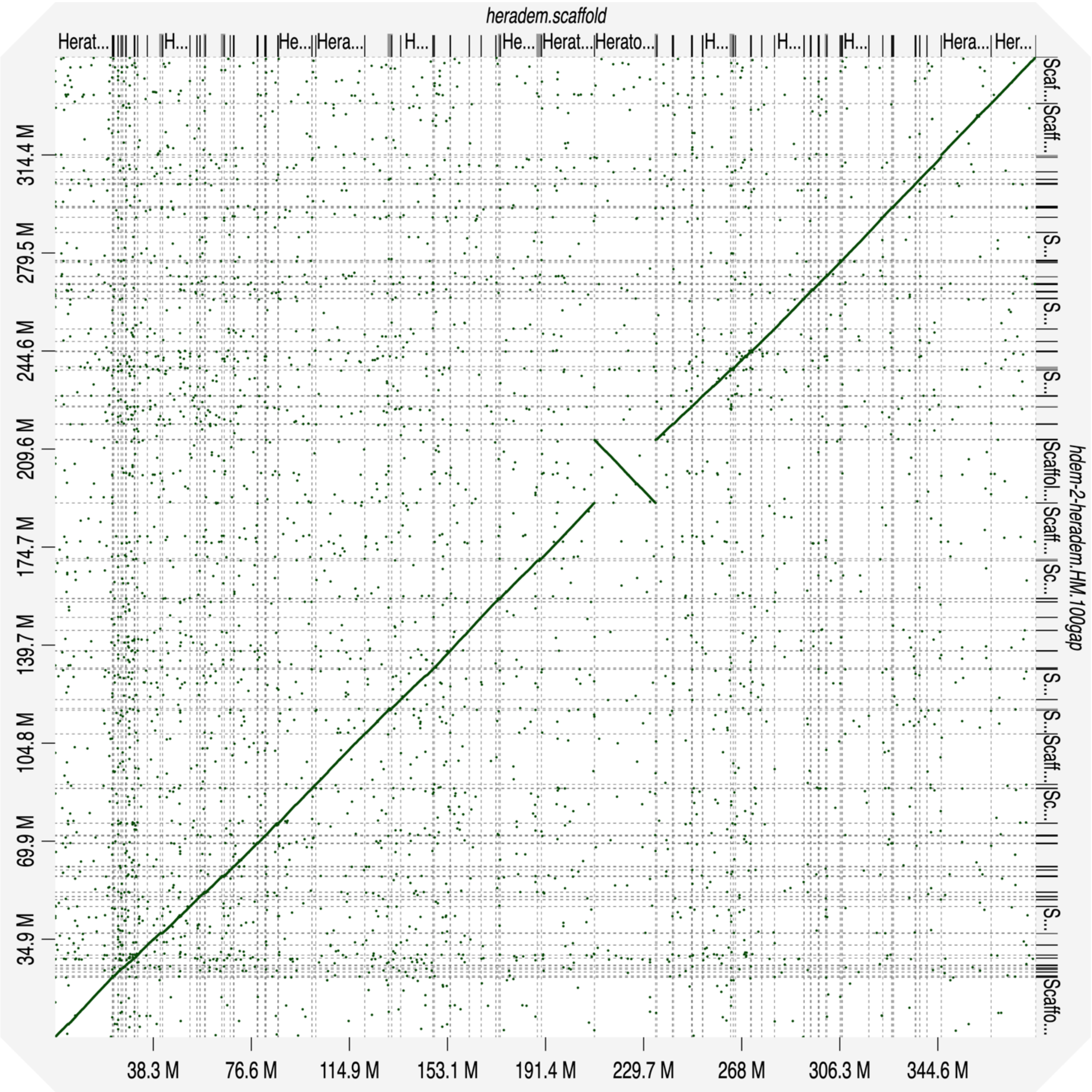


**Supplemental Fig. S35** - Dot plot showing the alignment of the reference-guided *H. demeter* genome assembly (y-axis) to the *H. erato demophoon* reference genome (x-axis), used as reference to guide the scaffolding process. *H. erato* reference genome scaffolds are ordered, from chromosome 1 to chromosome 21.

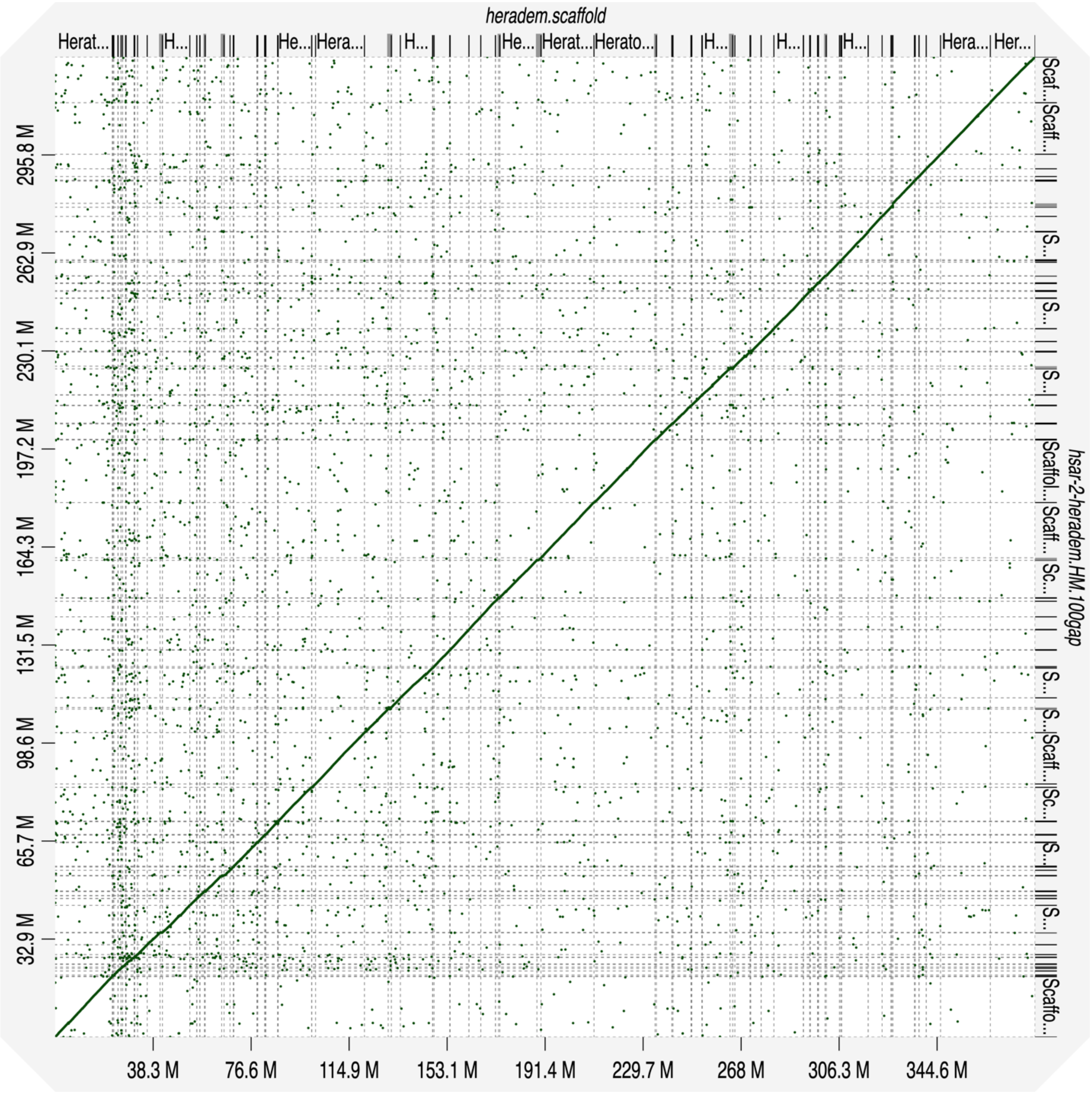


**Supplemental Fig. S36** - Dot plot showing the alignment of the reference-guided *H. sara* genome assembly (y-axis) to the *H. erato demophoon* reference genome (x-axis), used as reference to guide the scaffolding process. *H. erato* reference genome scaffolds are ordered, from chromosome 1 to chromosome 21.

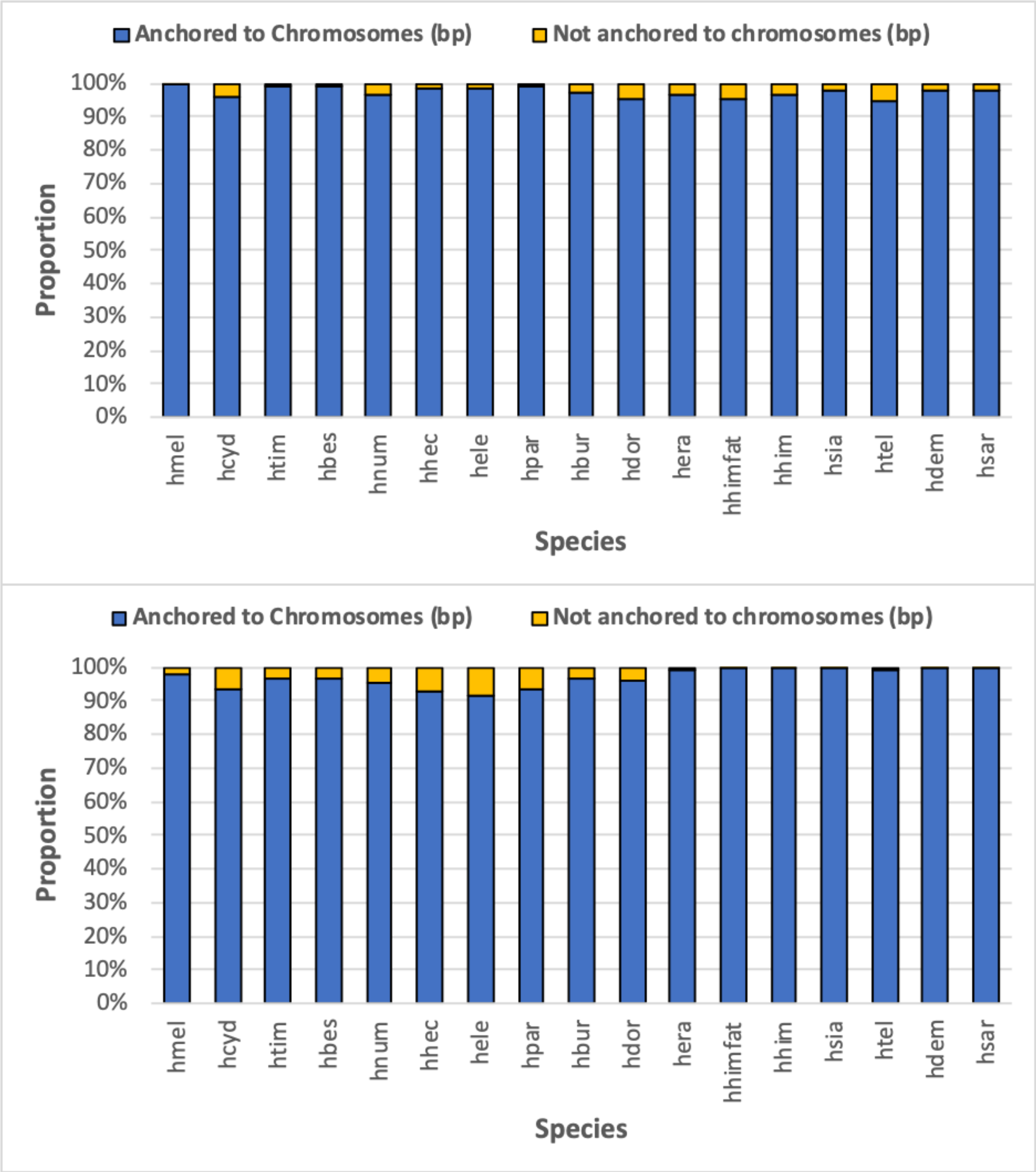


**Supplemental Fig. S37** - Proportion of the reference-scaffolded assemblies anchored to chromosomes, using *H. melpomene* (top panel) and *H. erato demophoon* (bottom panel) as reference. Species codes are as in Figure 1.

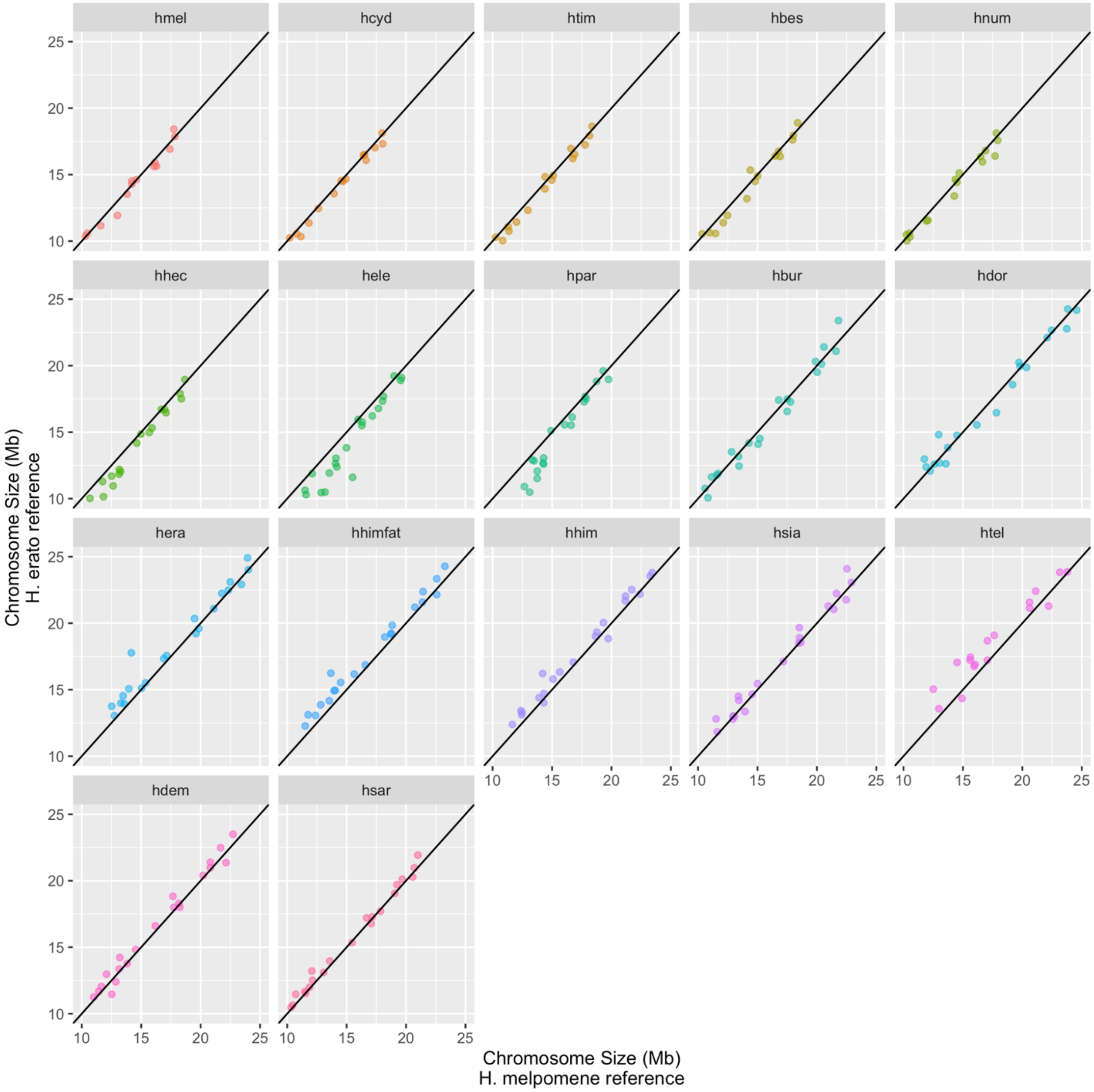


**Supplemental Fig. S38** - Chromosome size correlations between the reference-guided assemblies using either *H. melpomene* (x-axis) or *H. erato demophoon* (y-axis) to guide the scaffolding process. Species codes are as in Figure 1.

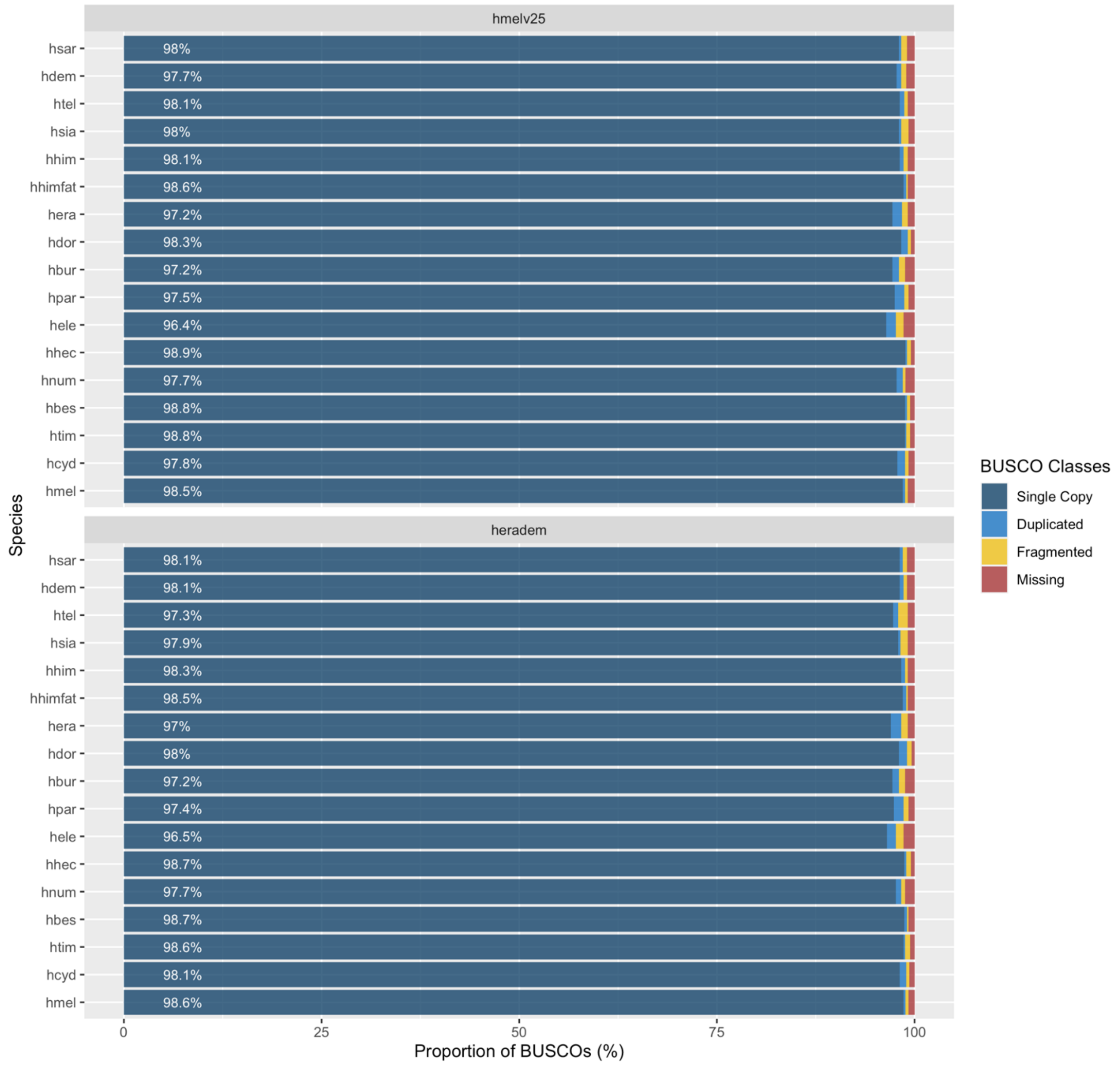


**Supplemental Fig. S39** - BUSCO gene content, for the reference-guided assemblies based on the *H. melpomene* (hmelv25; top panel) and the *H. erato demophoon* (heradem; bottom panel) reference genomes. Species codes are as in Figure 1.

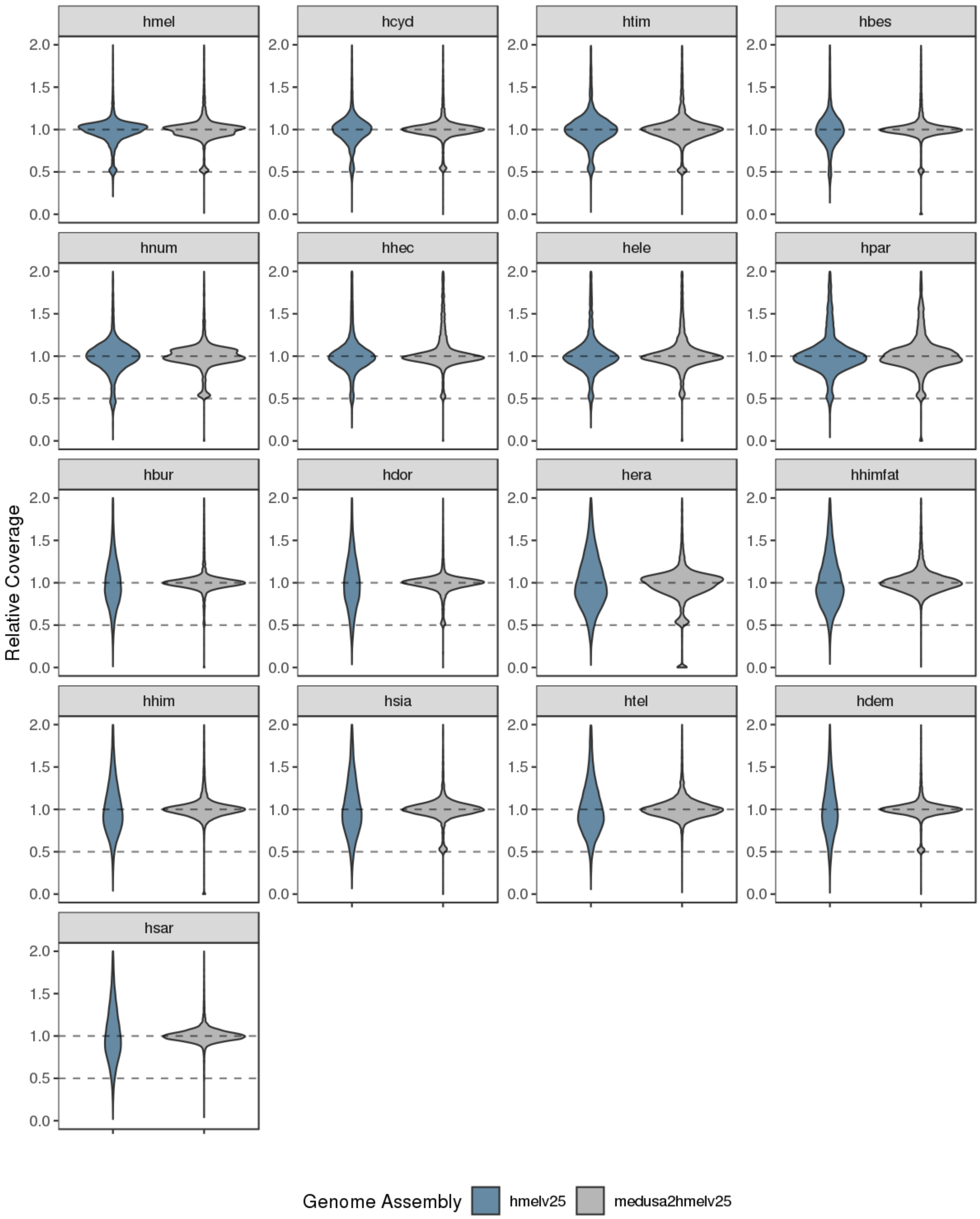


**Supplemental Fig. S40** – Relative depth of coverage across the genome when mapping reads to the *H. melpomene* reference genome (hmelv25) or against the reference-guided assembly of the same species, based on the same reference genome (medusa2hmelv25; e.g. *H cydno* to the *H. cydno* reference-guided assembly, guided by the *H. melpomene* reference genome). Species codes are as in Figure 1.

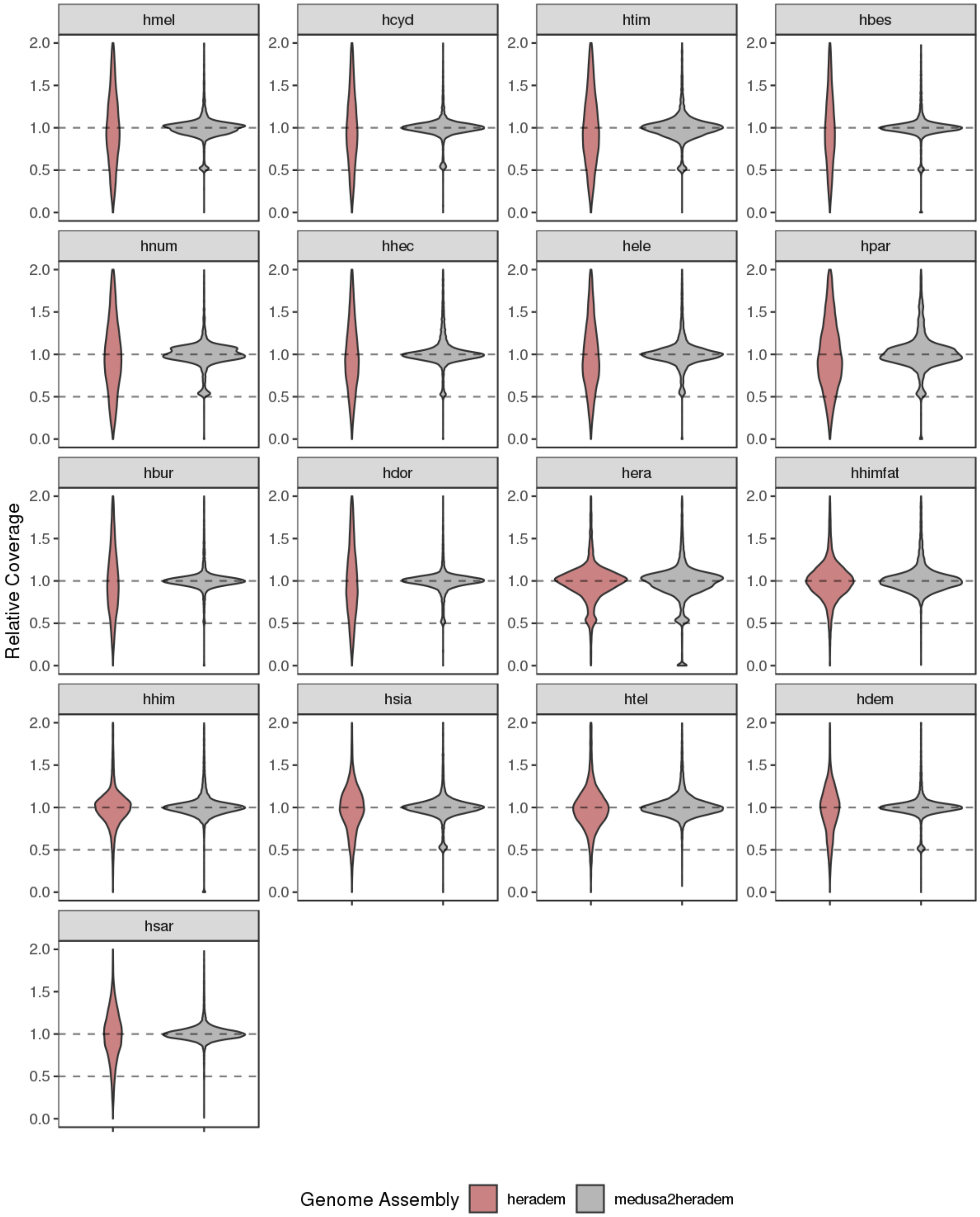


**Supplemental Fig. S41** - Relative depth of coverage across the genome when mapping reads to the *H. erato demophoon* reference genome (heradem) or against the reference-guided assembly of the same species, based on the same reference genome (medusa2heradem; e.g. *H cydno* to the *H. cydno* reference-guided assembly, guided by the *H. erato demophoon* reference genome). Species codes are as in Figure 1.

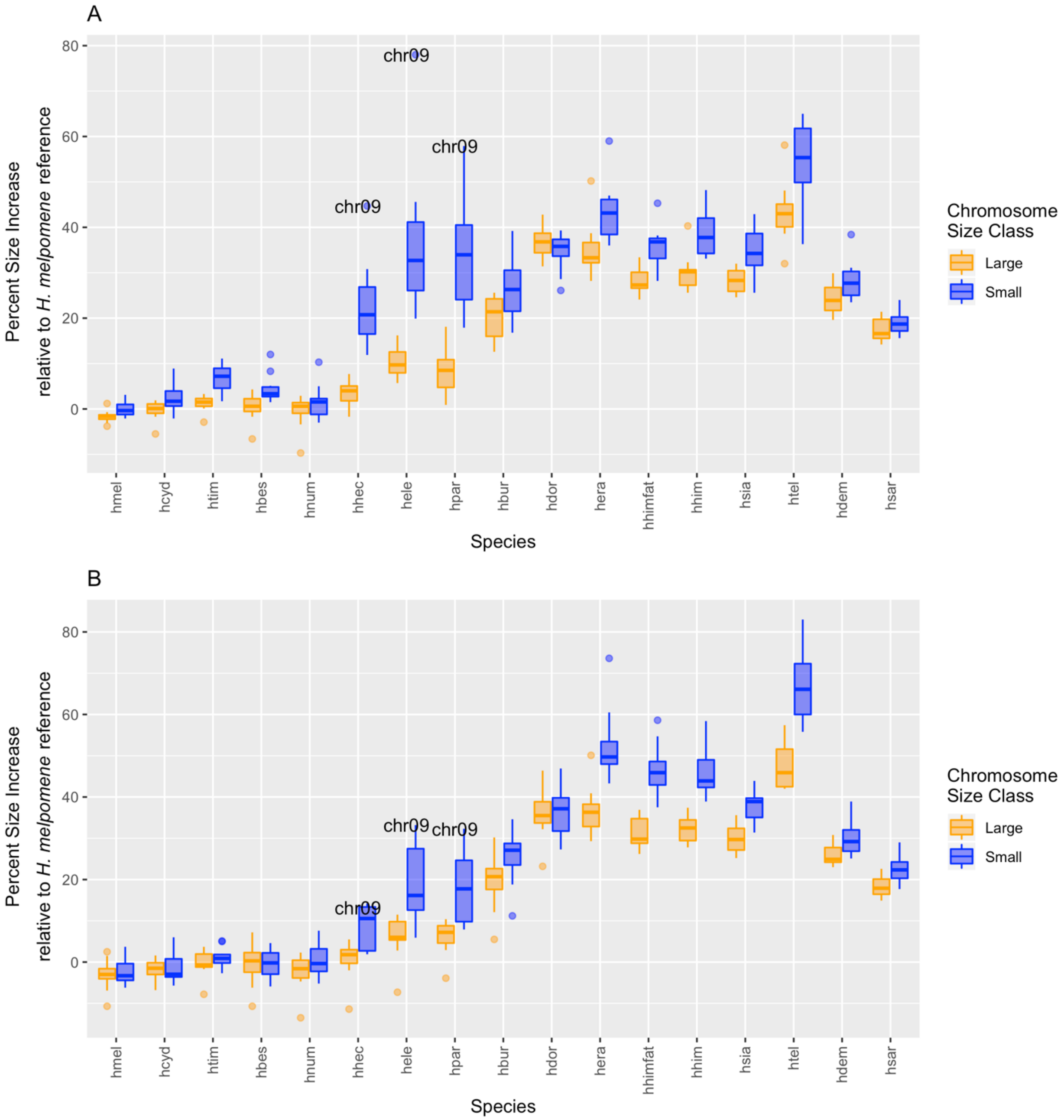


**Supplemental Fig. S42** – Percent increase in size of reference-guided assemblies chromosomes relative to *H. melpomene* reference genome chromosome sizes. Comparisons were performed based on the reference-guided assemblies using both *H. melpomene* (A) and *H. erato* demophoon references (B). Increase In size is generally more accentuated in small chromosomes (blue) and within the *melpomene*-silvaniform clade is more marked for the *H. hecale*, *H. elevatus* and *H. pardalinus* species trio. Species codes are as in Figure 1.

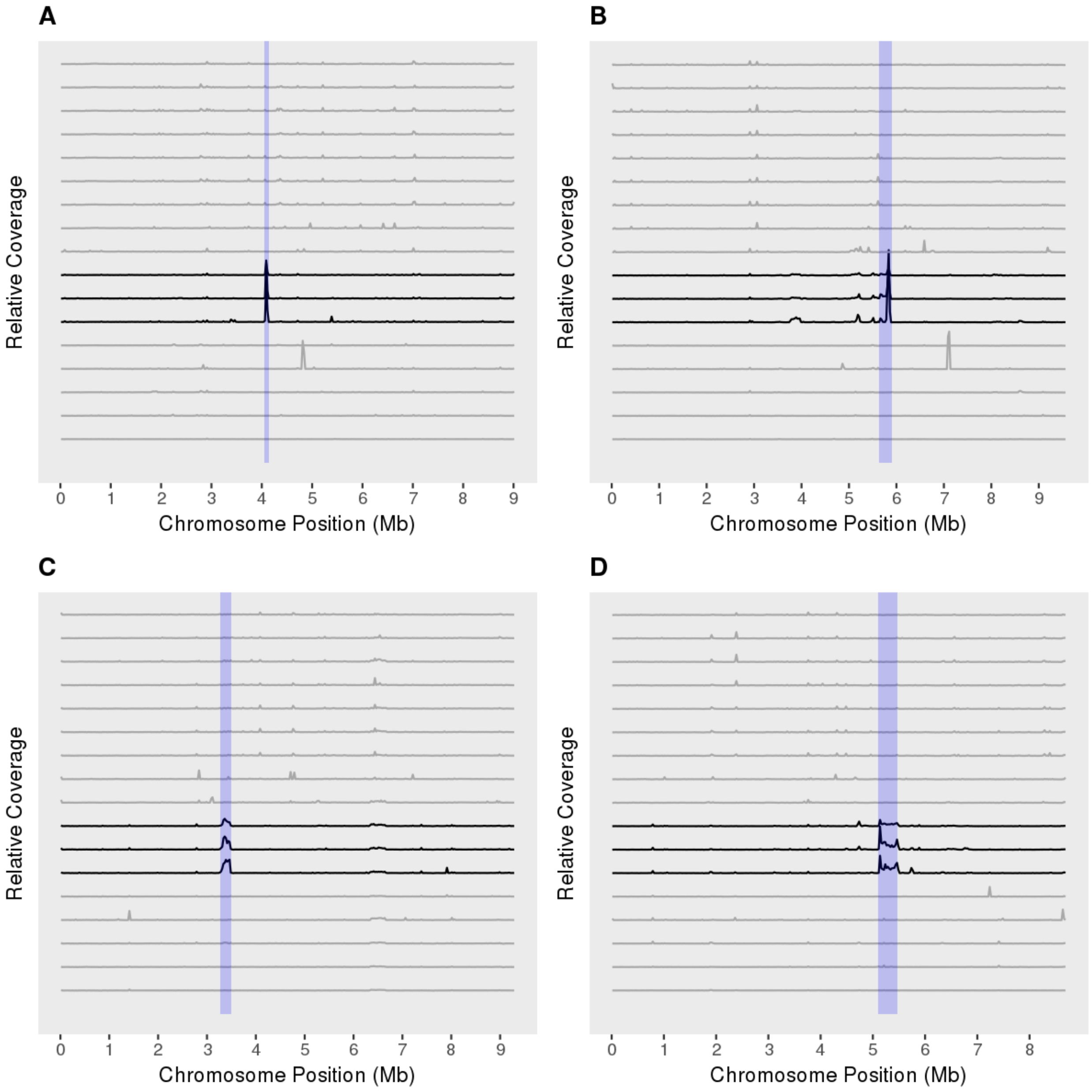


**Supplemental Fig. S43** – Genomic regions with exceptionally high coverage specifically in the *H. hecale*, *H. elevatus* and *H. pardalinus* species trio. The relative coverage (y-axis) along the chromosomes (x-axis) is based on the mapping of short-sequencing data of each of the species to the *H. melpomene* reference, and calculated in 25 kb non-overlapping windows. The species trio relative coverages are depicted by the black lines, while grey lines represent the relative coverage in all other species. The blue bars depict the genomic regions of interest. In order, panels A-D represent chromosomes 2, 4, 8 and 9. In each panel, species are in the following order (from bottom to top): hmel – *H. melpomene*; hcyd – *H. cydno*; htim – *H. timareta*; hbes – *H. besckei*; hnum – *H. numata*; hhec- *H. hecale*; hele – *H. elevatus*; hpar – *H. pardalinus*; hbur – *H. burneyi*; hdor – *H. doris*; hera – *H. erato*; hhimfat – *H. himera* (father); hhim – *H. himera*; hsia – *H. hecalesia*; htel – *H. telesiphe*; hdem – *H. demeter*; hsar – *H. sara*.

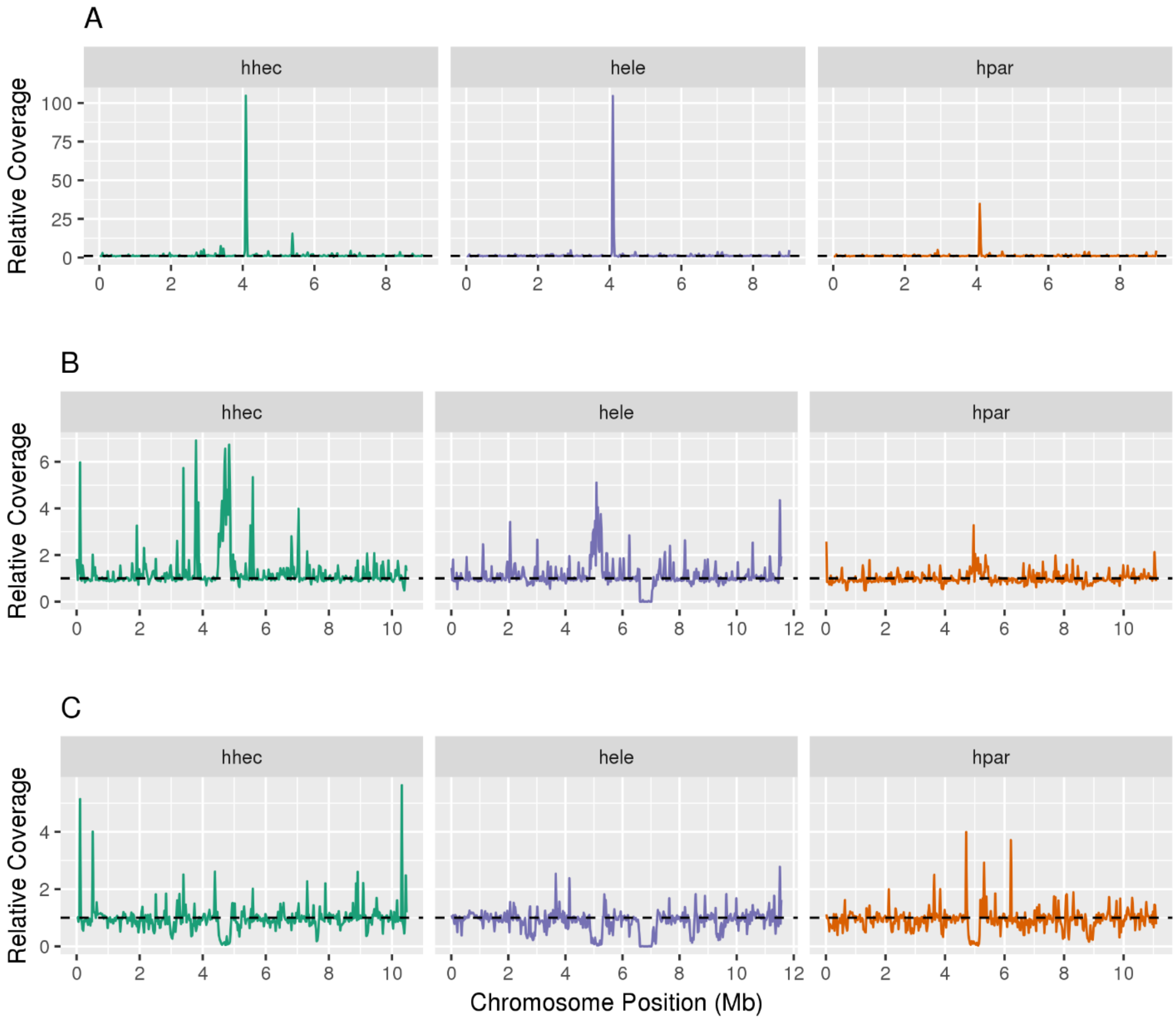


**Supplemental Fig. S44** - Relative depth of coverage of chromosome 2. (A) *H. hecale*, *H. elevatus* and *H. pardalinus* original *w2rap* reads mapped onto the *H. melpomene* reference genome. (B) *H. hecale*, *H. elevatus* and *H. pardalinus* reads mapped onto their own reference-guided scaffold assemblies. (C) H. melpomene reads mapped onto the reference-guided assemblies of the species trio. In panel A, the repeat regions are identifiable by the sharp increase in relative coverage which is less accentuated when mapping to the reference-guided assemblies (note scale differences for both axes, panel B). Mapping *H. melpomene* reads to the reference-guided assemblies of the species trio allows detection of the repeat regions in these three species, which is shown by a drop in coverage (panel C). The repeat regions are ca. 375 kb (*H. hecale*), 425 kb (*H. elevatus*) and 400 Mb (*H. pardalinus*) long while only 50 kb long in *H. melpomene*. Species codes: hhec – *H. hecale*; hele – *H. elevatus*; hpar – *H. pardalinus*.

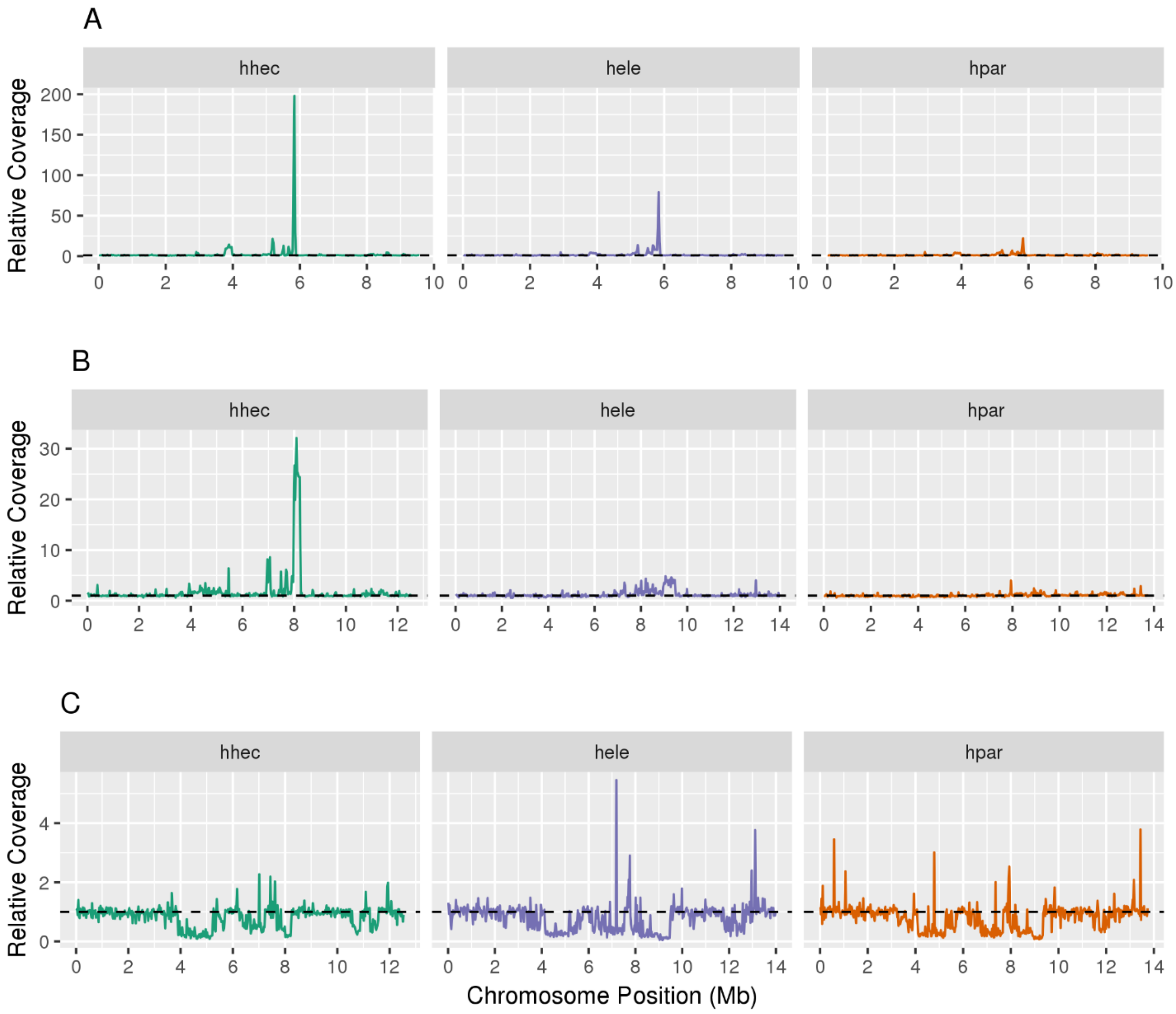


**Supplemental Fig. S45** - Relative depth of coverage of chromosome 4. (A) *H. hecale*, *H. elevatus* and *H. pardalinus* original *w2rap* reads mapped onto the *H. melpomene* reference genome. (B) *H. hecale*, *H. elevatus* and *H. pardalinus* reads mapped onto their own reference-guided scaffold assemblies. (C) *H. melpomene* reads mapped onto the reference-guided assemblies of the species trio. In panel A, the repeat regions are identifiable by the sharp increase in relative coverage which is less accentuated when mapping to the reference-guided assemblies (note scale differences for both axis; panel B). Mapping of *H. melpomene* reads to the three reference-guided assemblies of the species trio allows detection of the repeat regions in these three species, which is shown by a drop in coverage (panel C). The repeat regions are ca. 1.450 Mb (*H. hecale*), 1.350 Mb (*H. elevatus*) and 1.600 Mb (*H. pardalinus*) long while only 225 kb long in *H. melpomene*. Species codes: hhec – *H. hecale*; hele – *H. elevatus*; hpar – *H. pardalinus*.

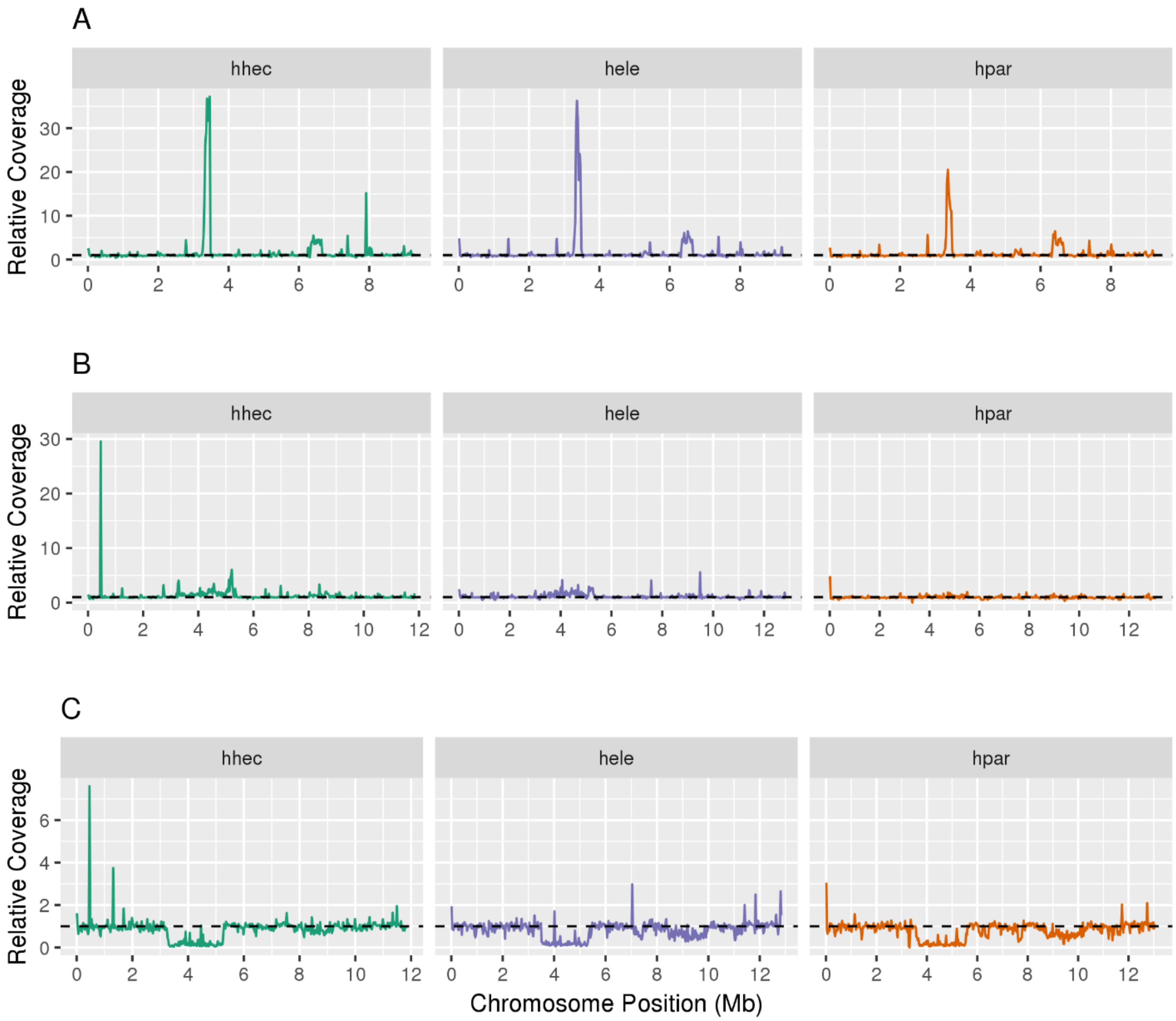


**Supplemental Fig. S46** – Relative depth of coverage of chromosome 8. (A) *H. hecale*, *H. elevatus* and *H. pardalinus* original *w2rap* reads mapped onto the *H. melpomene* reference genome. (B) *H. hecale*, *H. elevatus* and *H. pardalinus* reads mapped onto their own reference-guided scaffold assemblies. (C) *H. melpomene* reads mapped onto the reference-guided assemblies of the species trio. In panel A, the repeat regions are identifiable by the sharp increase in relative coverage which is less accentuated when mapping to the reference-guided assemblies (note scale differences for both axes, panel B). Mapping of *H. melpomene* reads to the three reference-guided assemblies of the species trio allows detection of the repeat regions in these three species, which is shown by a drop in coverage (panel C). The repeat regions are ca. 1.95 Mb (*H. hecale*), 1.750 Mb (*H. elevatus*) and 2.250 Mb (*H. pardalinus*) long while only 175 kb long in *H. melpomene*. Species codes: hhec – *H. hecale*; hele – *H. elevatus*; hpar – *H. pardalinus*.

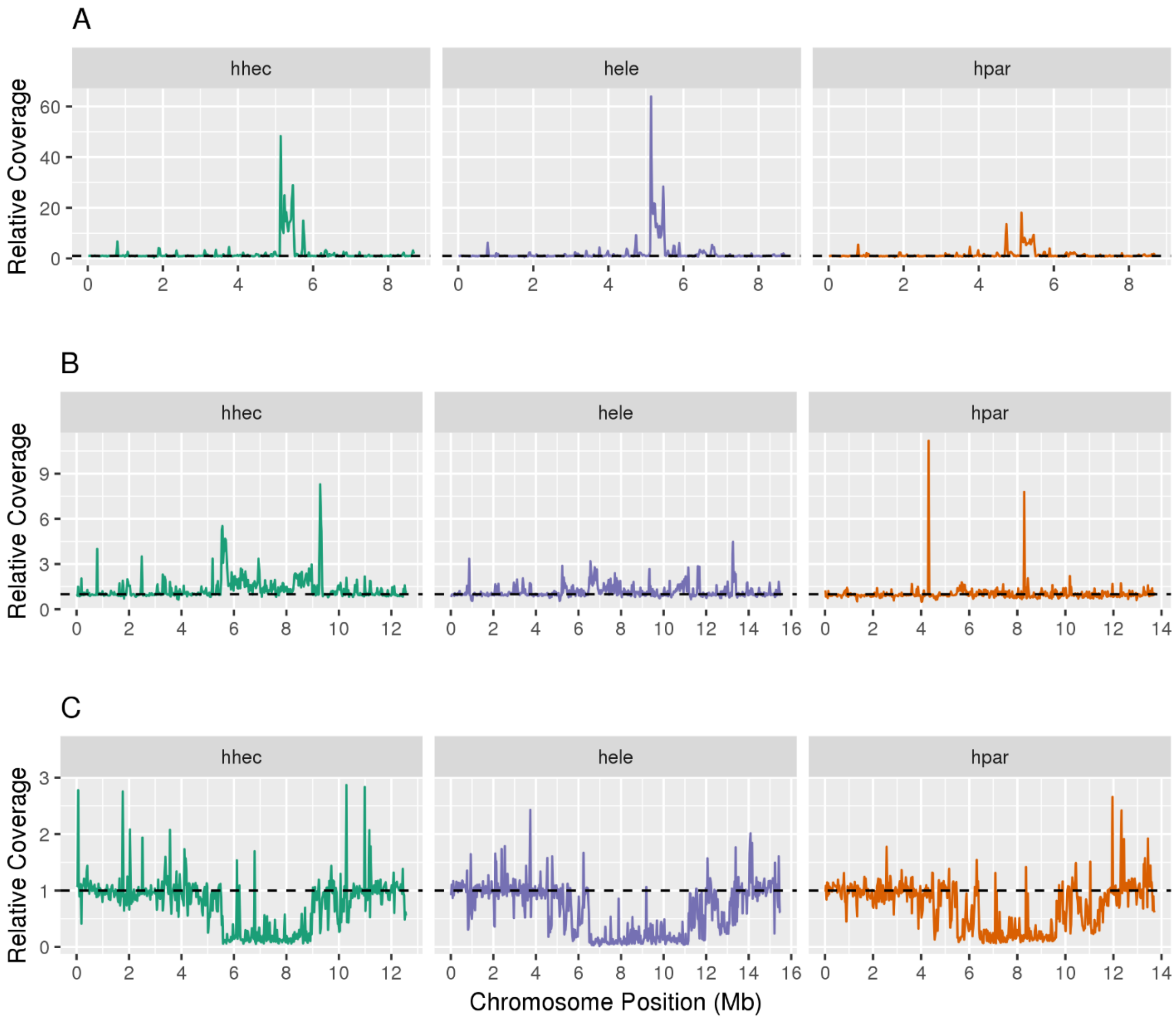


**Supplemental Fig. S47** - Relative depth of coverage of chromosome 9. (A) *H. hecale*, *H. elevatus* and *H. pardalinus* original *w2rap* reads mapped onto the *H. melpomene* reference genome. (B) *H. hecale*, *H. elevatus* and *H. pardalinus* reads mapped onto their own reference-guided scaffold assemblies. (C) *H. melpomene* reads mapped onto the reference-guided assemblies of the species trio. In panel A, the repeat regions are identifiable by the sharp increase in relative coverage which is less accentuated when mapping to the reference-guided assemblies (note scale differences for both axes, panel B). Mapping of *H. melpomene* reads to the three reference-guided assemblies of the species trio allows detection of the repeat regions in these three species, which is shown by a drop in coverage (panel C). The repeat regions are ca. 3.350 Mb (*H. hecale*), 3.925 Mb (*H. elevatus*) and 4.125 Mb (*H. pardalinus*) long while only 325 kb long in *H. melpomene*. Species codes: hhec – *H. hecale*; hele – *H. elevatus*; hpar – *H. pardalinus*.

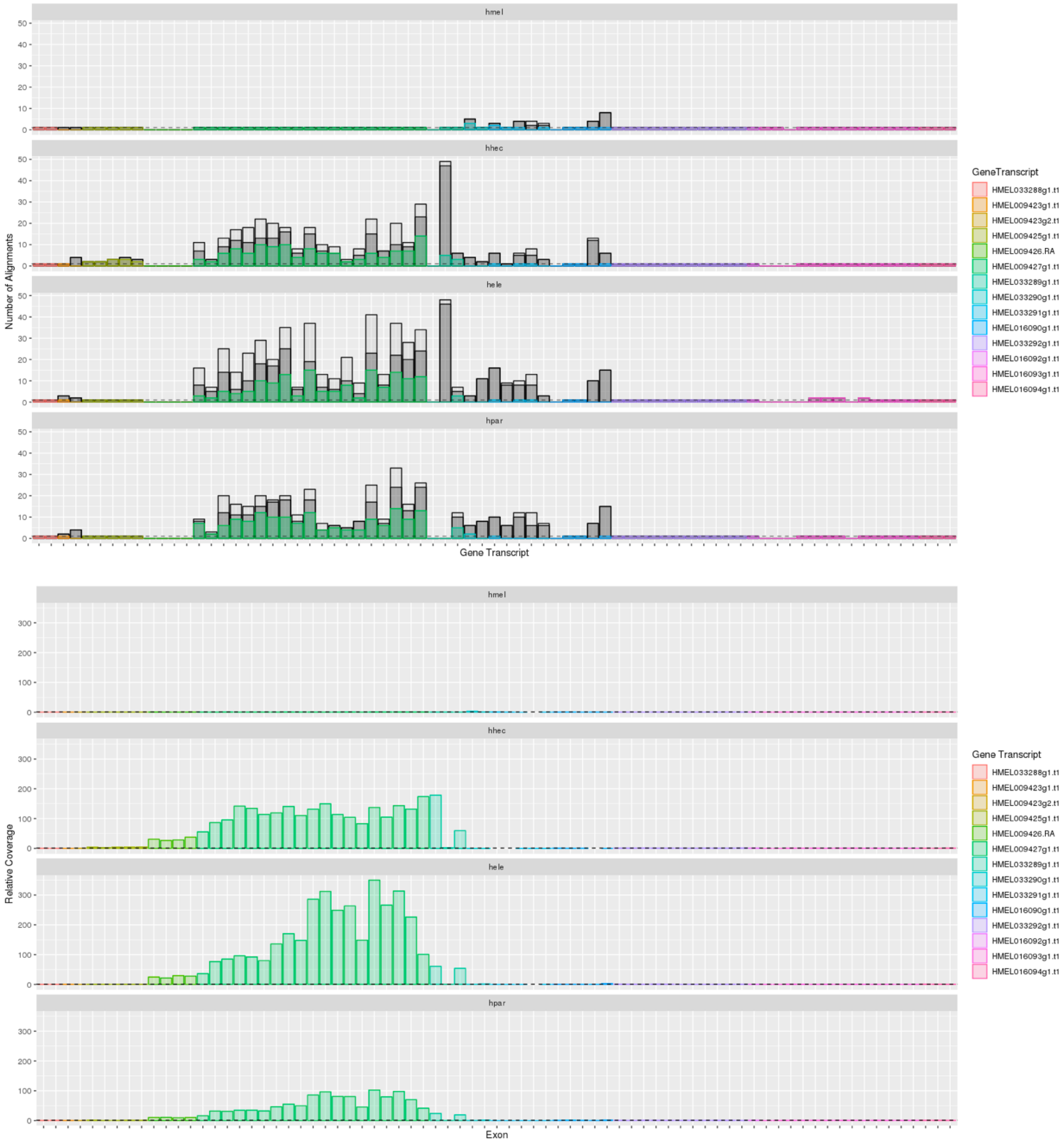


**Supplemental Fig. S48** – Exon copy number (CN) estimates in the repeat region on chromosome 2. CN estimates are based either on the number of alignments of *H. melpomene* reference exon sequences to the new assemblies (top panel) or normalized read coverage based on mapping to the *H. melpomene* assembly (bottom panel). On the top panel, colored bars depict the number of alignments to the expected chromosome, while the dark grey bars depict alignments to other chromosomes and the light grey bars alignments to scaffolds not anchored to chromosomes. The dashed horizontal lines on both panels represent a CN of one. The new *H. melpomene* assembly was also included as a control. Species codes: hmel -*H. melpomene*; hhec - *H. hecale*; hele - *H. elevatus*; hpar - *H. pardalinus.*

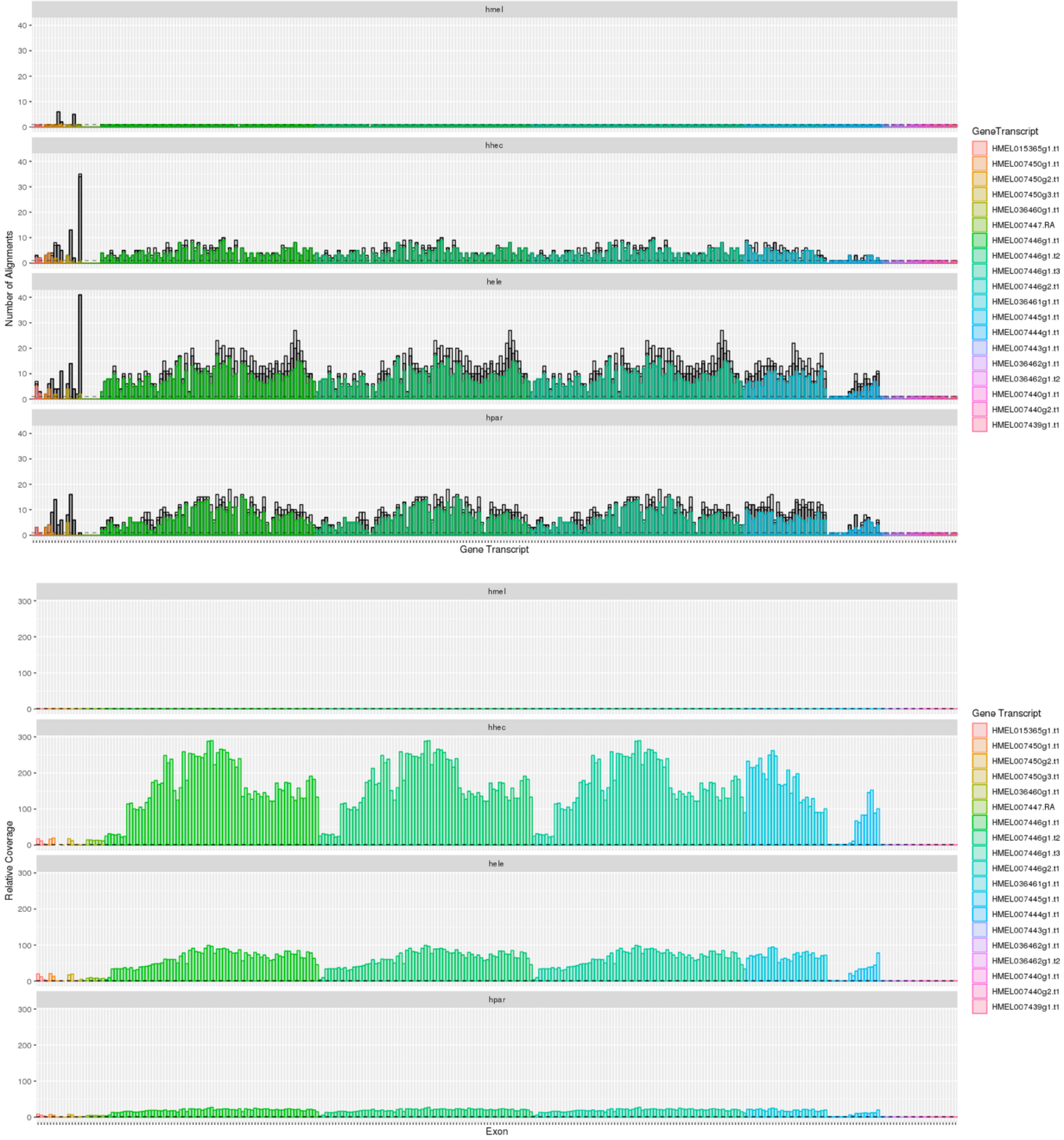


**Supplemental Fig. S49** - Exon copy number (CN) estimates in the repeat region on chromosome 4. CN estimates are based either on the number of alignments of *H. melpomene* reference exon sequences to the new assemblies (top panel) or normalized read coverage based on mapping to the *H. melpomene* assembly (bottom panel). On the top panel, colored bars depict the number of alignments to the expected chromosome, while the dark grey bars depict alignments to other chromosomes and the light grey bars alignments to scaffolds not anchored to chromosomes. The dashed horizontal lines on both panels represent a CN of one. The new *H. melpomene* assembly was also included as a control. Species codes: hmel - *H. melpomene*; hhec - *H. hecale*; hele - *H. elevatus*; hpar - *H. pardalinus*.

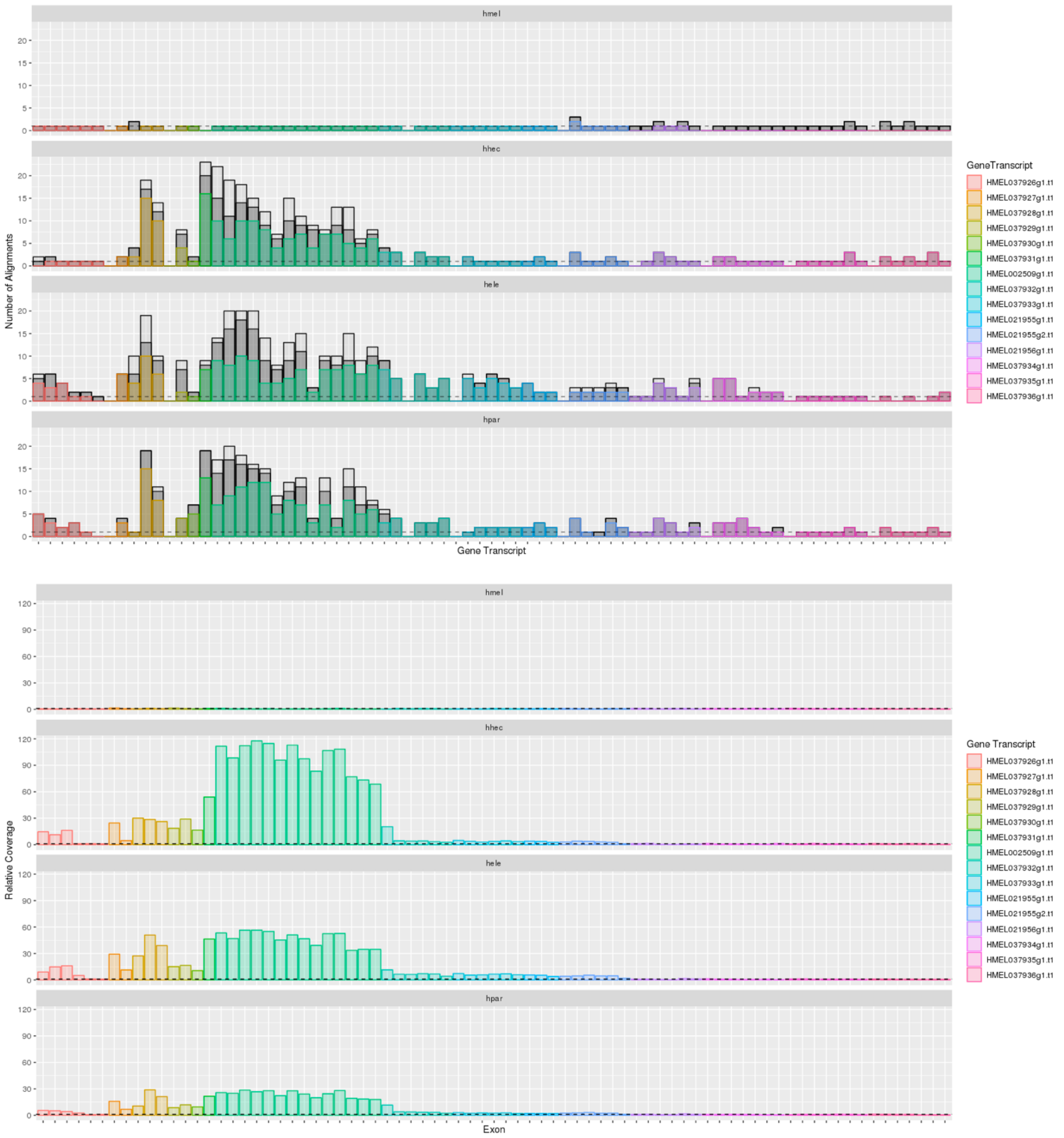


**Supplemental Fig. S50** - Exon copy number (CN) estimates in the repeat region on chromosome 8. CN estimates are based either on the number of alignments of *H. melpomene* reference exon sequences to the new assemblies (top panel) or normalized read coverage based on mapping to the *H. melpomene* assembly (bottom panel). On the top panel, colored bars depict the number of alignments to the expected chromosome, while the dark grey bars depict alignments to other chromosomes and the light grey bars alignments to scaffolds not anchored to chromosomes. The dashed horizontal lines on both panels represent a CN of one. The new *H. melpomene* assembly was also included as a control. Species codes: hmel - *H. melpomene*; hhec - *H. hecale*; hele - *H. elevatus*; hpar - *H. pardalinus*.

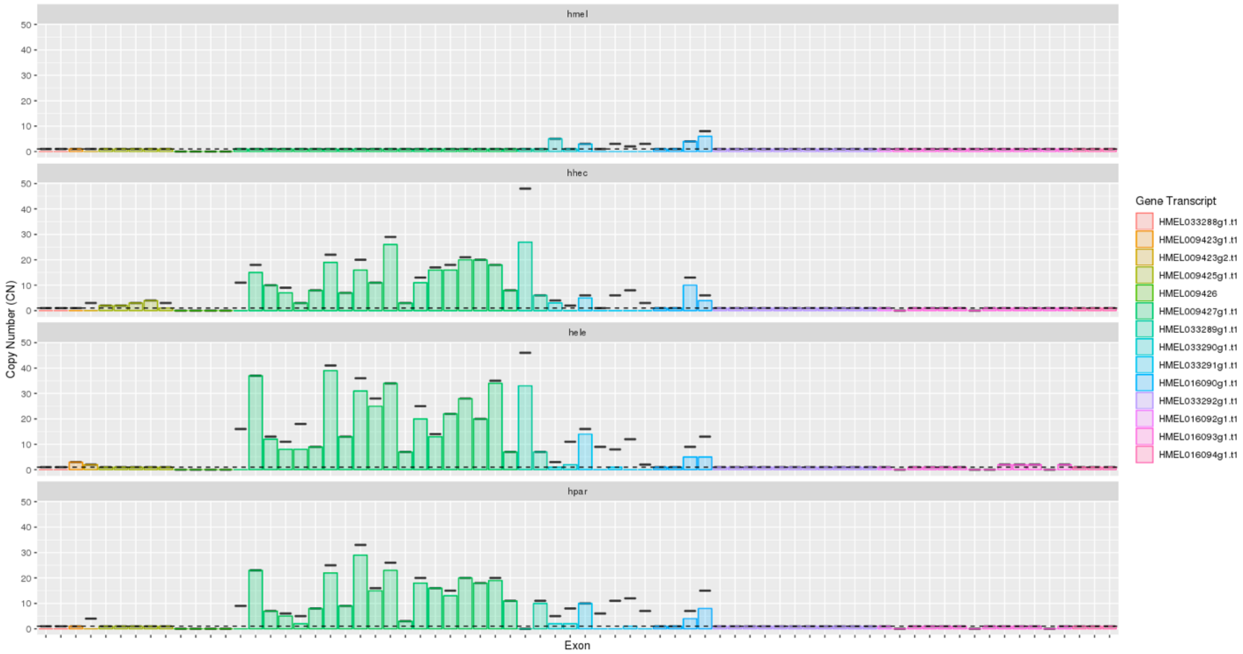


**Supplemental Fig. S51** – Exonic CNs on chromosome 2 before (black lines) and after (colored bars) filtering for stop codons. The dashed black line indicates CN of 1. Species codes: hmel - *H. melpomene*; hhec - *H. hecale*; hele - *H. elevatus*; hpar - *H. pardalinus*.

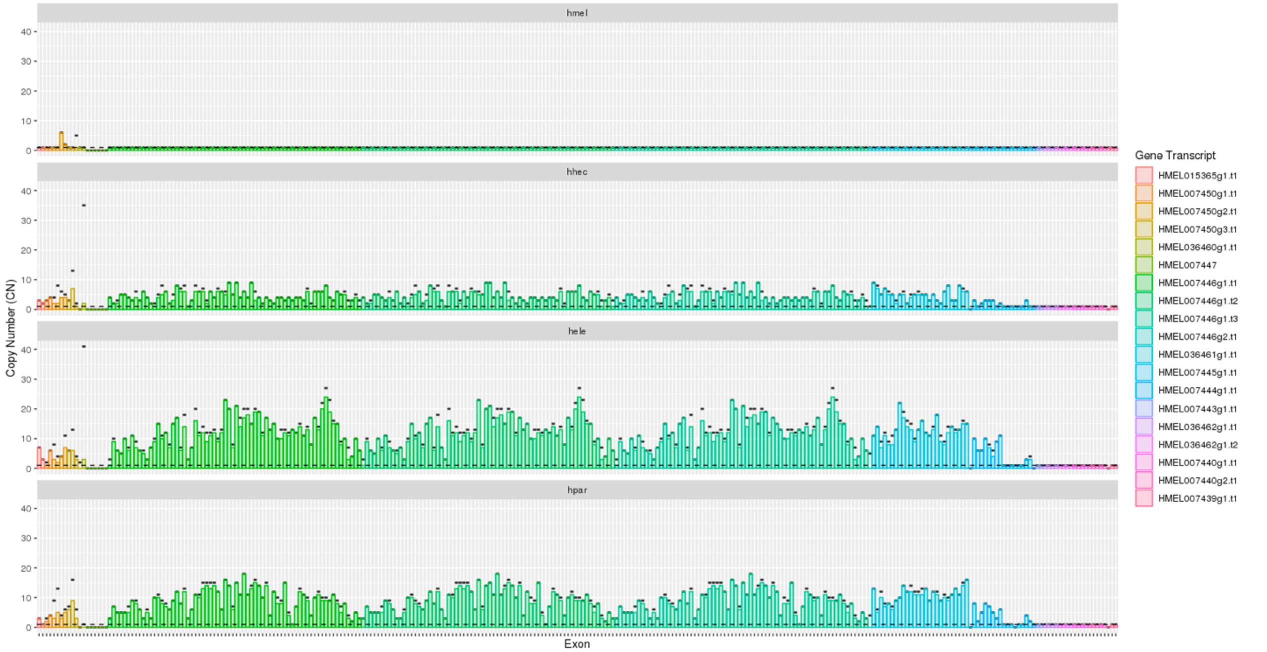


**Supplemental Fig. S52** - Exonic CNs on chromosome 4 before (black lines) and after (colored bars) filtering for stop codons. The dashed black line indicates CN of 1. Species codes: hmel -*H. melpomene*; hhec - *H. hecale*; hele - *H. elevatus*; hpar - *H. pardalinus*.

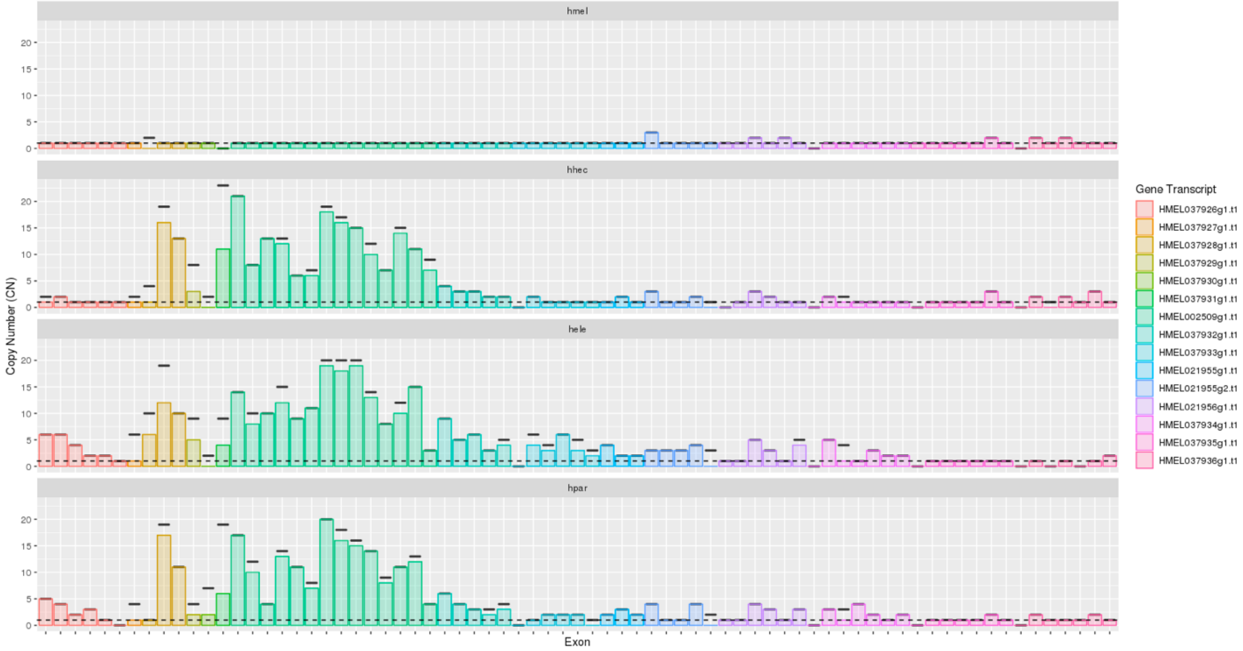


**Supplemental Fig. S53** - Exonic CNs on chromosome 8 before (black lines) and after (colored bars) filtering for stop codons. The dashed black line indicates CN of 1. Species codes: hmel - *H. melpomene*; hhec - *H. hecale*; hele - *H. elevatus*; hpar - *H. pardalinus*.

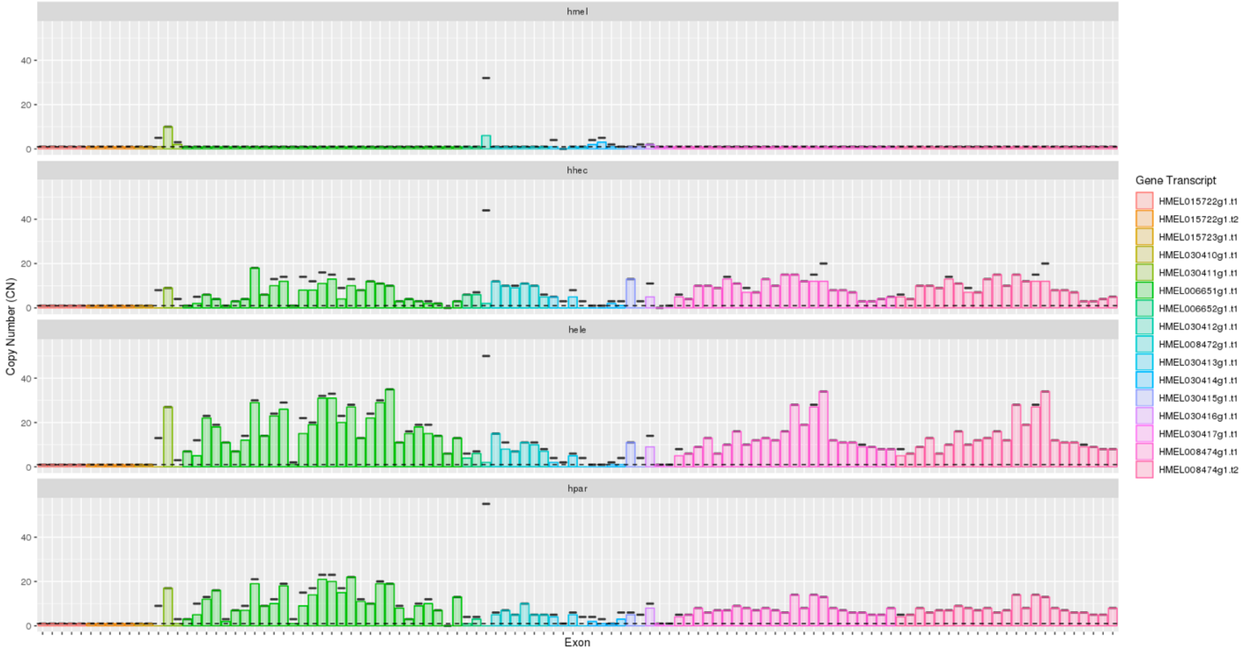


**Supplemental Fig. S54** - Exonic CNs on chromosome 9 before (black lines) and after (colored bars) filtering for stop codons. The dashed black line indicates CN of 1. Species codes: hmel - *H. melpomene*; hhec - *H. hecale*; hele - *H. elevatus*; hpar - *H. pardalinus*.

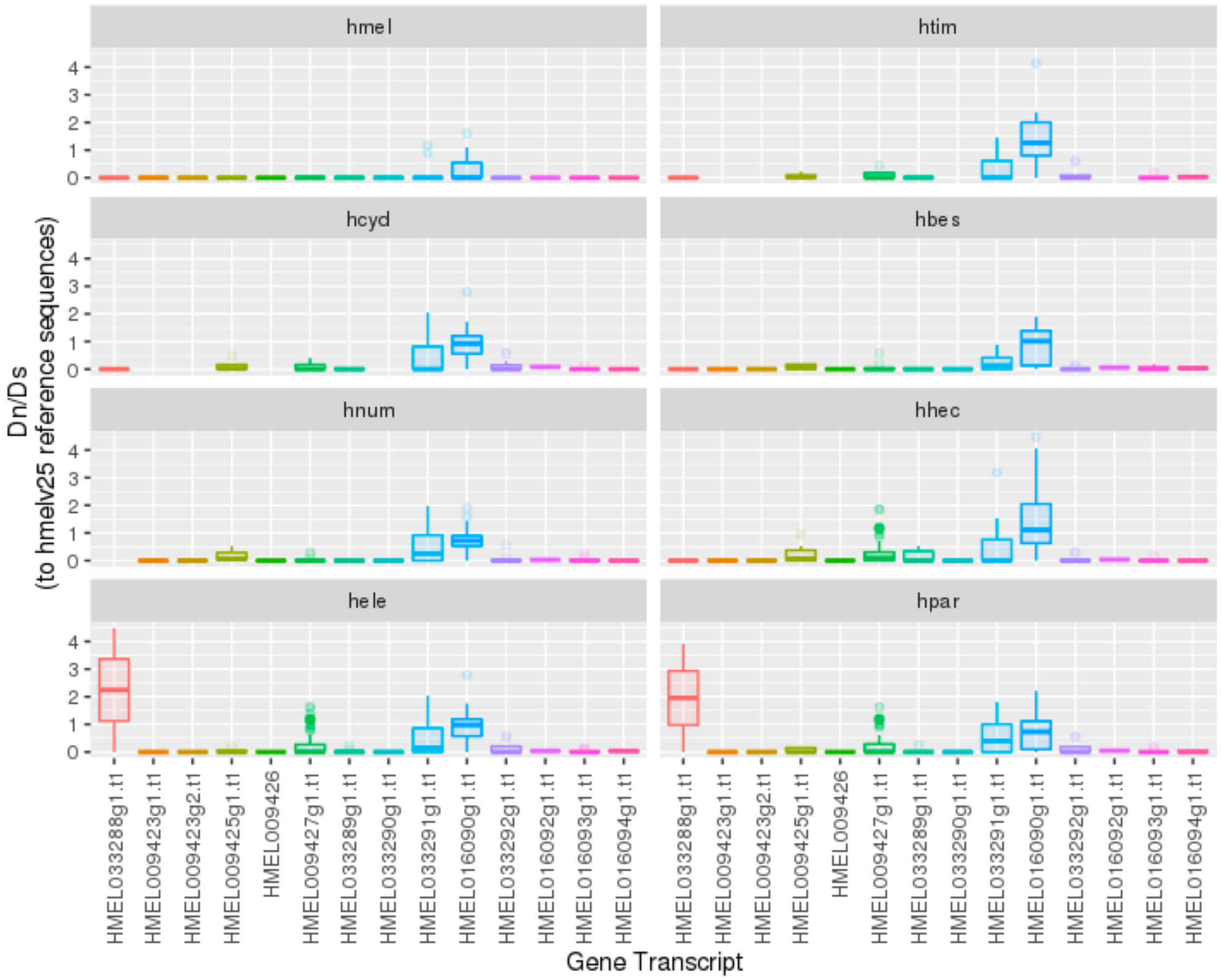


**Supplemental Fig. S55** - Dn/Ds estimates per exon in genes within the repeated regions on chromosome 2. Each boxplot shows the distribution of Dn/Ds estimates of all exon in each gene transcript. Dn/Ds was calculated between each exonic copy and the reference *H. melpomene* exonic sequence. Species codes: hmel - *H. melpomene*; htim - *H. timareta*; hcyd - *H. cydno*; hbes - *H. besckei*; hnum - *H. numata*; hhec - *H. hecale*; hele - *H. elevatus*; hpar - *H. pardalinus.*

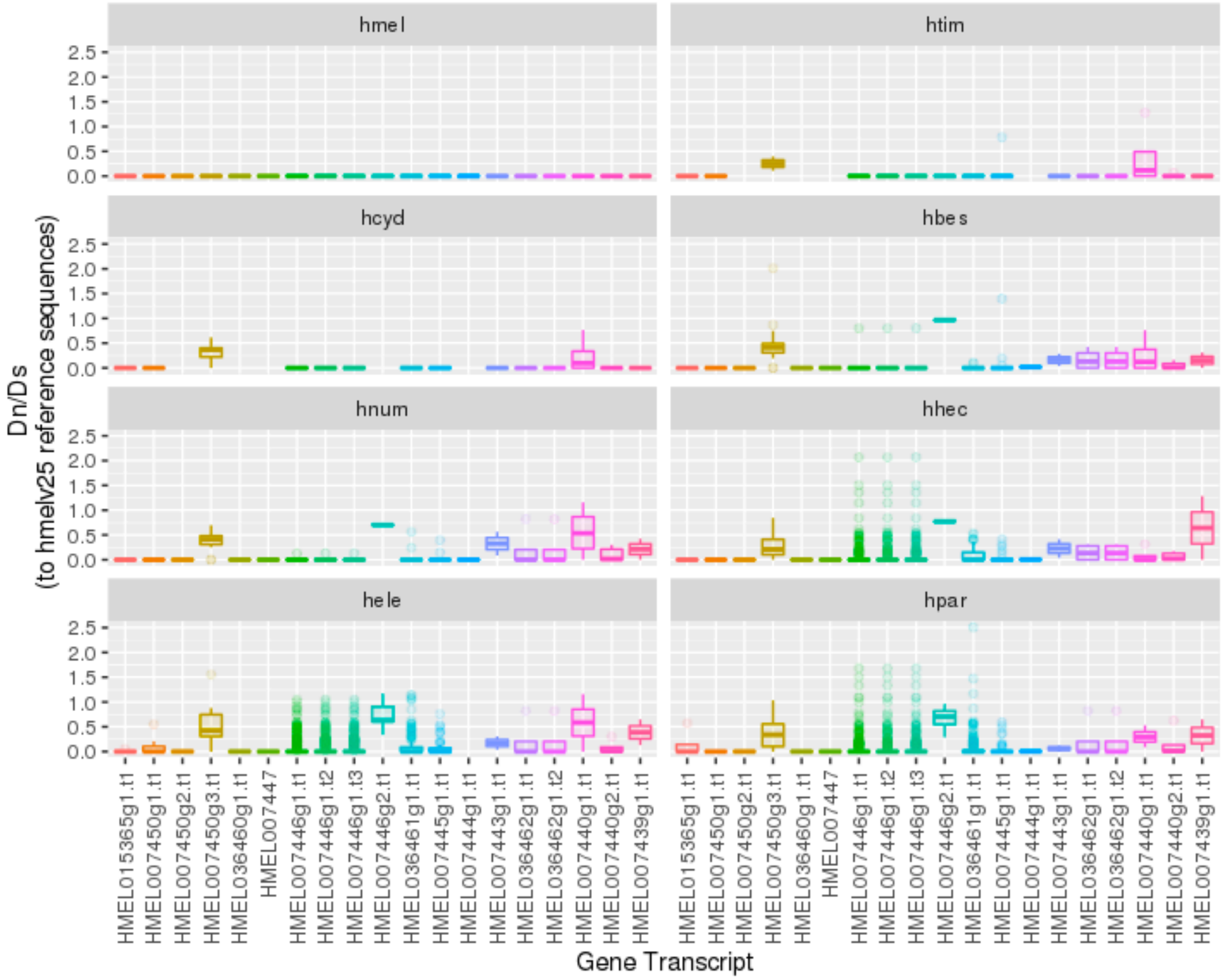


**Supplemental Fig. S56** - Dn/Ds estimates per exon in genes within the repeated regions on chromosome 4. Each boxplot shows the distribution of Dn/Ds estimates of all exon in each gene transcript. Dn/Ds was calculated between each exonic copy and the reference H. melpomene exonic sequence. Species codes: hmel - *H. melpomene*; htim - *H. timareta*; hcyd - *H. cydno*; hbes - *H. besckei*; hnum - *H. numata*; hhec - *H. hecale*; hele - *H. elevatus*; hpar - *H. pardalinus*..

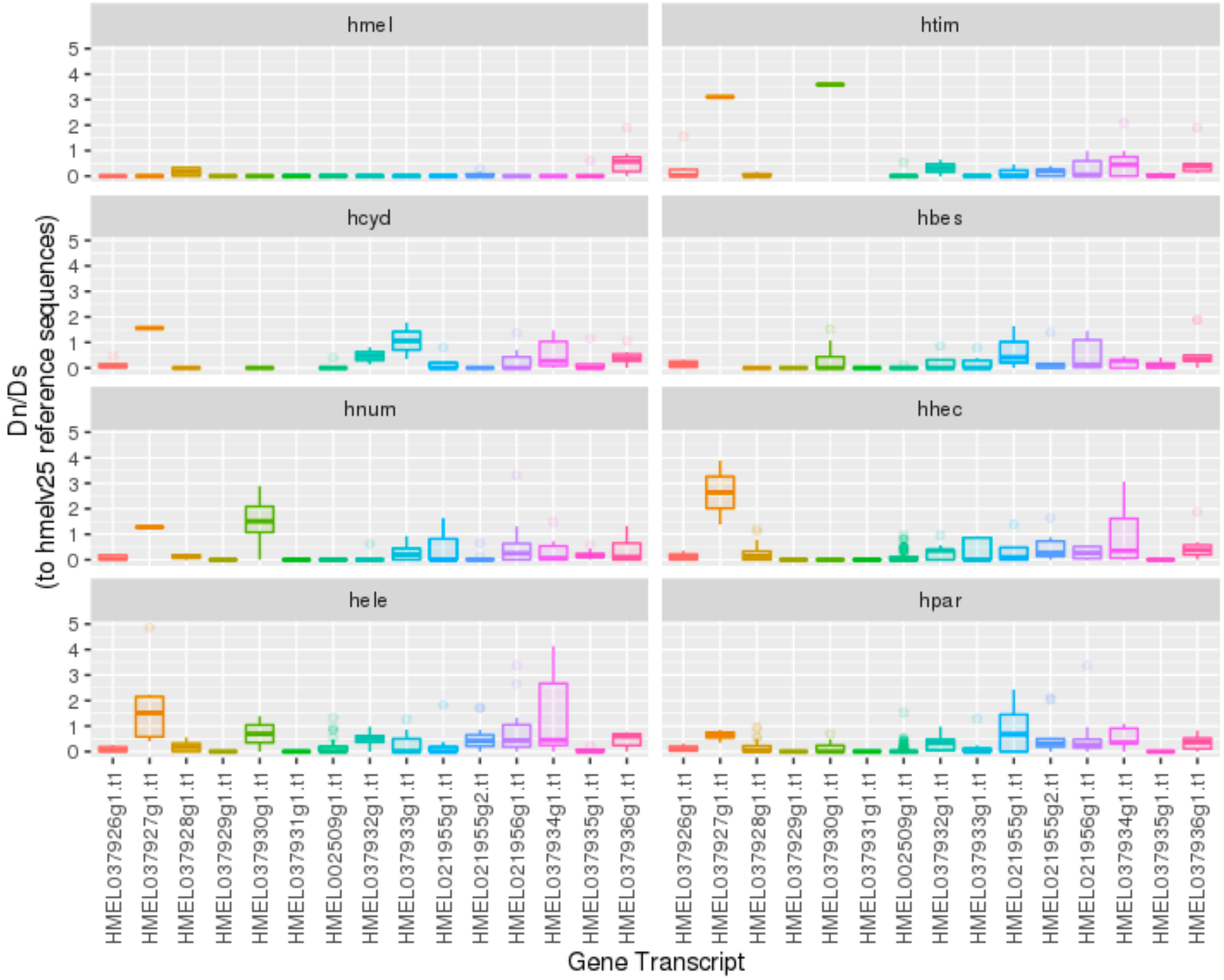


**Supplemental Fig. S57** - Dn/Ds estimates per exon in genes within the repeated regions on chromosome 8. Each boxplot shows the distribution of Dn/Ds estimates of all exon in each gene transcript. Dn/Ds was calculated between each exonic copy and the reference H. melpomene exonic sequence. Species codes: hmel - *H. melpomene*; htim - *H. timareta*; hcyd - *H. cydno*; hbes - *H. besckei*; hnum - *H. numata*; hhec - *H. hecale*; hele - *H. elevatus*; hpar - *H. pardalinus.*

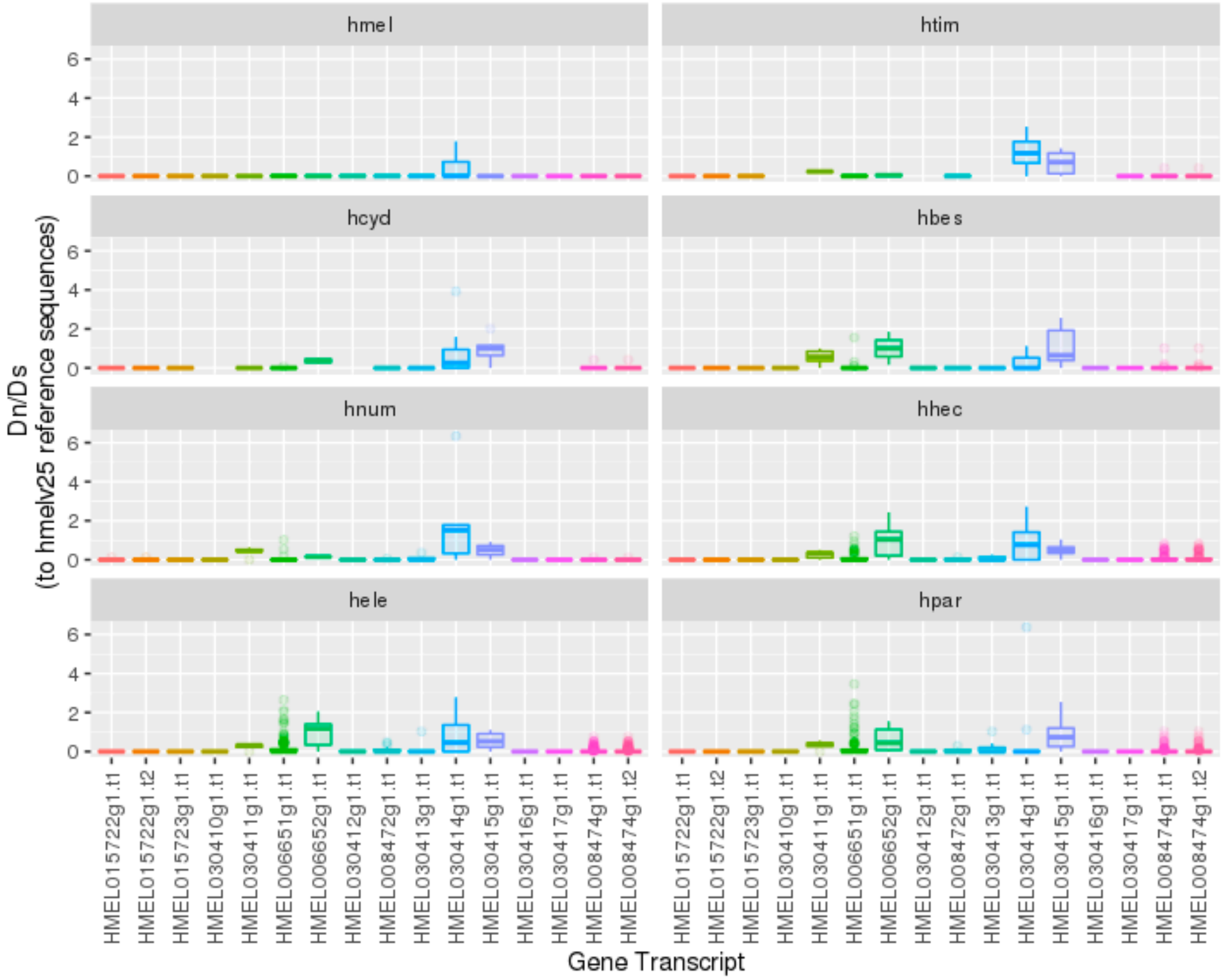


**Supplemental Fig. S58** - Dn/Ds estimates per exon in genes within the repeated regions on chromosome 9. Each boxplot shows the distribution of Dn/Ds estimates of all exon in each gene transcript. Dn/Ds was calculated between each exonic copy and the reference H. melpomene exonic sequence. Species codes: hmel - *H. melpomene*; htim - *H. timareta*; hcyd - *H. cydno*; hbes - *H. besckei*; hnum - *H. numata*; hhec - *H. hecale*; hele - *H. elevatus*; hpar - *H. pardalinus.*

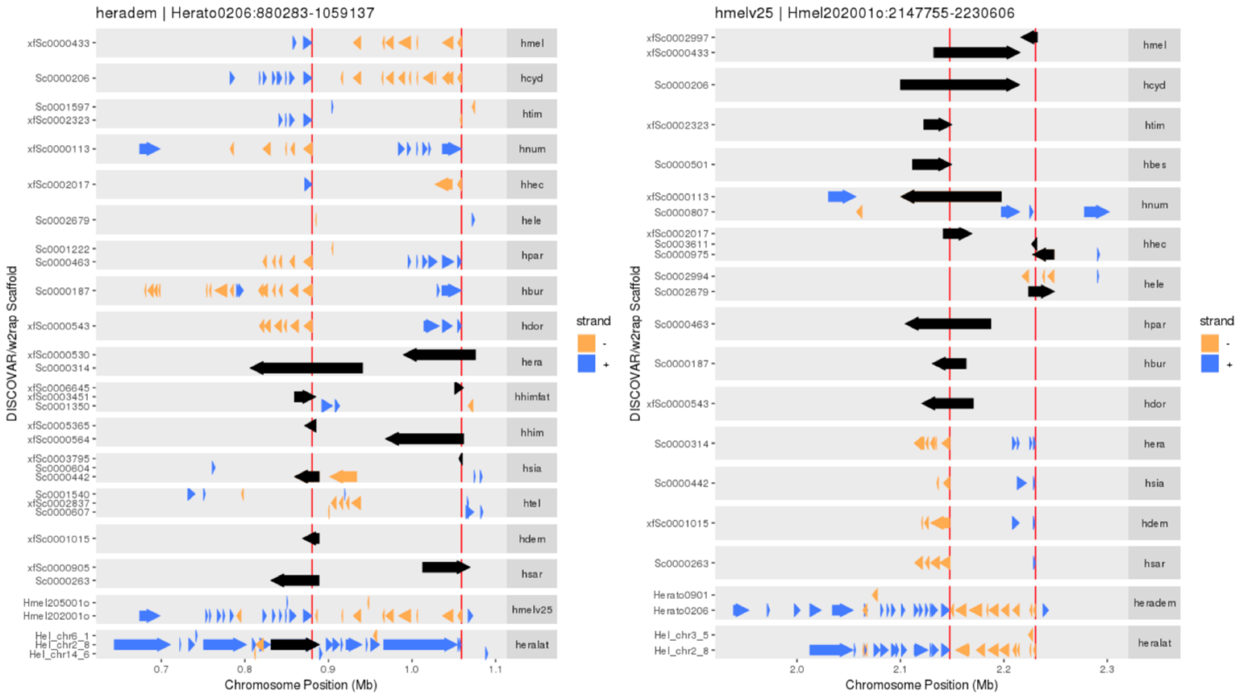


**Supplemental Fig. S59** – Inversion on chromosome 2. Alignments of scaffolds to the *H. erato demophoon* (left) and *H. melpomene* (right) reference genomes are represented by the arrows. The direction and colour of the arrows represent whether the alignments are to the forward strand (blue rightwards arrows) or the reverse strand (yellow leftwards arrows). Black arrows represent alignments spanning the inversion breakpoints (red vertical lines). Species codes are as in Figure 1.

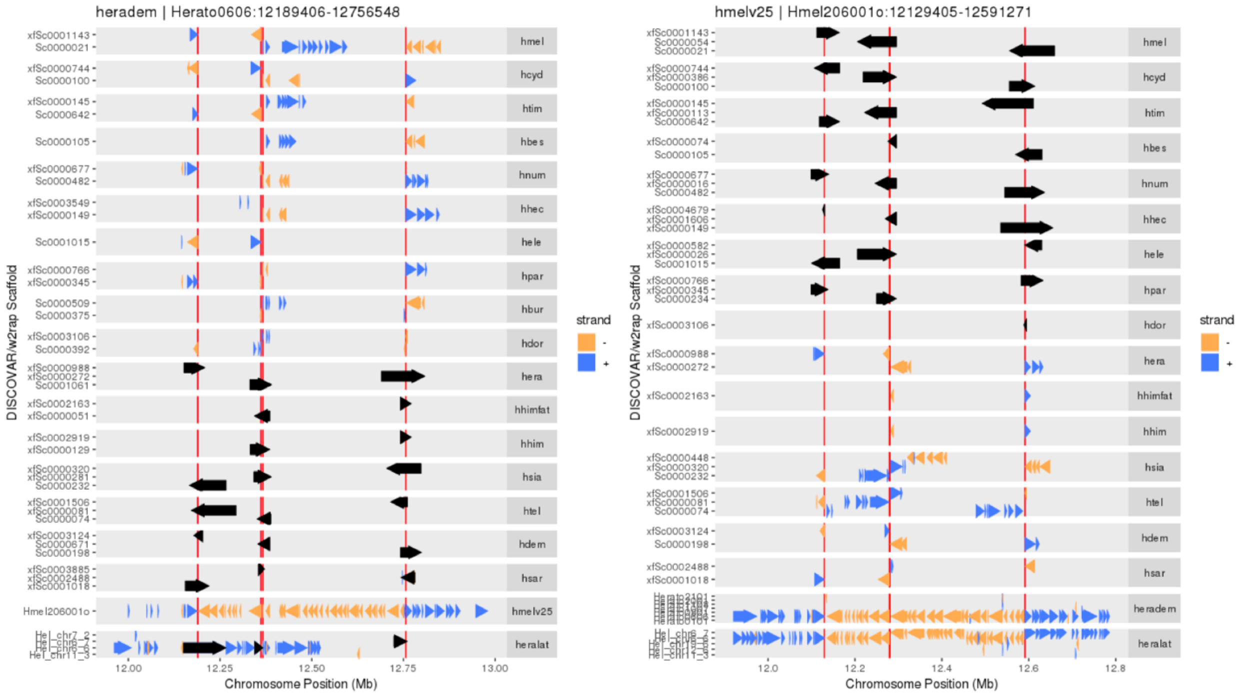


**Supplemental Fig. S60** – Tandem inversions on chromosome 6. Alignments of scaffolds to the *H. erato demophoon* (left) and *H. melpomene* (right) reference genomes are represented by the arrows. The direction and colour of the arrows represent whether the alignments are to the forward strand (blue rightwards arrows) or the reverse strand (yellow leftwards arrows). Black arrows represent alignments spanning the inversion breakpoints (red vertical lines). Species codes are as in Figure 1.

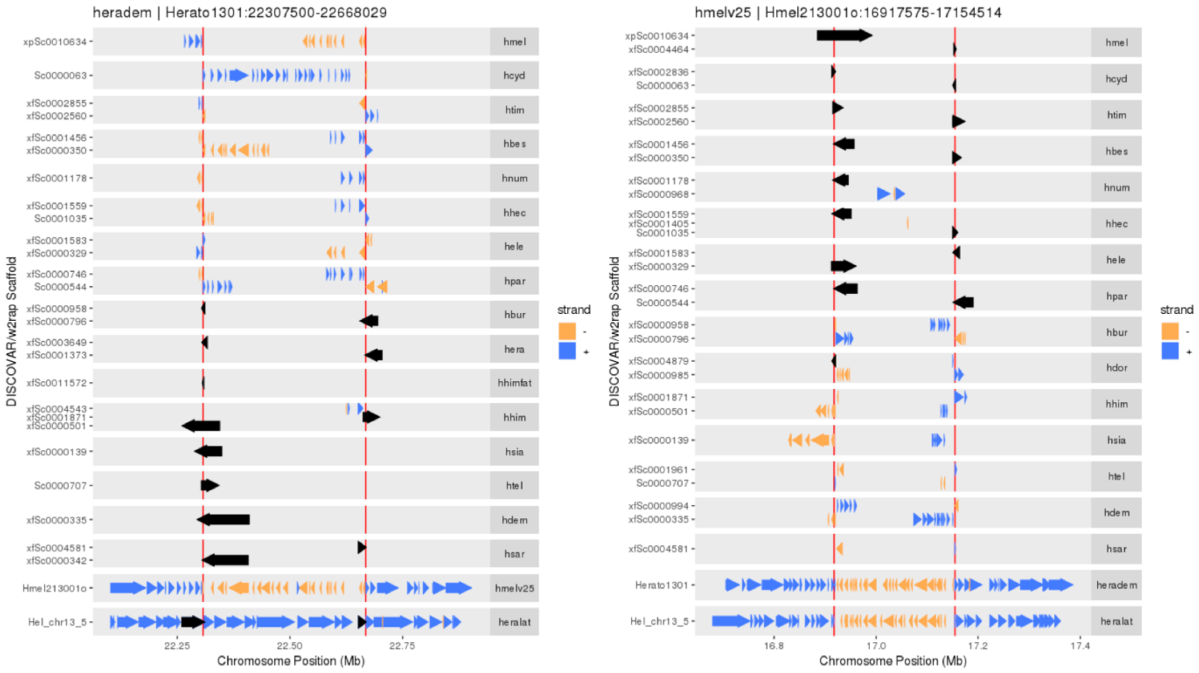


**Supplemental Fig. S61** - Inversion on chromosome 13. Alignments of scaffolds to the *H. erato demophoon* (left) and *H. melpomene* (right) reference genomes are represented by the arrows. The direction and colour of the arrows represent whether the alignments are to the forward strand (blue rightwards arrows) or the reverse strand (yellow leftwards arrows). Black arrows represent alignments spanning the inversion breakpoints (red vertical lines). Species codes are as in Figure 1.

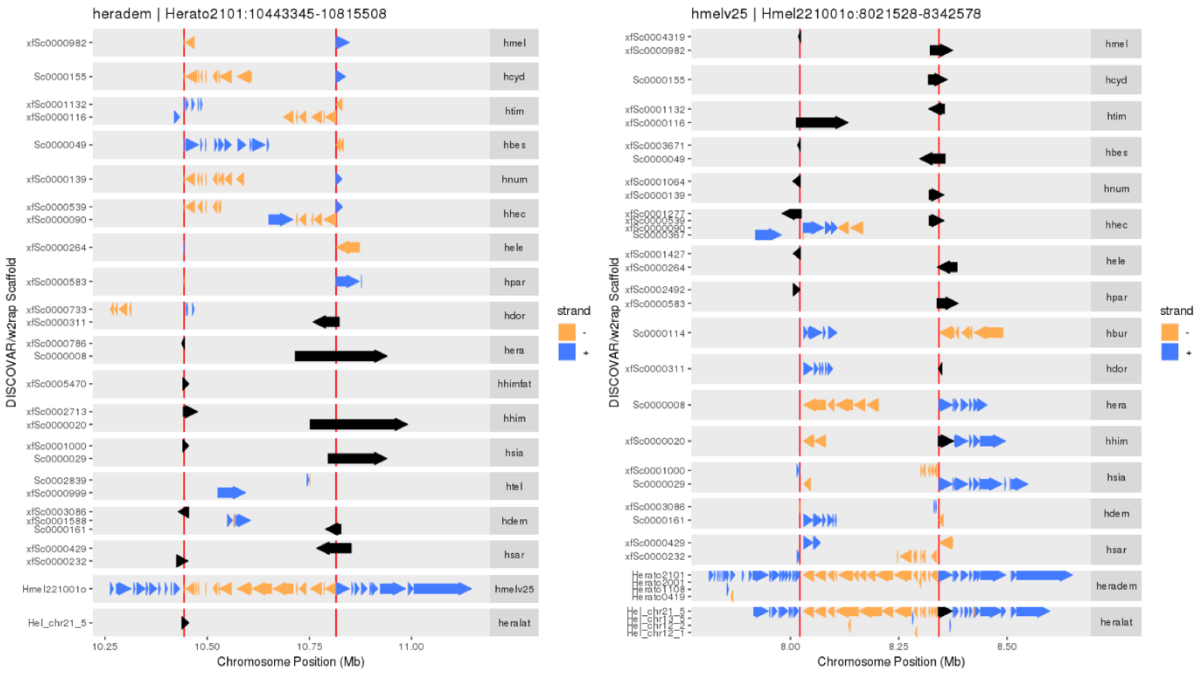


**Supplemental Fig. S62** - Inversion on the Z chromosome, i.e. chromosome21. Alignments of scaffolds to the *H. erato demophoon* (left) and *H. melpomene* (right) reference genomes are represented by the arrows. The direction and colour of the arrows represent whether the alignments are to the forward strand (blue rightwards arrows) or the reverse strand (yellow leftwards arrows). Black arrows represent alignments spanning the inversion breakpoints (red vertical lines). Species codes are as in Figure 1.

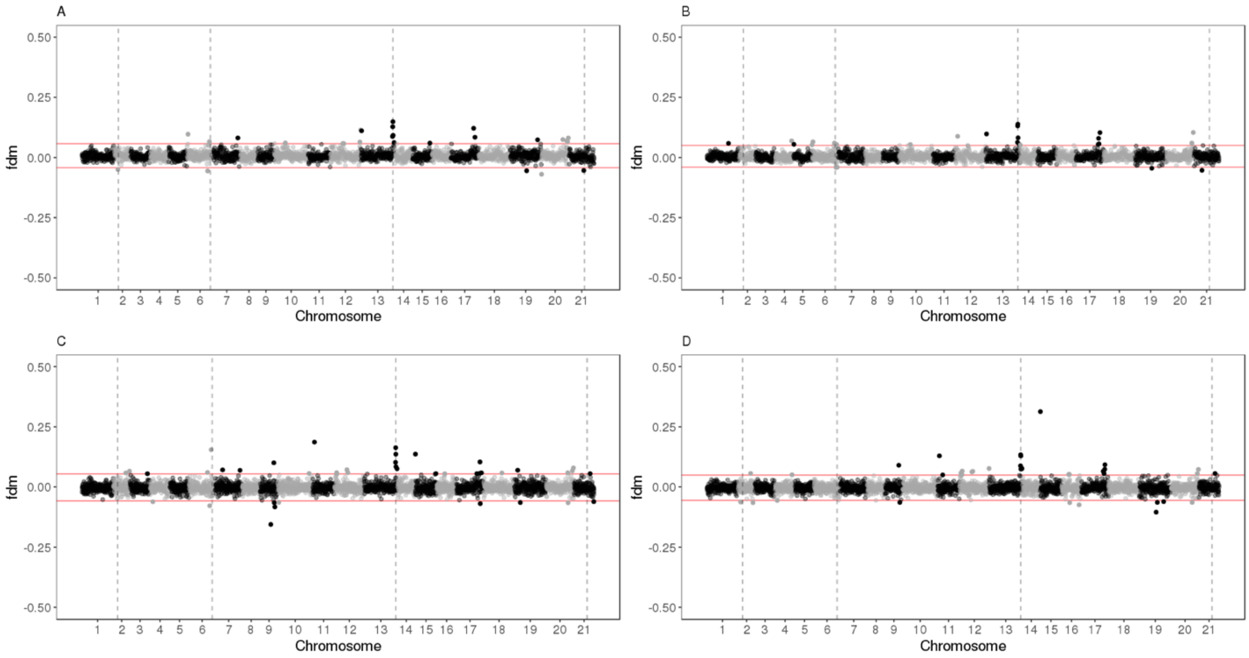


**Supplemental Fig. S63** – The $f_{dM}$ statistic along the genome. The $f_{dM}$ statistic was calculated in 25 kb non-overlapping windows across the genome, based on mapping of re-sequencing data to the *H. melpomene* (A,C) and the *H. erato demophoon* (B,D) reference genomes. In the top two panels (A,B) we test an excessive share of variation between *H. melpomene* and *H. erato demophoon* with *H. burneyi*, and in the bottom panels (C,D) with *H. doris*. *H. melpomene* and H. burneyi (or *H. doris*) were considered to be the ingroup species and H. erato the outgroup. Derived alleles were determined using *E. tales*. Chromosomes are shown with alternating grey and black colours. The location of inversions is given by the dashed vertical lines.

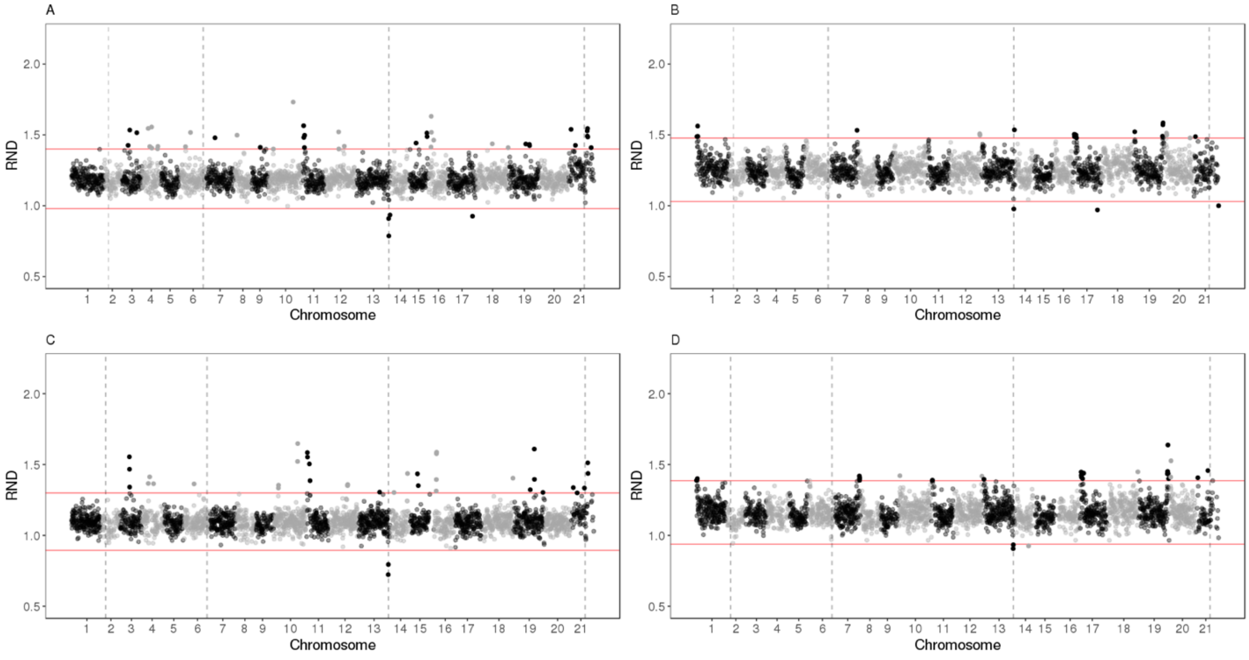


**Supplemental Fig. S64** – Divergence of H. erato to either *H. burneyi* (A,B) or *H. doris* (C,D), normalized by the *H. erato - H. melpomene* divergence. The relative divergence was calculated in 25 kb non-overlapping windows across the genome, based on mapping of re-sequencing data to the H. melpomene (A,C) and the *H. erato* (B,D) reference genomes. Chromosomes are shown with alternating grey and black colours. The location of inversions is given by the dashed vertical lines.

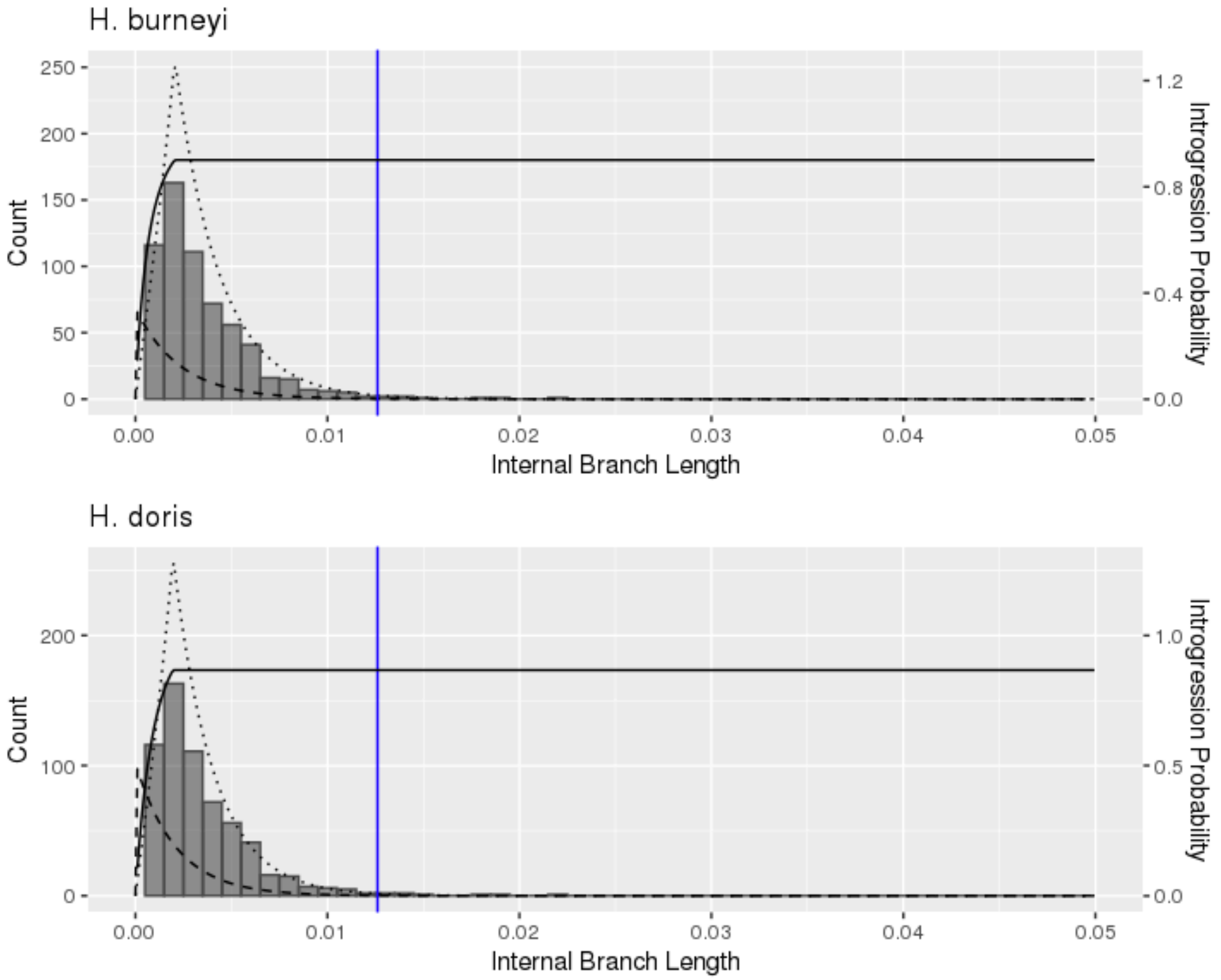


**Supplemental Fig. S65** – Histogram of internal branch lengths in windows with the topology (*H. melpomene*,(*H. erato*,*H. burneyi*)) – top panel – and (*H. melpomene*,(*H. erato*,*H. doris*)) – bottom panel. The inferred ILS (dashed line) and introgression (dotted line) distributions as were inferred by QuIBL under the ILS + Introgression model. The average internal branch length in the chromosome 13 inversion is shown as a blue vertical line. The probability of introgression as a function of internal branch length is shown as a full black line.

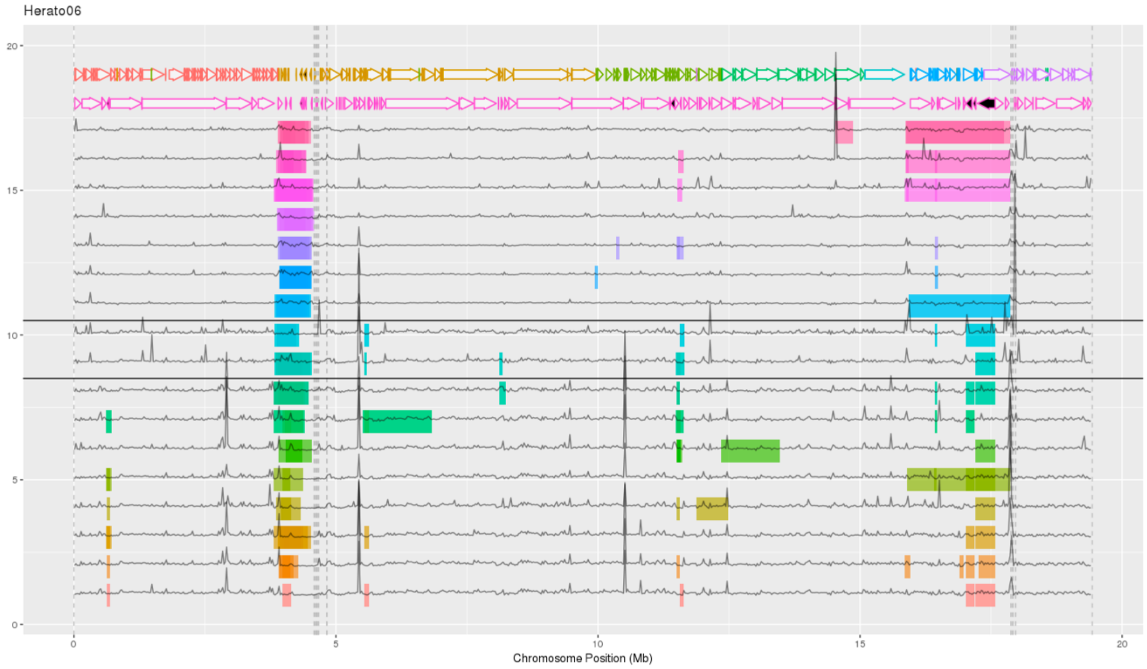


**Supplemental Fig. S66** – Candidate inversions on chromosome 6. Candidate inversions are depicted by the coloured rectangles in each species (each line represents one species) and the grey lines represent relative coverage along the chromosome. The top two lines show the alignment (represented by the arrows) of *H. erato lativitta* (top line) and *H. melpomene* (second line from the top) reference genomes scaffolds to the *H. erato demophoon* reference genome. Each scaffold is depicted using a unique colour and black filled arrows represent inverted alignments. The vertical dashed lines represent scaffold boundaries within chromosome 6 of the *H. erato demophoon* reference genome. Note for example the repetitive region near 4-5 Mb (common to all species), where there are several candidate inversions (within and across species). These likely represent artifacts resulting from poor alignments, as suggested by the fragmented and/or lack of alignments of scaffolds of the two reference genomes (*H. erato lativitta* and *H. melpomene*) to the *H. erato demophoon* reference genome in this region. The list of species from bottom to top is: hmel, htim, hcyd, hbes, hnum, hhec, hele, hpar, hbur, hdor, hera, hhimfat, hhim, hsia, htel, hdem, hsar. Species codes are as in Figure 1.

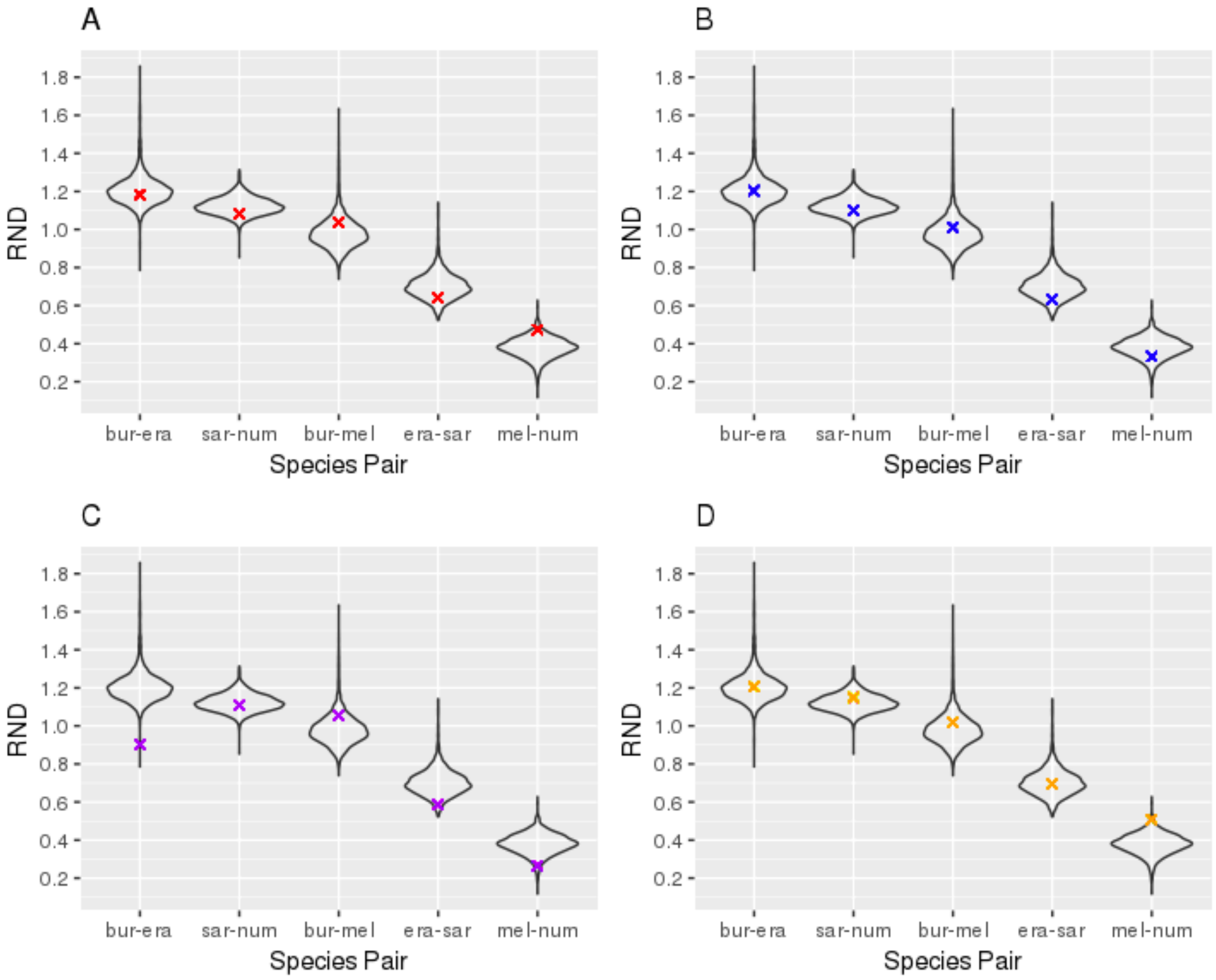


**Supplemental Fig. S67** – Relative divergence between species pairs, based on mapping of re-sequencing data to the *H. melpomene* reference genome. The relative divergence was calculated in non-overlapping 25 kb windows, by estimating the genetic distance between each population pair divided by the genetic distance between *H. melpomene* and *H. erato*. The coloured crosses depict relative divergence estimates within the inversion regions on chromosome 2 (A), chromosome 6 (B), chromosome 13 (C) and chromosome 21 (D). Species codes are as in Figure 1.

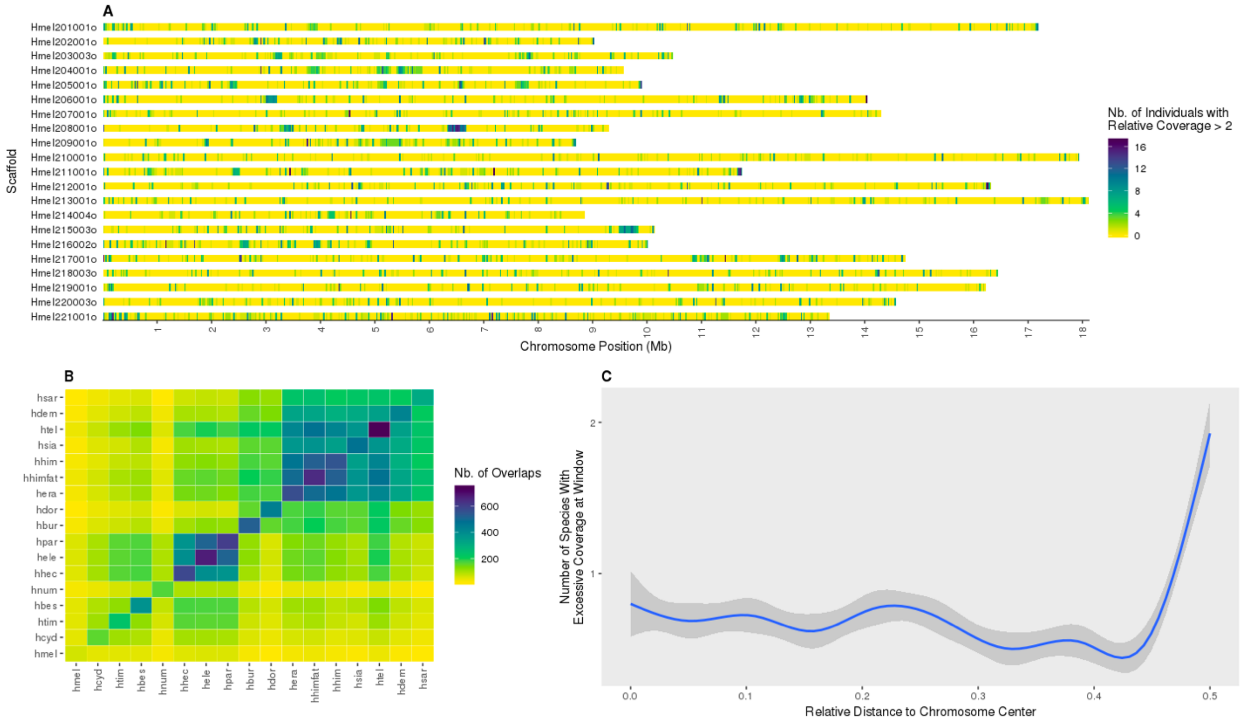


**Supplemental Fig. S68** – Summary of repeat regions across species. In each species, we consider repeat regions those windows with a relative coverage greater than 2. Relative coverage was calculated in 25 kb non-overlapping sliding windows along each of the largest scaffolds in each chromosome. **(A)** Distribution and frequency of repeat regions along chromosomes. While 79% of the windows are free of repeat regions across all species, 8% contains a repetitive region present in a single species while the remaining 13% harbour repeat regions present in at least 2 species. **(B)** Number of repeat regions shared between species in each species pair. Species in the *erato*/*sara* clade and in the *hecale*/*elevatus*/*pardalinus* trio, show a higher number of repeat regions. **(C)** Relation between the frequency of repeat regions (across species) and chromosome position. Repeat regions shared across species are more common at chromosome ends. Together these results suggest that repeat regions are not uniformly distributed across the genome and tend to be common between species. Species codes are as in Figure 1.

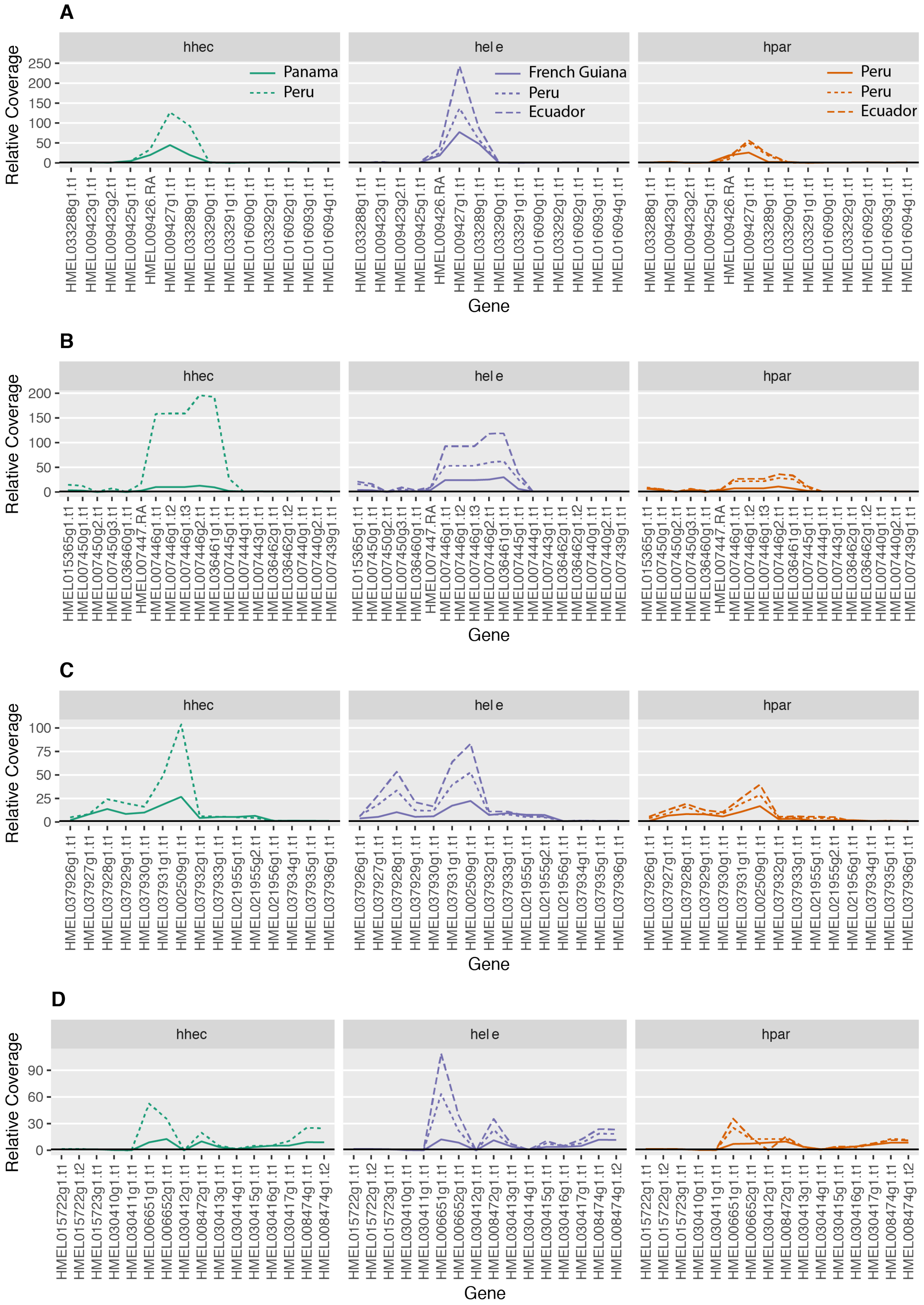


**Supplemental Fig. S69** - Gene copy number (CN) estimates for the repeat regions in (A) chromosome 2, (B) chromosome 4, (C) chromosome 8 and (D) chromosome 9. The normalized read coverage of reads mapped to the *H. melpomene* reference was used as a proxy of CN. Gene CN was estimated both for extra-Amazonian (full lines) and Amazonian (dashed lines) populations of *H. hecale* (hhec), *H. el*evatus (hele) and *H. pardalinus* (hpar). Details about the populations sampled can be found in Supplemental Table S11.
